# End-to-end reconstruction for cryo electron ptychographic single-particle analysis

**DOI:** 10.64898/2026.10.31.763295

**Authors:** S. You, N. Palatkin, S. Shao, P. M. Pelz

**Author notes:** These authors contributed equally (S. You, N. Palatkin, S. Shao).

## Abstract

Cryo-electron ptychographic (cryo-ptycho) single-particle analysis (SPA) is a diffractive imaging method for protein structure determination. Current reconstruction workflows sequentially reconstruct a 2D phase micrograph from each scan, and then apply conventional SPA software to the resulting projections to obtain a 3D potential map. Here, we introduce End2EndPtychoSPA, an end-to-end reconstruction algorithm that directly fits a 3D electrostatic potential volume and particle poses to 4D-STEM diffraction data using a differentiable multislice forward model. On simulated haemoglobin data where the sequential route recovers initial poses, End2EndPtychoSPA with a learnable vitreous-ice layer reaches 3.22 Å resolution against the ground truth (FSC 0.5), compared with 3.41 Å for the sequential route. We also compare End2EndPtychoSPA with the sequential route on four published experimental cryo-ptycho SPA datasets. On apoferritin (11,552 particles), the End2EndPtychoSPA map improves agreement with an external atomic model over both our matched sequential route on the same particles and the deposited EMD-19425 half-map mean, and shifts the unmasked half-map FSC 0.143 crossing from 6.83 Å to 5.99 Å. On T20S, TMV and Phi92, End2EndPtychoSPA does not yet improve upon the 2D route.

**Synopsis:** End2EndPtychoSPA refines a 3D electrostatic potential and the particle poses directly against cryo-4D-STEM diffraction data with a differentiable multislice forward model. Starting from a 2D ptychographic reconstruction, it improves the reconstructed volume on simulated haemoglobin and experimental apoferritin, while pre-registered tests on three further specimens show where it does not yet improve upon the two-step approach.

## 1. Introduction

Electron ptychography reconstructs the complex transmission function of a specimen from a set of two-dimensional diffraction patterns recorded at each position of a scanned coherent electron probe (Hoppe, 1969; Rodenburg & Faulkner, 2004; Maiden & Rodenburg, 2009; Rodenburg, 2008; Ophus, 2019; Jiang *et* al., 2018). Because ptychography provides quantitative phase contrast with a wide spatial-frequency transfer band at low electron dose, it has been applied to radiation-sensitive biological specimens (Pelz et al., 2017; Zhou et al., 2020; Huang et al., 2023; Li & Li, 2026), alongside other four-dimensional scanning transmission electron microscopy (4D-STEM) methods such as tilt-corrected bright-field STEM (Yu et al., 2025) and 4D-STEM cryotomography (Seifer et al., 2024). Pei et al. (2023) reported the first SPA reconstructions from cryo-ptycho data. Subsequently, Küçükoğlu *et al*. (2024*d*) reconstructed apoferritin, the Phi92 bacteriophage tail sheath and tobacco mosaic virus (TMV) to sub-nanometre resolution from low-dose 4D-STEM scans and deposited both the raw diffraction datasets and the reconstructed maps. Li, Li et al. (2026) then demonstrated that a mismatch between real-space scan step and reciprocal-space detector sampling limits such reconstructions and corrected this sampling condition for the 20S proteasome (T20S), while more recent work has extended cryo-ptycho SPA to higher resolution using direct electron counting (Li, Shen et al., 2026).

However, all of these studies follow a two-step reconstruction workflow, which we refer to as the ‘2D route’. In this approach, each 4D-STEM scan is first inverted independently into a two-dimensional phase micrograph, after which individual particles are picked and processed using standard single-particle software developed for conventional plane-wave cryo-EM (Scheres, 2012; Punjani et al., 2017). Each scan is reconstructed as a single 2D object, although the specimen has finite thickness and may contain particles that overlap along the beam direction. Regularisation and errors in the probe and scan positions can also affect the resulting particle images, which SPA then treats as projections of a common 3D structure. These effects can limit the subsequent 3D reconstruction. Furthermore, dynamical scattering simulations of TMV embedded in vitreous ice at 300 kV indicate that single-slice models lose information at resolutions finer than about 2.5 Å. Modelling the specimen as several slices, each thinner than 6 nm, is needed to describe electron propagation accurately at these higher spatial frequencies. (Leidl *et al*., 2023).

To avoid intermediate two-dimensional reconstructions, the three-dimensional electrostatic potential can be reconstructed directly from the measured diffraction patterns, as demonstrated in electron and X-ray tomography for a single specimen imaged across a tilt series (Broek & Koch, 2012; 2013; Gilles *et al*., 2018; Kandel *et al*., 2019; Chang *et al*., 2020; Du *et al*., 2021; Ding *et al*., 2022; Lee *et al*., 2023; Pelz *et al*., 2023; Romanov *et al*., 2025; You *et al*., 2025). Related methods for plane-wave electron microscopy also account for the specimen’s extent along the beam direction, which is neglected when the specimen is represented as a single projected layer. Ewald-sphere curvature corrections account for propagation geometry within a single-scattering framework (DeRosier, 2000; Wolf *et al*., 2006; Russo & Henderson, 2018). Conjugated Holographic Reconstruction (Gureyev *et al*., 2022) back-propagates each defocused micro-graph with a conjugated phase factor to correct Ewald-sphere curvature and could replace standard contrast-transfer-function (CTF) correction in RELION or FREALIGN, while Ghostbuster (Yeo *et al*., 2024) retrieves the exit-wave phase and models intraparticle scattering within the first Born approximation; both approaches have been demonstrated in numerical simulations. Multislice forward models have also been combined with plane-wave tilt series to account for multiple scattering: Ren *et al*. (2020) reconstructed three-dimensional potentials from simulated phase-contrast tilt series, and PhaseT3M incorporated Bayesian optimization of tilt-series alignment and lens aberrations to reach 1.6 Å resolution on a Co_3_O_4_ nanoparticle by electron tomography (Lee *et al*., 2026).

In single-particle analysis, however, thousands of identical macromolecules are embedded in vitreous ice in unknown orientations rather than being imaged as a single tilted object. In X-ray free-electron laser diffraction, the analogous problem of reconstructing a three-dimensional object from two-dimensional diffraction patterns of randomly oriented particles is known as cryptotomography (Loh et al., 2010). The expand–maximize–compress algorithm of Loh & Elser (2009) reconstructs a 3D diffraction-intensity distribution from noisy 2D patterns, accounting for their unknown orientations through a Poisson likelihood model. A separate phase-retrieval step then recovers the particle’s real-space structure. Because EMC assumes plane-wave illumination of an isolated particle, each diffraction pattern depends only on the particle orientation and is invariant to lateral translation. Differentiable forward models have also been developed for conventional plane-wave cryo-EM micrographs (Ullrich et al., 2019; Zhong et al., 2021; Levy et al., 2022), where particle poses and a 3D volume are estimated from 2D images rather than from 4D-STEM diffraction patterns.

In this work, we introduce End2EndPtychoSPA, an end-to-end multislice reconstruction algorithm that jointly reconstructs a 3D macromolecular volume and per-particle poses directly from cryo-4D-STEM diffraction data. We test the algorithm first on simulated datasets with known ground truth, and then compare it with our own 2D reconstructions and published maps on experimental datasets. As illustrated in Figure 1, our forward model places rotated and translated copies of a shared three-dimensional electrostatic potential volume into each local scan window, propagates the focused electron probe through the composite potential using the multislice method (Cowley & Moodie, 1957; Chen et al., 2021), and compares the predicted farfield diffraction intensities with the measured 4D-STEM counts through a Poisson negative log-likelihood. Using automatic differentiation, we jointly refine the shared 3D potential volume, the particle orientations and positions, and a per-window defocus offset directly against the diffraction data.

**Figure 1.**
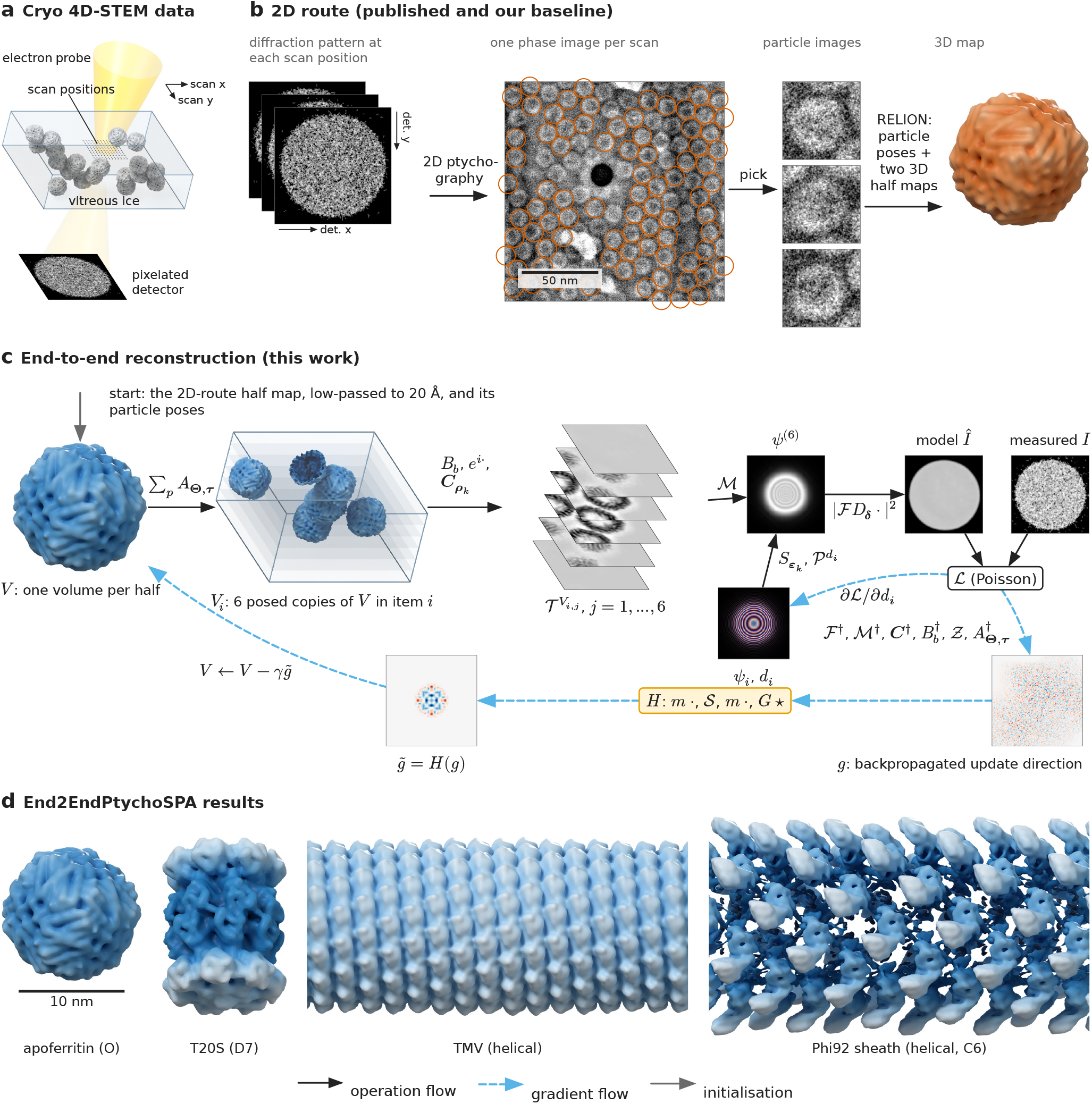
Comparison of the two-step 2D reconstruction route and direct end-to-end refinement from cryo 4D-STEM data. (a) A focused electron probe scans the vitrified specimen. A pixelated detector records one two-dimensional diffraction pattern at each scan position. The schematic shows 19 of the 39 picked particles in one scan, the vitreous ice layer (blue box, not to scale) and one 12 × 12 window of scan positions (dots); the arrows give the scan directions *x* and *y*. (b) In the published 2D route, ptychography reconstructs one 2D phase image from each scan. Particles are picked in the phase images (circles), and RELION refines the particle poses and two independent 3D half maps from the particle images (half-map mean shown). The arrows on the diffraction patterns give the detector axes *x* and *y*. (c) In End2EndPtychoSPA, each gold-standard half set is used to directly reconstruct one shared 3D potential volume *V* from the raw diffraction patterns, initialized from the 2D-route half map and particle poses. The illustrated scan window contains 6 posed copies of *V*. Multislice propagation through 6 slices predicts the far-field diffraction intensities, which are compared with the measured patterns through a Poisson negative log-likelihood (NLL; Section 2.3). Backpropagation yields the update direction *g* (Equation 13). The preconditioner and filter cascade *H* (Figure 3) transforms *g* into the preconditioned update 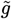 applied to *V*. Particle poses and per-window defocus offsets are refined simultaneously while the base probe remains fixed. Solid black arrows denote the forward model; dashed blue arrows indicate backpropagation and parameter updates. The tensor images in (c) are real tensors of this item, shown as qualitative illustrations with their own display scales: model and measured patterns as log_10_ counts on one common scale, the probe with amplitude as brightness and phase as hue, and the gradients on a signed scale that is symmetric for each image. (d) Final End2EndPtychoSPA reconstructions shown at a common scale (half-map means coloured by radial distance from the particle centre or helical axis, and contoured at the atomic-model volume; Phi92 has no atomic model of its whole tail tube and is contoured at 1.5 standard deviations inside its cylinder); helical reconstructions are truncated to a length of 360 Å.

Our contribution beyond the state of the art is fourfold:

1. We adapt and implement an end-to-end ptychographic tomography framework for cryo-ptychographic SPA that jointly refines a shared 3D electrostatic potential volume, per-particle poses and per-window defocus offsets from 4D-STEM diffraction patterns while accounting for overlapping neighbour particles within each scan window.
2. We test the convergence of the algorithm using ground-truth multislice simulations. On myoglobin (17.9 kDa; Section S9), local gradient-based refinement does not recover particle orientations that are lost during the initial 2D ptychographic step. On haemoglobin (64.5 kDa; Section 3.1), where the 2D route provides coarse initial orientations, End2EndPtychoSPA combined with a learnable vitreous-ice background model refines the poses and improves the 3D reconstruction relative to the ground truth.
3. We compare End2EndPtychoSPA on four published experimental cryo-4D-STEM datasets against our own 2D route on the same raw diffraction data, half-set assignments and symmetry constraints. The particle sets are identical for apoferritin and TMV; for Phi92 the two routes share 1,648 of 1,651 segments (Section 2.1). We also compare with the deposited EMDB maps as an external reference. Their particle selection and post-processing differ from ours; only for apoferritin is the published particle count known, and we match it.
4. We apply the low-pass filter last in the volume gradient hook. A spatial support mask applied after the filter would move the strongly correlated low-frequency update into high spatial frequencies, couple the two gold-standard half maps there and inflate the half-map Fourier shell correlation (FSC) (Scheres & Chen, 2012; Henderson *et al*., 2012). We check every map with an empirical diagnostic based on high-frequency phase randomization (Chen *et al*., 2013) that detects a high-frequency FSC rebound.

## 2. Methods

### 2.1. Datasets and experimental comparison protocol

We tested End2EndPtychoSPA on four publicly available cryo-4D-STEM datasets (Table 1), each with a deposited 3D map from the 2D route. The apoferritin (EMPIAR-12236 (Küçükoğlu *et al*., 2024*a*)), tobacco mosaic virus (TMV; EMPIAR-12234 (Küçükoğlu *et al*., 2024*b*)) and Phi92 bacteriophage tail-sheath (EMPIAR-12233 (Küçükoğlu *et al*., 2024*c*)) datasets were acquired by Küçükoğlu *et al*. (2024*d*), and the 20S proteasome dataset (T20S; EMPIAR-13137 (Li *et al*., 2025*a*), session 2) was recorded by Li, Li *et al*. (2026). Section 2.6 describes our ground-truth multislice simulations and their reconstruction workflows.

**Table 1.**
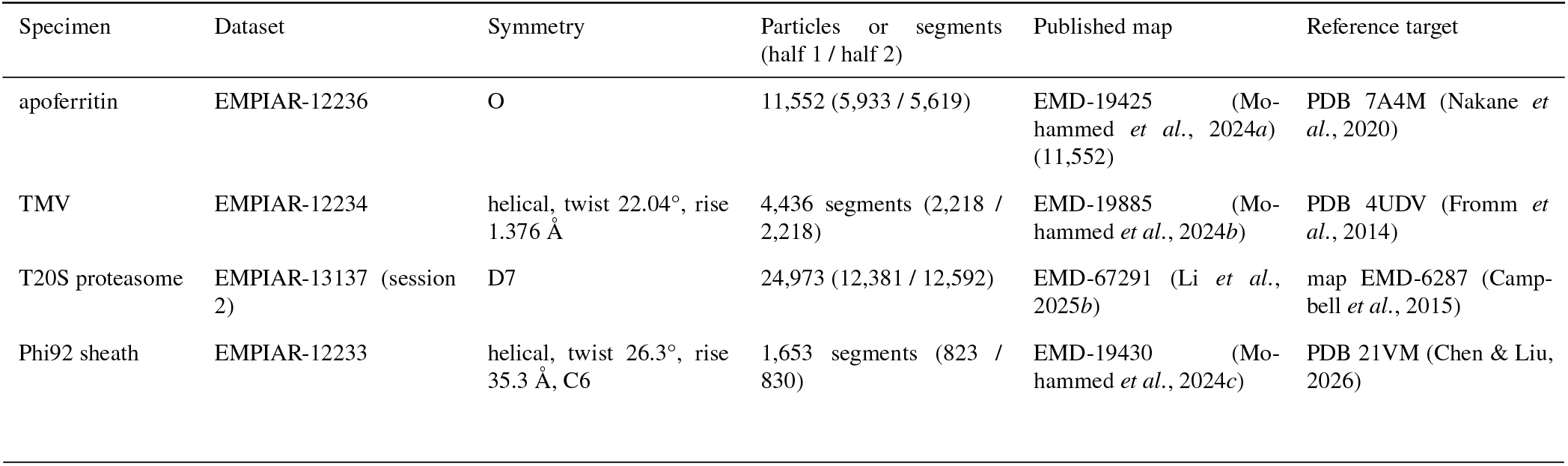
Experimental cryo 4D-STEM datasets analysed in this work. Particle and helical-segment counts are taken from the End2EndPtychoSPA input tables and reported as total (half 1 /half 2); the number of reconstructed items can be smaller (Section 2.3.3). For apoferritin, we match the published particle count of 11,552; for TMV, T20S and Phi92, the particle counts used in the deposited maps are not recorded in the public depositions. Reference atomic models and maps are used only for evaluation after reconstruction and do not enter as an initial model, alignment reference or support mask.

| Specimen | Dataset | Symmetry | Particles or segments (half 1 / half 2) | Published map | Reference target |
|---|---|---|---|---|---|
| apoferritin | EMPIAR-12236 | O | 11,552 (5,933 / 5,619) | EMD-19425 (Mo-hammed *et al.*, 2024*a*) (11,552) | PDB 7A4M (Nakane *et al.*, 2020) |
| TMV | EMPIAR-12234 | helical, twist 22.04°, rise 1.376 Å | 4,436 segments (2,218 / 2,218) | EMD-19885 (Mo-hammed *et al.*, 2024*b*) | PDB 4UDV (Fromm *et al.*, 2014) |
| T20S proteasome | EMPIAR-13137 (session 2) | D7 | 24,973 (12,381 / 12,592) | EMD-67291 (Li *et al.*, 2025*b*) | map EMD-6287 (Camp-bell *et al.*, 2015) |
| Phi92 sheath | EMPIAR-12233 | helical, twist 26.3°, rise 35.3 Å, C6 | 1,653 segments (823 / 830) | EMD-19430 (Mo-hammed *et al.*, 2024*c*) | PDB 21VM (Chen & Liu, 2026) |

End2EndPtychoSPA and our 2D-route baseline use the same raw diffraction data, point-group or helical symmetry constraints and gold-standard half-set assignments for each specimen, and the same particle or helical-segment coordinates except where stated below. On apoferritin, we match the published particle count of 11,552 (Küçükoğlu *et al*., 2024*d*), although the original particle coordinates were not deposited. On TMV, End2EndPtychoSPA uses all 4,436 helical segments from our 2D-route refinement. On Phi92, the input table contains every second picked segment (1,653 segments in total, of which 1,651 load into End2EndPtychoSPA because two scan windows cross the scan boundary), and our 2D-route baseline is refined on 1,648 of those segments. On T20S, the particle coordinates used for the deposited map are not publicly available, so a particle-matched comparison with the published reconstruction (Li, Li *et al*., 2026) is not possible.

### 2.2. Baseline 2D reconstruction route using RELION and cryoSPARC

In our 2D reconstruction route, we first reconstruct one 2D phase micrograph per 4D-STEM scan and then process the extracted particle images using standard single-particle workflows. For apoferritin, we first analysed each scan with single-sideband (SSB) ptychography (Rodenburg et al., 1993; Pennycook et al., 2015; You et al., 2026) to determine the scan-to-detector rotation angle, the low-order lens aberrations and an empirical detector noise gain. Using these initial calibration parameters, we then performed an iterative single-slice 2D ptychographic reconstruction for each scan while jointly refining all 12 coherent probe aberration coefficients up to fourth order. Particle coordinates were assembled from two sources: on 60 scans, we transferred 11,971 particle picks and initial poses from a reproduction of the published cryoSPARC workflow (Punjani et al., 2017), and on 29 additional scans we picked 9,708 candidate particles using a circular disk template in RELION (Scheres, 2012), retaining 7,093 particles after 3D classification. This combined pool of 19,064 particles across 89 scans was refined with RELION 3D auto-refine under octahedral (O) symmetry without CTF correction and without a solvent mask, using scan-pure gold-standard half sets defined in advance by alternating the sorted scan indices. Finally, to match the published dataset size of 11,552 particles, we ranked the refined pool by RELION maximum-likelihood posterior probability, selected the top 11,552 particles while preserving the pool’s half-set ratio (5,933 in half 1 and 5,619 in half 2), and re-refined this subset under identical RELION settings (designated run R1).

Because the half sets are split by scan, no 4D-STEM scan contributes particles to both R1 halves. The 11,552 selected R1 particles come from 85 of the 89 scans and reach an unmasked half-map FSC of 0.143 at 6.83 Å resolution. We use the R1 half maps, refined poses and particle table both as our 2D-route baseline and as the initialization for End2EndPtychoSPA (Section 2.4).

For the remaining three experimental datasets, our 2D-route baselines consist of a cryoSPARC helical refinement of 4,436 TMV segments, a RELION 3D auto-refinement of 24,973 T20S particles from session 2, and a cryoSPARC helical refinement of 1,648 of the 1,653 Phi92 segments in the End2EndPtychoSPA particle table (Punjani *et al*., 2017; He & Scheres, 2017).

### 2.3. Forward model

End2EndPtychoSPA extends end-to-end ptychographic tomography (You *et al*., 2025) to particles in unknown orientations, treating each macromolecule as one tilt view of a shared three-dimensional potential. We first review the ptychographic tomography forward model in the notation of You *et al*. (2025) and then introduce the extensions for cryo single-particle analysis, keeping the same operator symbols where the two formulations agree. Figure 2 shows an example apoferritin reconstruction item together with its resampled diffraction data on the model detector grid. Table S1 lists the symbols, and Supporting Information S1 gives the full equations for the apoferritin configuration. reconstruction

**Figure 2.**
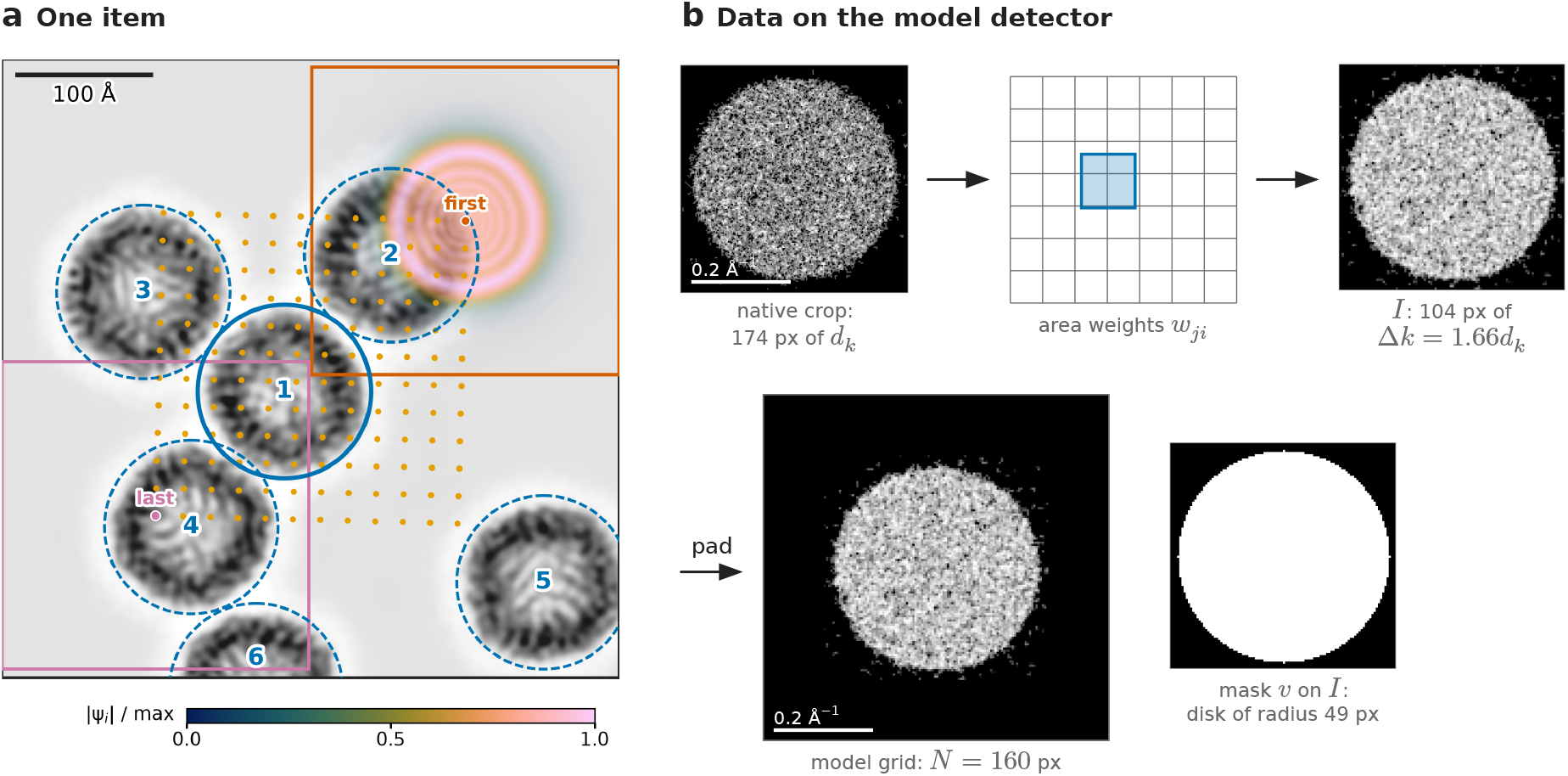
Geometry and detector representation of a single End2EndPtychoSPA reconstruction item from the apoferritin dataset. (a) Real-space geometry of an exemplary picked particle. Grey: the projected electrostatic potential sum *V*_*i*_ of the six particle copies within the item field. Orange dots: the 12 × 12 scan positions, spaced 20.6 Å apart. Squares: the probe windows (160 model pixels or 230 Å across) of the first (vermillion) and the last (purple) scan position. The item field covers the 12 × 12 scan window plus one probe window margin. Numbered circles mark all included particle copies *N*_*i*_ (copy 1, solid: the central target particle), drawn with the 65 Å inclusion radius of the neighbour selection rule. Copy 6 is cut by the field edge. Colour overlay: the incident probe amplitude |*ψ*_*i*_| at the first scan position, relative to its maximum; its opacity follows the amplitude. (b) Diffraction data on the model detector grid. Each native 174-pixel detector crop is integrated onto model pixels of width 1.66*d*_*k*_ using exact area weights, producing a 104 × 104 pixel block that is zero-padded to the 160 × 160 pixel model grid. The Poisson data loss is evaluated inside the central bright-field pupil disk of 49 model pixels in radius (mask *v*). All patterns are drawn at one reciprocal-space scale (scale bars: 0.2 Å^−1^). Thus the 104-pixel block covers almost the same area as the native crop (the model pixels that the crop covers completely), and the padded model grid is larger. All panels display tensors extracted from stage 2, epoch 15 of the apoferritin refinement. Grey values are scaled individually for visual clarity (native crop: square-root intensity scale; model-grid patterns: logarithmic scale).

#### 2.3.1. End-to-end ptychographic tomography

In end-to-end ptychographic tomography (You *et al*., 2025), 4D-STEM diffraction datasets of a single specimen with three-dimensional potential *V* are recorded across *N*_angles_ stage tilts, where tilt *l* is parameterized by Euler angles Θ_*l*_ and in-plane translation *τ*_*l*_. For each tilt, the affine transformation operator 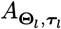 rotates and shifts the three-dimensional sampling coordinates, the axial binning operator *B*_*b*_ sums the rotated potential along the beam direction into *T* slices of thickness *b* voxels, and the cropping operator 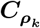 extracts the local two-dimensional patch of each slice illuminated at scan position *ρ*_*k*_. The multislice operator then alternates phase-grating transmission *T* and Fresnel free-space propagation *P*^Δ*z*^ over the slice thickness Δ*z* across the *T* potential slices *V*_*j*_ (*j* = 1, …, *T*). The multislice operator *ℳ*_*ψ*_ maps an incident wave *ψ* and the slices *V*_*j*_ to the exit wave (Cowley & Moodie, 1957; Maiden, Humphry, & Rodenburg, 2012; Tsai et al., 2016; Chen et al., 2021; Diederichs et al., 2024):

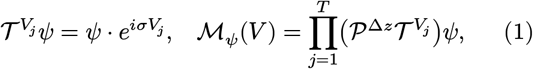

where *σ* denotes the relativistic electron interaction constant. The potential *V* is complex: its real part *ℜ*(*V*) gives the phase shift and its imaginary part ℑ(*V*) gives the amplitude attenuation. You *et al*. absorb the crop into a batched multislice operator 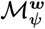; we write 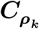 explicitly. With these definitions in place, the forward exit-wave model for scan position *k* at tilt *l* and the corresponding predicted diffraction intensity take the form

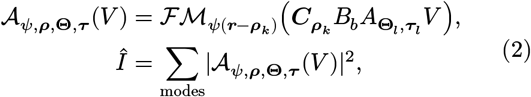

where *r* is the real-space coordinate, *ℱ* denotes the two-dimensional Fourier transform and the incoherent sum runs over orthogonal mixed-state probe modes (Thibault & Menzel, 2013). Their Algorithm 3 (End2EndPtychoTomo) jointly optimizes *V*, the probe modes *ψ* and the scan positions *ρ* (Maiden, Humphry, Sarahan et al., 2012) tilt by tilt using minibatch stochastic gradient descent (Tripathi et al., 2024) on the amplitude loss

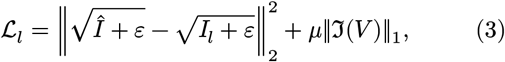

where *ε* is a small positive constant that keeps the square root differentiable at zero, *μ* is the weight of the *L*_1_ penalty, *I*_*l*_ contains the measured intensities for tilt *l* and an *L*_1_ shrinkage step is applied to ℑ(*V*) after each minibatch. The tilt orientations Θ_*l*_ and translations *τ*_*l*_ are estimated before 3D refinement by joint alignment of 2D ptychographic phase projections (their Algorithm 2) and are not updated in End2EndPtychoTomo.

#### 2.3.2. Extension to single-particle analysis with unknown particle poses

In cryo-EM single-particle analysis, the specimen consists of thousands of identical macromolecules frozen in random orientations and positions across multiple micrographs. If each particle *p* is identified with a single tilt view *l*, the tomographic forward model carries over with the tilt index *l* replaced by the particle index *p* and with the stage alignment (Θ_*l*_, *τ*_*l*_) replaced by the unknown particle pose (Θ_*p*_, *τ*_*p*_). Just as joint linear tomography provides the initial tilt alignment in You *et al*., the 2D ptychographic SPA workflow provides initial estimates of the particle poses (Θ_*p*_, *τ*_*p*_), which End2EndPtychoSPA refines jointly with the shared three-dimensional potential *V*.

Adapting end-to-end ptychographic tomography to cryo single-particle 4D-STEM data requires four changes:

1. **Particle pose refinement**. In end-to-end ptychographic tomography (You et al., 2025), orientations and translations are refined from phase projections and held fixed during end-to-end reconstruction. Here, particle poses are likewise initialized from the 2D SPA refinement, but may retain errors associated with the reconstructed images and projection approximation. We therefore refine them locally together with *V* against the original diffraction data, using a prior to constrain departures from the initial poses.
2. **Overlapping particles within a single probe window**. A tomographic tilt series images a single isolated object, whereas a vitrified cryo-EM specimen contains nearby macromolecules whose scan windows overlap. A single probe footprint can intersect multiple particles at once, so we model the local potential of item *i* as a sum of posed copies of *V* (Equation 8).
3. **Unknown particle height in the vitreous ice layer**. Because individual particles sit at different heights *z* in the ice layer, we introduce a learnable per-item free-space propagation distance 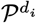 before the potential slices to absorb local height and defocus variations (Equation 10), shared by all particle copies in item *i*.
4. **Shared macromolecular identity and internal symmetry**. Because every particle is a copy of the same macromolecule, often with point-group or helical symmetry, we accumulate gradients across all particles into a single shared volume *V* and apply the point-group or helical symmetry to the volume gradient (Section 2.4). Particle heterogeneity is not considered yet.

With these extensions, the forward model for diffraction pattern *k* in item *i* can be written as

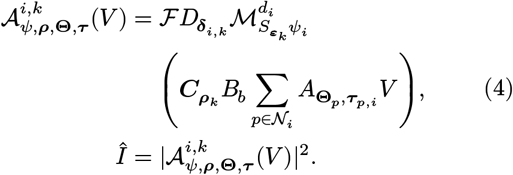

Here *ψ*_*i*_ is the incident probe of item *i* (Section 2.3.4), *N*_*i*_ is the set of particle copies in item *i* and *τ*_*p*,*i*_ is the lateral offset of particle *p* from the item centre (both defined with Equation 8). Compared with Equation 2, Equation 4 introduces three additional operators. First, the sub-pixel shift operator 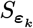 translates the item probe *ψ*_*i*_ by the fractional pixel remainder ***ε***_*k*_ of scan position *ρ*_*k*_ that cannot be represented by the integer cropping operator 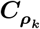 (Equation 5). Second, the modified multislice operator 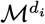 prepends a free-space propagation over the learnable axial offset *d*_*i*_ of item *i* (Equation 10). Third, the exit-wave phase ramp 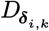 applies a sub-pixel reciprocal-space shift ***δ***_*i*,*k*_ to account for residual beam descan motion on the detector. In the special case of a single isolated particle per item (*N*_*i*_ = {*p*}), zero axial offset (*d*_*i*_ = 0), unit descan ramp (*D*_***δ***_ = 1) and fixed poses, Equation 4 reduces to the single-mode form of Equation 2 with particle *p* replacing tilt *l*. Table 2 compares End2EndPtychoTomo and End2EndPtychoSPA, and Algorithm 1 gives the full reconstruction loop. Below we define each operator and parameter for the apoferritin reconstruction.

**Table 2.**
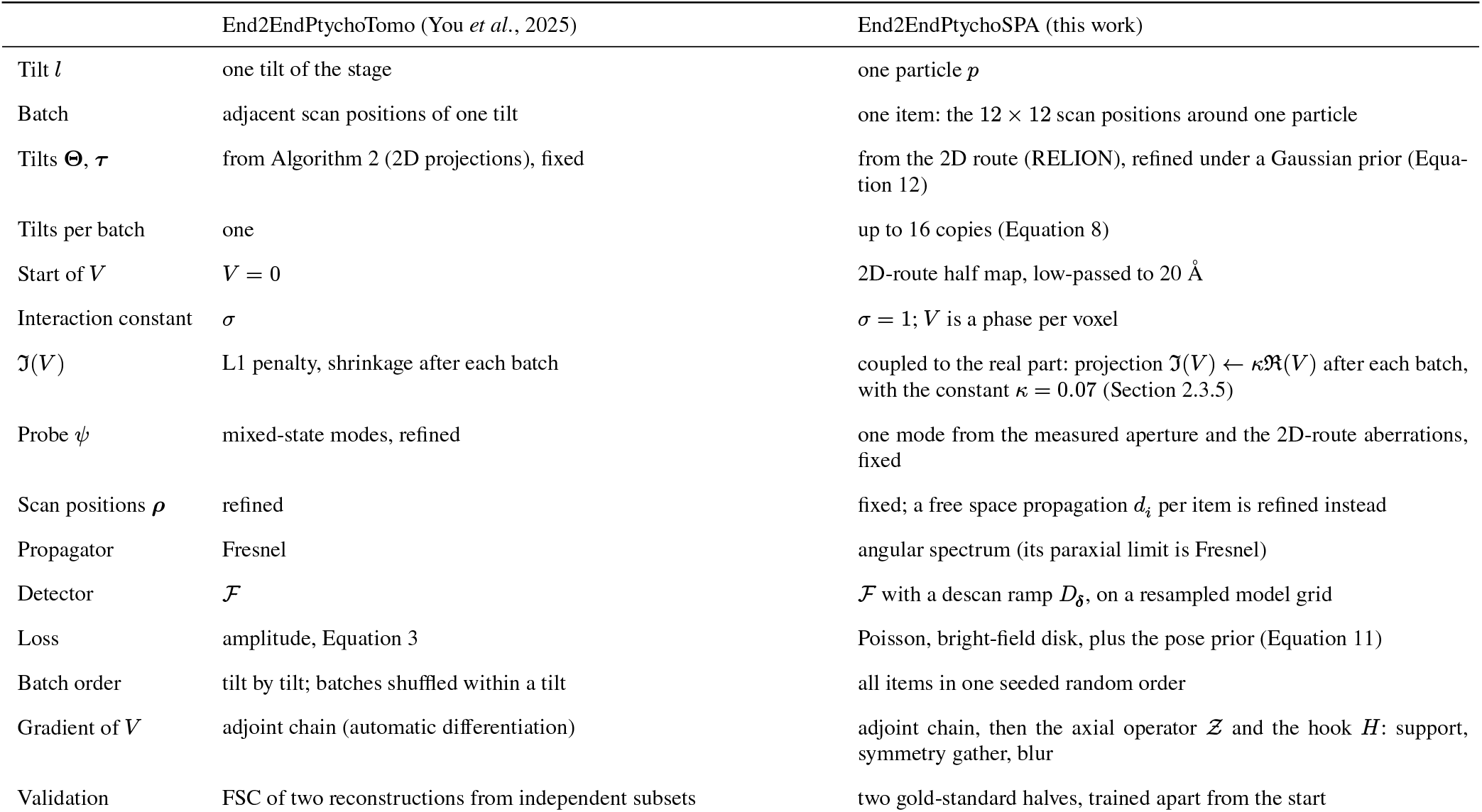
Comparison between end-to-end ptychographic tomography (You et al., 2025) and End2EndPtychoSPA for cryo single-particle analysis. Both algorithms share the three-dimensional volume representation *V* and the core operators *A*_Θ,*τ*_, *B*_*b*_, *C*_*ρ*_, *T*, *P, ℳ* and *ℱ*, and both optimize the volume directly against 4D-STEM diffraction patterns via minibatch stochastic gradient descent.

|  | End2EndPtychoTomo (You *et al.*, 2025) | End2EndPtychoSPA (this work) |
|---|---|---|
| Tilt *l* | one tilt of the stage | one particle *p* |
| Batch | adjacent scan positions of one tilt | one item: the 12 × 12 scan positions around one particle |
| Tilts **Θ**, ***τ*** | from Algorithm 2 (2D projections), fixed | from the 2D route (RELION), refined under a Gaussian prior (Equa-tion 12) |
| Tilts per batch | one | up to 16 copies (Equation 8) |
| Start of *V* | *V* = 0 | 2D-route half map, low-passed to 20 Å |
| Interaction constant | *σ* | *σ* = 1; *V* is a phase per voxel |
| ℑ(*V*) | L1 penalty, shrinkage after each batch | coupled to the real part: projection ℑ(*V*) ← *κ*ℜ(*V*) after each batch, with the constant *κ* = 0.07 (Section 2.3.5) |
| Probe *ψ* | mixed-state modes, refined | one mode from the measured aperture and the 2D-route aberrations, fixed |
| Scan positions ***ρ*** | refined | fixed; a free space propagation *d*<sub>*i*</sub> per item is refined instead |
| Propagator | Fresnel | angular spectrum (its paraxial limit is Fresnel) |
| Detector | ℱ | ℱ with a descan ramp *D*<sub>***δ***</sub>, on a resampled model grid |
| Loss | amplitude, Equation 3 | Poisson, bright-field disk, plus the pose prior (Equation 11) |
| Batch order | tilt by tilt; batches shuffled within a tilt | all items in one seeded random order |
| Gradient of *V* | adjoint chain (automatic differentiation) | adjoint chain, then the axial operator 𝒵 and the hook *H*: support, symmetry gather, blur |
| Validation | FSC of two reconstructions from independent subsets | two gold-standard halves, trained apart from the start |

#### 2.3.3. Reconstruction items and scan geometry

End2EndPtychoSPA divides the dataset into particle-centred reconstruction items. Each item consists of the 12 × 12 scan positions around one picked particle from the input table (144 diffraction patterns per item, forming one minibatch). Any candidate item whose 12 × 12 scan window extends beyond the boundary of the 4D-STEM scan is excluded. Such particles still enter neighbouring items as particle copies. For apoferritin, this rule leaves 5,710 and 5,440 items in the two half sets (11,150 of the 11,552 particles; the 2D route uses all 11,552). For T20S, the run also caps each half set at 9,000 items, leaving 8,984 and 8,986 items of the 24,973 particles. Nearby particles on the same micrograph can share diffraction patterns across overlapping 12 × 12 windows, but because the half sets are split by scan, shared diffraction patterns never cross between the two half sets. For apoferritin, the calibrated scan step is Δ*r* = 20 × 1.03 = 20.6 Å, where the scale factor of 1.03 corrects the nominal 20 Å scan step (Li, Li et al., 2026), and the scan-to-detector rotation angle *θ*_*s*_ for each scan is taken from its initial SSB fit. The probe with scan index *k* has the integer raster position ***t***_*k*_, which maps into object-grid pixel coordinates according to

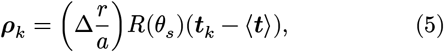

where *a* is the real-space object pixel size (Equation 6), *R*(*θ*_*s*_) is the two-dimensional rotation by *θ*_*s*_ and ⟨***t***⟩ is the mean raster position of the scan. The integer part of *ρ*_*k*_ determines the real-space crop window 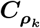, while the fractional remainder ***ε***_*k*_ is applied to the incident probe through the sub-pixel Fourier phase-ramp shift operator 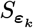 of You *et al*.

The native detector has a reciprocal-space pixel pitch of *d*_*k*_ = 0.00261823 Å^−1^. Around the bright-field disk of each frame, we first extract a 174 × 174 native-pixel crop and, at data loading, integrate this crop onto coarser model detector pixels of pitch Δ*k* = 1.66*d*_*k*_ using exact area weighting to conserve electron counts. Only the 104 × 104 model pixels per frame that are completely covered by the native crop enter the data loss. Embedding this block into a *N* = 160 pixel model Fourier grid yields a real-space object pixel size of

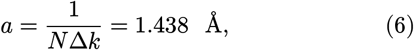

corresponding to a real-space probe window width of 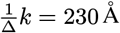. Because the dark-field detector region in low-dose biological 4D-STEM is dominated by shot noise, we evaluate the data loss only inside a circular pupil mask of radius 49 model pixels centred on the bright-field disk (whose nominal 4 mrad radius corresponds to about 47 model pixels, plus a small margin).

#### 2.3.4. Incident probe, lens aberrations and beam descan

We construct the incident probe wavefunction for each scan from its recorded vacuum aperture intensity. The aperture amplitude *A*_*s*_ is area-resampled onto the model detector grid in the same manner as the diffraction data, and its reciprocal-space phase is synthesized from all 12 coherent aberration coefficients refined for that scan during the 2D ptychographic step:

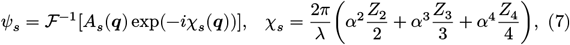

where *s* is the scan index, ***q*** is the reciprocal-space coordinate, *α*= *λ* |***q***| is the scattering semi-angle, *λ* is the relativistic electron wavelength at 300 kV and *Z*_*n*_ denotes the azimuthal aberration polynomials of radial order *n* (defined explicitly in Supporting Information S1). For each item *i*, we normalize the incident probe intensity to match the mean measured electron counts of that item within the 49-pixel bright-field pupil disk; this gives the item probe *ψ*_*i*_. The probe is represented by a single coherent mode and remains fixed throughout 3D refinement.

To account for residual shift of the diffraction pattern across the detector during beam scanning (descan), we fit a two-dimensional linear plane to the bright-field centre of mass across all frames of each scan. The integer-pixel component of this fitted plane shifts the native 174-pixel crop window on a frame-by-frame basis, while the residual sub-pixel offset ***δ***_*i*,*k*_ is applied to the exit wave via the real-space phase ramp 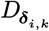, shifting the predicted far-field pattern by the exact fractional pixel remainder.

#### 2.3.5. Shared volume representation, particle copies and neighbour inclusion

We represent the macromolecule as a single shared complex three-dimensional array *V* discretized on a grid of 168 × 322 × 322 voxels (*zyx*, corresponding to an axial thickness of 241 Å). Setting *σ* = 1 in Equation 1, the real part *ℜ*(*V*) represents the electrostatic phase shift per voxel in radians (initialized with the scale described in Section 2.4), and the imaginary part ℑ(*V*) models amplitude attenuation. Instead of an *L*_1_ shrinkage step on ℑ(*V*) as in Equation 3, we couple the imaginary part to the real part after each minibatch update via ℑ(*V*) = *κℜ*(*V*) with *κ* = 0.07, so that a single real 3D potential sets both the elastic phase shift and the amplitude contrast. This is the complex constraint of coherent diffractive imaging (Clark et al., 2010), which is also used in ptychography (Xu et al., 2024).

Each picked particle *p* is parameterized by its Euler angles Θ_*p*_, with rotation matrix 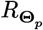, and its in-plane translation *τ*_*p*_. The local three-dimensional potential *V*_*i*_ illuminated within item *i* is the sum of posed copies of *V* over all particles whose centres fall within the item’s extended object field:

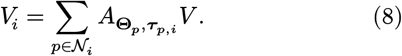

Here, the affine sampling operator 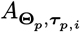 evaluates *V* by trilinear interpolation at coordinates 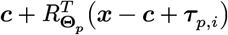, where ***x*** denotes the voxel coordinates of the item volume, ***c*** denotes the volume centre and *τ*_*p*,*i*_ is the lateral offset of particle *p* relative to the centre of item *i*. The index set *N*_*i*_ orders the central target particle of item *i* first, followed by its nearest neighbours up to a maximum of 16 copies, where any particle whose centre lies within 65 Å of the item’s object field boundary is included as a neighbour (apoferritin values; the haemoglobin simulation uses 35 Å and at most 64 copies, Table S7). To keep the two gold-standard half sets independent, neighbour copies are drawn only from the same half set. Because each neighbour particle shares its global pose parameters (Θ_*p*_, *τ*_*p*_) across every item in which it appears, gradients with respect to its pose are accumulated across all overlapping items.

#### 2.3.6. Multislice propagation and per-item defocus offset

Following trilinear sampling of the composite item volume *V*_*i*_, the axial binning operator *B*_*b*_ with bin factor *b* = 28 sums the potential over groups of 28 fine *z*-planes, yielding *T* = 6 potential slices *V*_*i*,*j*_ = [*B*_*b*_*V*_*i*_] of thickness Δ*z* = 28*a* = 40.3 Å, each acting via the phase-grating transmission operator 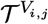 of Equation 1. A slice thickness of 40.3 Å (4.0 nm) satisfies the multislice sampling bound of < 6 nm found in dynamical simulations of proteins in vitreous ice at 300 kV (Leidl et al., 2023). Free-space propagation over an axial distance *d* is computed using the angular-spectrum propagator without its constant carrier phase,

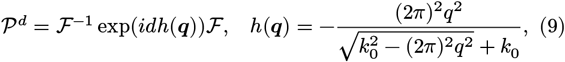

where *q* = |***q***| and 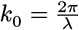 is the electron wavenumber; in the paraxial limit, Equation 9 reduces to the Fresnel propagator used by You *et al*. Upstream of the first potential slice, we propagate the sub-pixel-shifted probe across a learnable per-item free-space distance *d*_*i*_, adapting the adaptive free-path formulation of Cui et al. (2025) for tilted-surface ptychography to represent a single axial offset per item without a lateral tilt ramp. The multislice propagation operator then takes the form

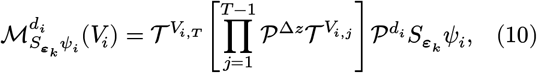

where the operators act from right to left as the slice index *j* runs from 1 to *T*, and the cropping operator 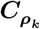 extracts the local probe window of each slice *V*_*i*,*j*_ at scan position *ρ*_*k*_. Unlike Equation 1, no post-specimen propagation is applied after slice *T*. The per-item offset *d*_*i*_ acts as a local defocus adjustment shared by all particle copies within item *i*. It absorbs vertical particle height variations within the ice layer as well as local defocus drift across the scan; we have not tested whether the fitted values measure particle heights, because all copies of an item share one axial plane and no independent height measurement exists for these data.

#### 2.3.7. Detector intensity prediction, Poisson loss and pose prior

The predicted detector intensity 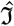 for scan position *k* in item *i* is given by the single-mode forward model in Equation 4, while the experimental measurements are stored as amplitudes 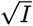 after dividing the raw detector counts by the scan’s empirical SSB noise gain. Instead of the amplitude loss of Equation 3, we use a Poisson negative log-likelihood loss (omitting data-only constants) averaged over all valid detector pixels *v*:

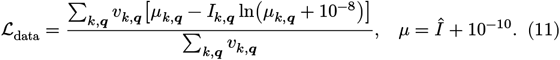

Because diffraction patterns in the overlap region between neighbouring items enter the loss once per item, and because fractional area resampling and noise-gain normalization modify the raw counting statistics, Equation 11 is a composite Poisson quasi-likelihood rather than an exact independent-frame likelihood.

In ptychographic tomography the stage tilts remain fixed during the end-to-end reconstruction, whereas here the particle poses are refined, and we regularize their updates with a Gaussian prior around the initial poses. At the epoch where pose refinement is activated, the current rotation matrices 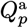 and lateral translations 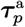 are recorded as anchor poses, marked by the superscript a, for the penalty

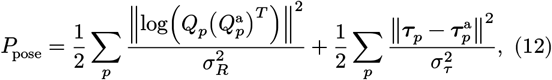

with 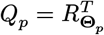. Here log is the logarithm map of the rotation group SO(3), written as a rotation vector, so its norm is the angle of the rotation from the anchor pose to the current pose. We use the rotational standard deviation *σ*_*R*_ = 2° and the translational standard deviation *σ*_*τ*_ = 3.3 object pixels (4.7 Å). This penalty is normalized by the number of detector block entries per minibatch, *n*_data_ = 144 × 104 × 104 (valid or not), and by the number of minibatches per epoch, *B*_epoch_, before being added to the batch loss.

### 2.4. Optimization and spatial-frequency gradient filters

#### 2.4.1. Optimizers and parameter updates

Algorithm 1 summarizes the joint optimization procedure, which follows Algorithm 3 of You *et al*. (2025) step by step while incorporating the single-particle extensions listed in Table 2. The two gold-standard half sets are refined as independent processes on separate GPUs using the same random seed. We update the three-dimensional volume *V* using stochastic gradient descent (SGD) with step size *γ*_1_ = 10, and update the particle Euler angles and lateral positions using SGD with step sizes *γ*_Θ_ = 10 and *γ*_*τ*_ = 10^4^, respectively. Each per-item defocus offset *d*_*i*_ is updated via a signed-gradient step of magnitude *γ*_*d*_ = 20 Å per visit, regularized by a weak quadratic tether 2*wd*_*i*_ with weight *w* = 6 × 10^−12^, while the incident probe and scan positions remain fixed.

For each item, the backpropagated update direction *g* for the shared volume *V* follows the adjoint backpropagation chain of You *et al*. (their Eqs. 18–19) summed over all included particle copies *p ∈ N*_*i*_, with the addition of an axial smoothing operator *Ƶ*:

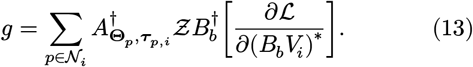

##### Algorithm 1

End2EndPtychoSPA joint refinement of the shared 3D potential volume, particle poses and per-item defocus offsets via minibatch SGD. Each gold-standard half set executes the algorithm independently on its assigned particles. Annotations in the right-hand column indicate the corresponding step in Algorithm 3 of You *et al*. (2025). For the apoferritin reconstruction, this procedure is executed across two sequential stages (Section 2.4.3).

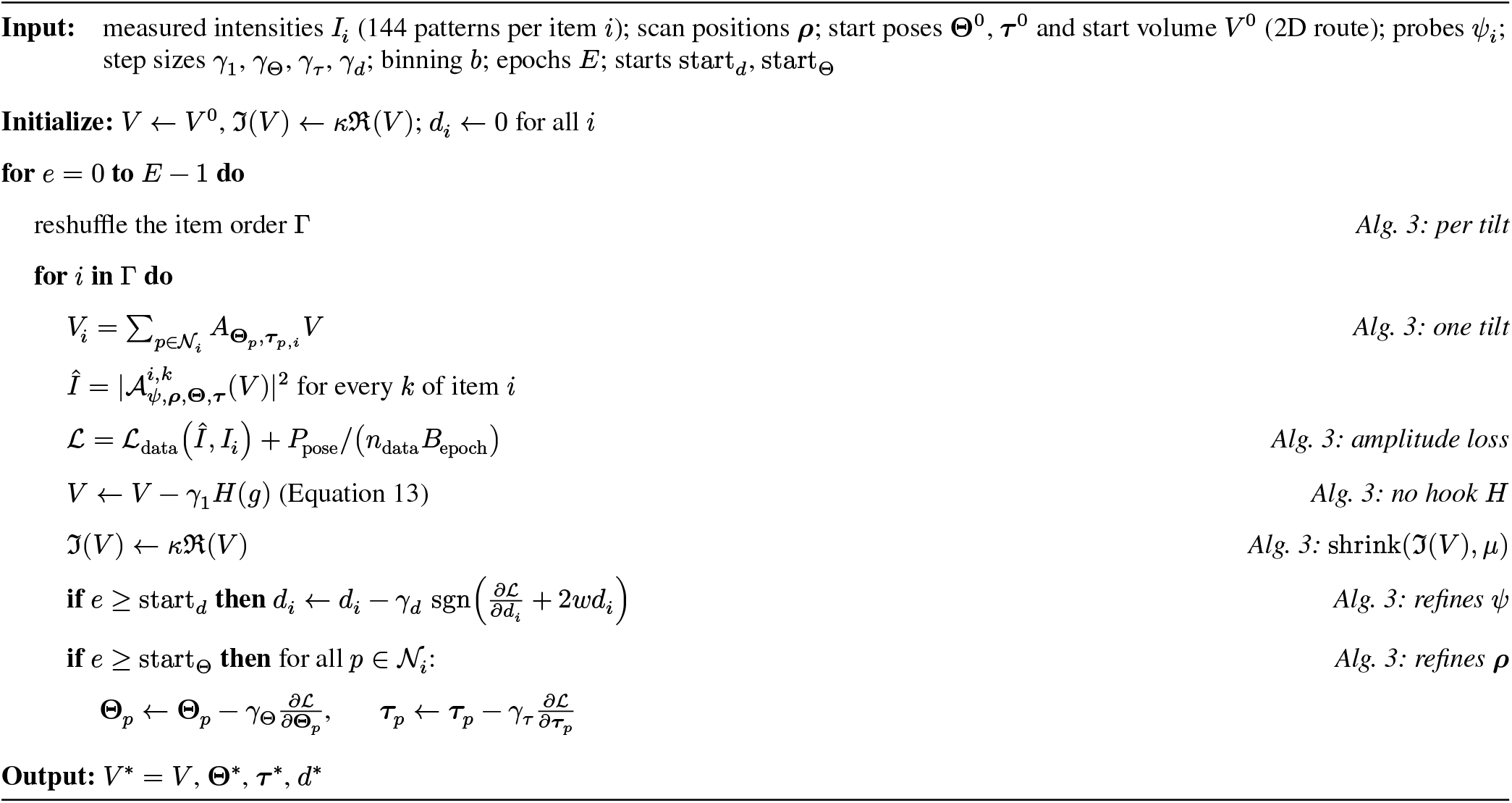

Here, † denotes the adjoint operator, the derivative is taken with respect to the complex conjugate (Wirtinger calculus), and *Ƶ* is a backward-only one-dimensional Gaussian smoothing filter applied along the beam axis *z* prior to trilinear adjoint splatting. Empirically this combines smooth gradients along the beam with cheap forward and backward passes. More expensive interpolation schemes may yield slightly better accuracy. Automatic differentiation evaluates the bracketed slice gradient through *ℱ*^†^, *ℳ*^†^ and 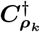, after which the gradient hook *H* preconditions *g* prior to the volume update.

#### 2.4.2. Spatial-frequency and symmetry gradient filters

Prior to each volume update, the gradient hook *H* applies four operations within the central cubic subregion of *V* : (i) multiplication by a soft spherical support mask *m*, which equals unity inside radius *r*_0_ = 72 Å and tapers smoothly to zero at 96 Å via a raised-cosine edge; (ii) point-group symmetrization *S*, which for apoferritin averages the gradient over the 24 proper rotations of the octahedral group O; (iii) isotropic three-dimensional Gaussian low-pass filtering *G* with standard deviation *σ*_*G*_ = 4 Å, evaluated in Fourier space using the exact Gaussian transfer function on a zero-padded grid; and (iv) zero-padded embedding of the central cube back into the full 168 × 322 × 322 volume. Because the four-fold symmetry axes of O are aligned with the Cartesian grid axes, every group operation is an exact signed coordinate permutation that maps grid voxels onto grid voxels without interpolation.

The order of these linear operations matters for the independence of the gold-standard half maps at high spatial frequencies. The hook applies the support mask, symmetrizes, re-applies the support mask and applies the Gaussian low-pass filter last (‘blurlast’ ordering):

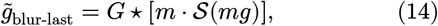

where ⋆ denotes convolution. A support mask applied after the low-pass filter would convolve the low-frequency update, which the two half maps share, with the Fourier transform of the mask.

This puts correlated power into high-frequency shells beyond the low-pass cutoff. With *G* last in Equation 14, the Gaussian transfer function attenuates the high-frequency ringing of the spatial mask. The final blur also widens the update beyond the outer mask edge. We call this widening the blur margin and take it as 3*σ*_*G*_, rounded up to whole voxels: ⌈(3*σ*)_*G*_/*a*⌉*a* = 12.9 Å. A box check confirms that the mask radius, its raised-cosine edge and the blur margin fit inside the central cube, so that the cube faces do not cut off the blurred update. If they do not fit, a raised-cosine face taper (at least 20 Å wide) rolls the mask to zero one blur margin inside the cube faces. For apoferritin, mask radius, edge and blur margin add up to 72 + 24 + 12.9 = 108.9 Å, which is less than the half-box extent of 120.1 Å, so no face taper is applied. The hook *H* acts identically on the real and imaginary components of the gradient and modifies only the update direction rather than projecting the accumulated volume. Figure 3 shows the spectral effect of the blur-last ordering on a synthetic low-frequency update.

**Figure 3.**
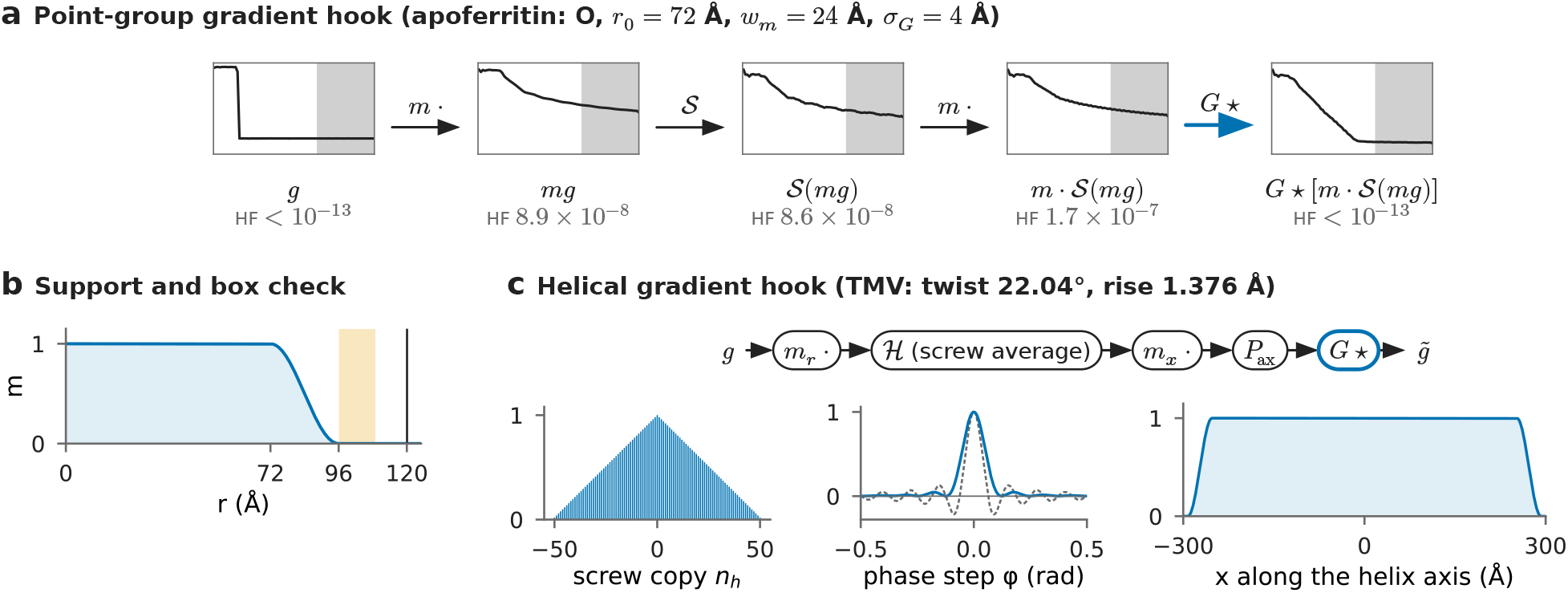
Computational graphs and spectral transfer behaviour of the volume gradient hooks. (a) Point-group gradient hook for apoferritin in the blur-last ordering, acting on a synthetic update shared between both halves that contains power exclusively below 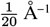. Each box displays the radially averaged power spectrum after the corresponding operation, on a log scale from |***k***| = 0 to Nyquist. The grey region marks the high-frequency (HF) diagnostic band |***k***| ≥ 1/4.5 Å^−1^ the value below each box (HF) is the fraction of the total power in that band. The multiplications by the spatial mask *m* move up to 1.7 × 10^−7^ of the update power into the HF band; the final Gaussian filter lowers the HF power fraction below 10^−13^. (b) Radial profile of the spherical support mask *m*(*r*) and bounding-box clearance check for the blur-last ordering. Orange: the blur margin ⌈3*σ*_*G*_/*a*⌉*a* = 12.9 Å after the mask edge at 96 Å. Black line: the half extent of the cube, Å. (c) Helical gradient hook used for TMV. Left: triangular Fejér weights 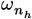 of the screw-averaging operator *ℌ*(*n*_max_ = 50). Centre: eigenvalue of *ℌ*as a function of the phase step. The Fejér weights (blue curve) guarantee a non-negative spectral transfer function, whereas uniform weights (dashed curve) produce negative spectral sidelobes. Right: axial profile of the face taper *m*_*x*_ along the helical axis.

In addition to the three-dimensional hook *H*, the backward-only operator *Ƶ* in Equation 13 smooths the gradient of the sampled potential stack along the beam direction *z* immediately after adjoint slice unbinning *B*^†^. This filter applies a one-dimensional Gaussian kernel with standard deviation 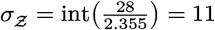 fine voxels truncated to a 23-voxel window; its forward pass is the identity operator, and its backward pass (which is not the adjoint of the forward pass) smooths both the volume gradient and the pose gradient.

#### 2.4.3. Gold-standard half sets, initialization and training schedule

End2EndPtychoSPA keeps the two scan-pure R1 particle half sets from the 2D route, so the two halves share no 4D-STEM scans, no detector frames and no neighbour particle copies. Each half set is initialized independently from its own R1 RELION half map to form *V* ^0^ and from the particle poses (Θ_*p*_, *τ*_*p*_) refined by RELION within that half set.

We construct the initial volume *V* ^0^ for each half set in three steps. First, we resample the corresponding RELION half map onto the voxel grid of *V*, apply a 20 Å Gaussian low-pass filter, clip negative voxel values to zero, and scale the map into radians by fitting the projected map to the SSB phase micrographs of that half set via least squares and multiplying the resulting slope by 0.5. Second, we embed the resampled map into the 168 × 322 × 322 volume box and invert it along the beam axis *z* to match the handedness convention of the pose parameterization.

Third, we zero all Fourier shells beyond 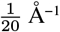 on the full three-dimensional voxel grid of *V* using a raised-cosine roll-off over two Fourier shells, so that the two initial volumes *V* ^0^ share no spatial frequencies finer than 20 Å.

For apoferritin, we execute Algorithm 1 across two sequential stages. Stage 1 begins from *V* ^0^ and runs for *E*= 8 epochs, activating per-item defocus refinement at epoch start_*d*_ = 2 and particle pose refinement at epoch start_Θ_ = 4. Stage 2 resumes from the volume *V*, poses (Θ_*p*_, *τ*_*p*_) and defocus offsets *d*_*i*_ saved at the end of stage 1 (epoch 7) and runs for *E*= 16 additional epochs with both defocus and pose refinement active from the start (start_*d*_ = start_Θ_ = 0). In each stage, the Gaussian pose prior in Equation 12 resets its anchor poses 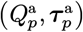 at epoch start, and the final apoferritin reconstruction is taken from the last epoch of stage 2.

#### 2.4.4. Specimen-specific adaptations for T20S, TMV and Phi92

For the T20S proteasome, ()the symmetry gather *S* under D7 symmetry uses trilinear interpolation, because seven-fold rotations do not map grid voxels onto grid voxels, and the refinement runs in three stages of 24, 16 and 8 epochs. For the helical specimens (TMV and Phi92), a Fejér-weighted screw-symmetry average *ℌ* about the helical axis replaces the point-group gather (Figure 3(c)), and the particle poses and defocus offsets are held fixed. As pre-declared, the final TMV reconstruction is taken from epoch 1 of 3, and the final Phi92 reconstruction from epoch 0 of 4 (the epoch with the highest 8–30 Å half-map band correlation). Section S2.1 gives the helical hook, its masks and the initialization.

### 2.5. Validation and quantitative evaluation metrics

#### 2.5.1. High-frequency rebound and phase-randomization diagnostic

Gold-standard resolution estimation requires that the two half-set reconstructions remain statistically independent (Henderson *et al*., 2012; Scheres & Chen, 2012). Because spatial-frequency gradient filters can couple the two halves at high spatial frequencies (Section 2.4), every reconstruction must pass a pre-declared high-frequency defect check on the full, unmasked native grid of its half maps: the pooled half-map FSC for 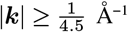 must not exceed 0.1, and a phase-randomization rebound check adapted from Chen *et al*. (2013) must pass. Section S2.2 gives all criteria and the specimen-specific rules. A reconstruction that fails the check is still reported, with its pre-registered comparison marked as not valid.

#### 2.5.2. Map-to-model evaluation metrics

To evaluate the reconstructed volumes, we compared each map, calculated as the mean of its two aligned half maps, with an independent atomic model or high-resolution cryo-EM map, after aligning it to the reference over handedness, isotropic voxel scale and sub-voxel translation (Section S2.3).

For each reconstruction, we report five map-to-model metrics:

- *C*: the real-space/Fourier band correlation with the reference model over the 8–30 Å spatial-frequency band.
- *C*_w_: the amplitude-whitened correlation over the 8–30 Å band, excluding Fourier shells associated with the scan lattice.
- FSC_10_ and FSC_8_: the map-to-model Fourier shell correlation evaluated at spatial frequencies of 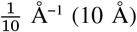 and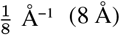.
- Model 0.5: the spatial resolution at which the map-to-model FSC first falls below 0.5.

*C*_w_ and the model 0.5 crossing exclude the Fourier shells of the scan lattice (Section S2.3). T20S omits *C*_w_ and is scored against the cryo-EM map EMD-6287 (Campbell et al., 2015), with PDB 1PMA as a secondary reference.

To reduce post-hoc selection bias, we pre-declared two quantitative decision thresholds before scoring the final End2EndPtychoSPA reconstructions. The benchmark threshold was the score of the deposited EMDB half-map mean plus a fixed margin. The start-improvement threshold was the score of the reconstruction’s own initialization plus the same margin. For apoferritin, these pre-declared margins are +0.010 in *C*_w_ and +0.030 in both FSC_10_ and FSC_8_. TMV uses the same margins applied to *C*, FSC_10_ and FSC_8_ T20S uses zero margins relative to its reference and start; and Phi92 evaluates *C* and FSC_8_ together with a reverse atomic-model fit as an additional bench-mark requirement.

Alongside the map-to-model metrics, we report the gold-standard half-map FSC. The half-map FSC was calculated without spatial masking and inter-half alignment on the full native voxel grid of each reconstruction. We used a shell width of 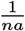, where *n* is the smallest box dimension, and report the resolution at which it crosses 0.143 (Rosenthal & Henderson, 2003; Heel & Schatz, 2005). For apoferritin, the high-frequency diagnostic found no shared high-frequency signal on the unmasked End2EndPtychoSPA grid (Section 3.2.1), so box-edge effects do not affect its unmasked half-map FSC. This does not hold in general: the helical reconstructions share correlations at the box boundaries (Section 3.2.3). We do not report spatially masked half-map FSC curves for End2EndPtychoSPA because multiplying the finished half maps by a tight spatial mask convolves shared 6–15 Å signal into higher-frequency shells.

#### 2.5.3. Scaling of resolution with particle number (ResLog analysis)

To measure how reconstruction quality scales with dataset size (ResLog analysis) (Rosenthal & Henderson, 2003), we drew nested random subsets containing 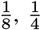 and 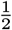 of the particles within each R1 gold-standard half set, keeping the scan-pure half-set assignments. Both workflows processed the same particle subsets: the 2D route re-ran RELION 3D auto-refine using the R1 settings on each subset, and End2EndPtychoSPA initialized from the resulting subset RELION half maps and poses using the apoferritin schedule. We define the resolution *d*_res_ at each subset size via the map-to-model FSC 0.5 crossing, and we also test sensitivity to scan-lattice shell placement and reference docking in Section 3.2.2 alongside the lattice-independent metric FSC. Fitting 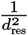 against ln *N* by ordinary least squares across the four particle counts (*N* denoting the total particles across both halves), we report the Rosenthal–Henderson *B*-factor and its standard error

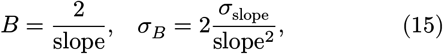

together with 95% confidence intervals computed from Student’s *t*-distribution with *N*_fit_ − 2 degrees of freedom for a fit to *N*_fit_ points (two for one series of four particle counts; three and six for the pooled End2EndPtychoSPA and 2D-route fits that include the second draw). To estimate variability across random subset draws, we performed a second random draw across all three subset fractions (plus a replicate full-*N* RELION run) for the 2D route and at 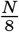 for End2EndPtychoSPA. Because the particle subsets are nested, their residuals are not independent, so the regression standard error assumes an independence that the data do not have and is not a complete measure of the sampling uncertainty. The End2EndPtychoSPA subset reconstructions also remain coupled to the 2D route through their initial maps and poses.

### 2.6. Ground-truth multislice simulations

To test End2EndPtychoSPA under controlled conditions where the ground-truth electrostatic potential and particle poses are known, we generated multislice cryo 4D-STEM datasets. In the primary reconstruction branch, RELION used a **de novo** initial volume from VDAM (variable-metric gradient descent with adaptive moments estimation) (Kimanius *et al*., 2021), and End2EndPtychoSPA used the resulting half maps and poses. Particle picking used low-resolution templates derived from the atomic model of the simulated protein, as described below. Two control branches used additional ground-truth inputs: a RELION refinement initialized from the 20 Å low-pass-filtered groundtruth model, and oracle reconstructions using the exact particle poses. The haemoglobin benchmark is presented in Section 3.1; the smaller myoglobin benchmark (Section S9) uses the same atomistic ice construction, electron-optical parameters and 2D reconstruction pipeline with a 60 Å ice thickness and 150 particles per micrograph.

Section S10 in the Supporting Information describes the atomistic specimen and ice model, the multislice 4D-STEM simulation, the 2D route on simulated data, the End2EndPtychoSPA configuration with the ice background model, and the groundtruth evaluation metrics. Section S11 gives the computational cost of all runs.

## 3. Results

We first test End2EndPtychoSPA on simulated multislice datasets with known ground truth (Section 3.1) and then compare it with the two-step 2D reconstruction route on four published experimental datasets (Section 3.2).

### 3.1. Ground-truth simulation of haemoglobin: joint pose and volume refinement with an explicit ice model

We used multislice simulations of macromolecules embedded in vitreous ice to assess how initial pose accuracy and solvent modelling affect end-to-end refinement. The known three-dimensional potential and particle poses provide references for evaluating reconstruction accuracy.

With the ice-background model, end-to-end refinement improved the map-to-truth FSC 0.5 resolution from 3.41 to 3.22 Å under the molecular mask and reduced the median orientation error from 2.64° to 1.79°. Without the ice model, the reconstruction deteriorated to 3.57 Å.

In simulations of sperm-whale myoglobin (17.9 kDa), global orientation search in the 2D route failed, and subsequent local End2EndPtychoSPA refinement did not recover the correct poses. Yet reconstruction of the same 2D projections using the true poses reached the Nyquist limit (Section S9, Figure S1), indicating that initial orientation recovery limited reconstruction under these conditions. We therefore used human deoxy-haemoglobin (PDB 2DN2, 64.5 kDa) as our primary ground-truth benchmark, for which the 2D route provided sufficiently accurate initial orientations for local refinement. We simulated 50 micro-graphs of deoxy-haemoglobin embedded in an 80 Å atomistic vitreous-ice layer under the same electron-optical conditions and 30 e/Å2 dose used for myoglobin (Section S10, Figure 4). Automated template picking on the direct ptychography phase micrographs achieved a pooled *F*_1_ score of 0.98 within the picking evaluation region (35 Å boundary margin; 10 Å matching tolerance; Section S10, Figure 4(b)).

**Figure 4.**
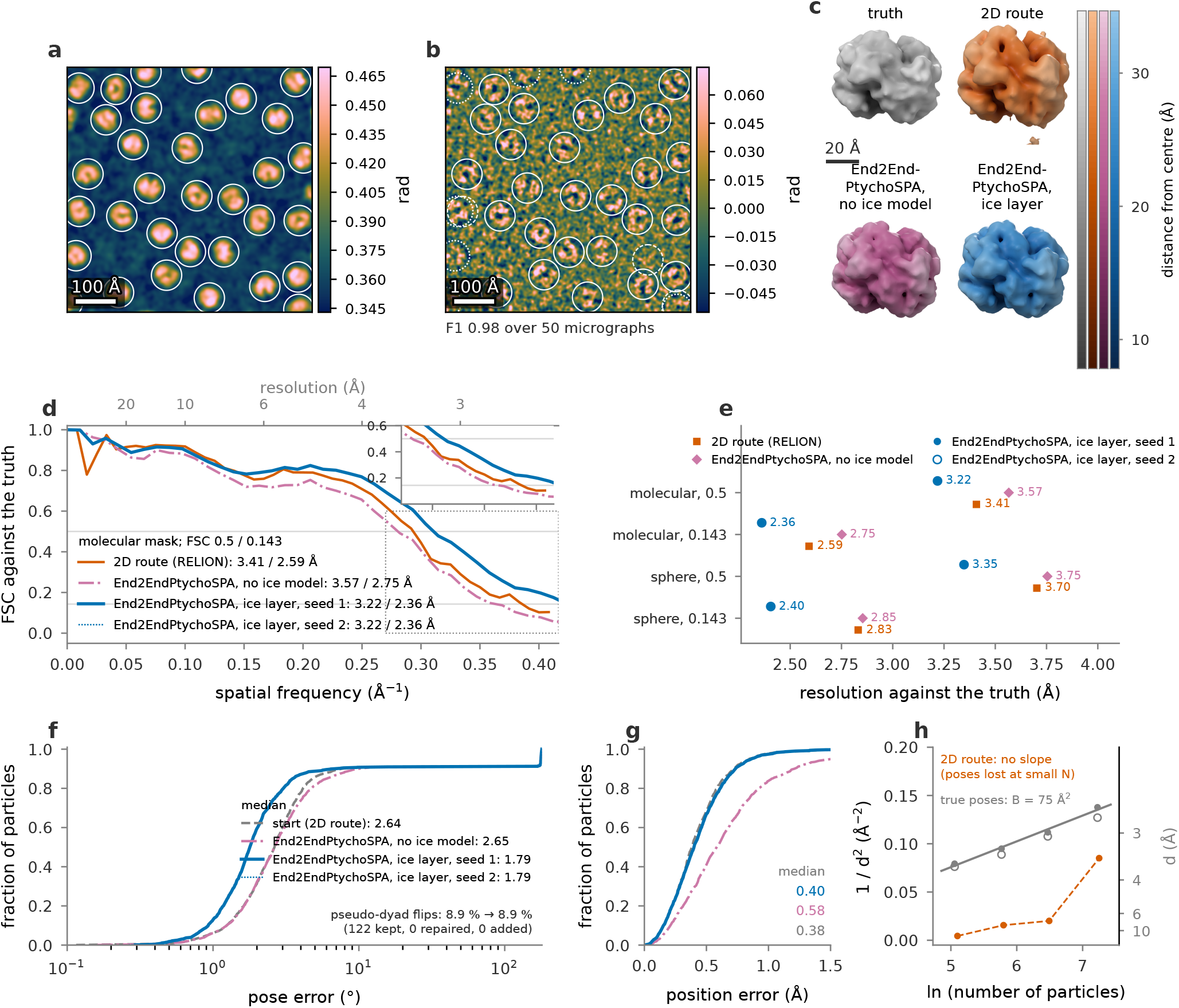
Ground-truth multislice simulation of human deoxy-haemoglobin (PDB 2DN2, 64.5 kDa). The 2D route uses RELION initialized from its own **de novo** VDAM model in C1, while End2EndPtychoSPA starts from the resulting RELION half maps and poses. (a) Ground-truth projected phase of micrograph 0, with circles marking the true particle centres; Gaussian-smoothed for display with *σ* = 5 Å. (b) Reconstructed 2D ptychographic phase micrograph overlaid with template picks (solid circles: matched to a true particle; dashed: unmatched false positives; dotted: missed true particles); Gaussian-smoothed for display with *σ* = 3 Å. (c) Isosurface renderings of the reconstructed maps aligned to the ground truth, Gaussian-smoothed with *σ* = 2 Å and contoured at the molecular volume. (d) Map-to-truth FSC curves evaluated inside the tight molecular mask, with FSC 0.5 and 0.143 crossings indicated in the legend. (e) Summary of FSC 0.5 and 0.143 crossings under both the molecular mask and the 42 Å soft sphere across reconstruction variants. The 2.36 Å molecular-mask crossing lies beyond the reference grid’s 2.4 Å Nyquist limit and cannot establish resolution at that scale; the spherical-mask crossing, approximately 2.40 Å, lies close to this limit. (f, g) Distributions of angular orientation error and in-plane translation error across the 1,374 particles matched to a ground-truth particle (out of 1,409 particles in RELION), compared against the initial RELION poses (dashed curves); inset annotations report the fraction of pseudo-dyad flips before and after refinement. (h) ResLog scaling of the 2D route (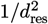 versus ln *N*, where *d*_res_ is the unmasked half-map FSC 0.143 crossing and open symbols denote the map-to-truth FSC 0.5 crossing). Because smaller particle subsets fail to align all particles **de novo**, no linear fit is shown for the **de novo** branch; the grey curve shows the oracle-pose control together with its linear fit *B* = 2/slope. No End2EndPtychoSPA subset series was run on haemoglobin.

Both routes used the same 1,409 particles and half-set assignments; pose errors were evaluated on the same 1,374 one-to-one matches within 10 Å of truth. Every particle contributed to end-to-end optimization, including those represented only as neighbouring copies (Table S7).

RELION refined the particle projections from a **de novo** VDAM initial model in C1, reaching 3.41 Å at map-to-truth FSC 0.5 under the molecular mask and a median orientation error of 2.64° among the matched particles. Reconstructing the 1,374 truth-matched 2D projections using the true particle poses reached 2.81 Å at FSC 0.5.

Because the *α*and *β* subunits of haemoglobin have similar folds (pseudo-*D*_2_ symmetry), 8.9% of the orientations assigned by RELION were flipped by about 180° around a pseudo-dyad axis.

We initialized End2EndPtychoSPA from the RELION half maps and poses, selecting hyperparameters without consulting the ground truth. To account for solvent outside the particle volume, we combined a static inpainted SSB phase background with a learnable two-dimensional phase layer constrained to zero within the particle supports (Section S10).

On their native evaluation grids (1.2 Å for RELION and 0.72 Å for End2EndPtychoSPA), end-to-end refinement improved the map-to-truth FSC 0.5 resolution from 3.41 to 3.22 Å under the molecular mask and from 3.70 to 3.35 Å under the soft spherical mask of 42 Å radius (Figure 4(d,e)). The median angular pose error decreased from 2.64° to 1.79°, while the median in-plane position error changed little, from 0.38 to 0.40 Å (Figure 4(f,g)).

The improvement persisted under both masks when the maps were compared on either common evaluation grid. We use FSC 0.5 for the primary comparison because some FSC 0.143 crossings lie beyond the ground-truth grid’s 2.4 Å Nyquist limit (Section S8).

Under the molecular mask at FSC 0.5, activating the learnable ice layer only in stage 3 yielded 3.32 Å, compared with 3.22 Å when active throughout refinement. Omitting the ice model yielded 3.57 Å, worse than the 3.41 Å RELION initialization, and left the median orientation error at 2.65°.

From a RELION initialization seeded with the ground-truth model low-pass filtered to 20 Å, refinement improved the molecular-mask FSC 0.5 resolution from 3.09 to 2.96 Å and reduced the median orientation error from 1.62° to 1.16° (Section S8).

Across three minibatch orderings, molecular-mask FSC 0.5 resolutions ranged from 3.21 to 3.22 Å. These runs shared the initial maps and half-set assignments, so they do not assess sensitivity to either (Section S8).

The improvements were confined to local refinement: all 122 inherited pseudo-dyad flips remained after optimization (8.9% of matched particles). All four initial flips also persisted in the 20 Å-start control (0.3% of matched particles).

The particle-subset analysis of the 2D route also shows poorer resolution with **de novo** alignment than with true poses, particularly for smaller subsets (Figure 4(h)). This supports initial orientation recovery as a limitation in this dataset; no End2EndPtychoSPA subset series was run on haemoglobin.

### 3.2. Experimental cryo 4D-STEM benchmarks

We next tested End2EndPtychoSPA on four experimental cryo 4D-STEM datasets using initial maps and poses from the 2D route, with dataset-specific filtering of the starting maps. Because experimental datasets have no known ground-truth potential and may include radiation damage, charging and beam-induced motion, we evaluated the reconstructions against external atomic models and maps, together with half-map diagnostics for high-frequency coupling (Section 2.5). The experimental protocols were pre-registered and fixed before the haemoglobin simulation study, so these reconstructions did not include the learnable ice layer introduced in Section 3.1.

#### 3.2.1. Apoferritin: comparison against the matched 2D route and deposited benchmark

On apoferritin, we compared the final End2EndPtychoSPA reconstruction with our internal 2D-route baseline, run R1, using the same 11,552 particle picks, scan-pure half sets and octahedral (O) symmetry. End2EndPtychoSPA reconstructed items for the 11,150 particles whose scan windows lay inside the scan (Section 2.3.3). We also compared the reconstruction with the deposited EMD-19425 map (Mohammed *et al*., 2024*a*; Küçükoğlu *et al*., 2024*d*) at the same particle count (Table 3).

**Table 3.** Quantitative evaluation of apoferritin reconstructions at 11,552 particles. Map-to-model columns report the 8–30 Å band correlation against PDB 7A4M with (*C*_w_) and without (*C*) radial amplitude whitening, together with the map-to-model FSC at 10 Å and 8 Å and the FSC 0.5 resolution, all evaluated on the half-map mean. Two of these scores exclude the Fourier shells of the scan lattice, and the two lattice steps give different values for them. *C*_w_ uses the nominal 20.0 Å step, as pre-declared; the calibrated 20.6 Å step gives 0.8617, 0.8438, 0.8445 and 0.8455 (rows 1–4), and every sign of the comparison stays the same. Model 0.5 uses the calibrated 20.6 Å step; the nominal step gives 6.90 Å for End2EndPtychoSPA and 7.41 Å for our 2D route. *C*, FSC_10_ and FSC_8_ are identical for both steps. The half-map 0.143 column is evaluated on the full native voxel grid without spatial masking. Benchmark thresholds equal the EMD-19425 half-map mean plus pre-declared margins of 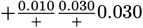, and start thresholds equal our 2D-route start plus the same margins.

| Map | *C*<sub>w</sub> | *C* | FSC<sub>10</sub> | FSC<sub>8</sub> | model 0.5 (Å) | half-map 0.143 (Å) |
|---|---|---|---|---|---|---|
| **End2EndPtychoSPA (this work)** | **0.8706** | **0.852** | **0.835** | **0.710** | **7.06** | **5.99** |
| EMD-19425, half mean | 0.8568 | 0.794 | 0.796 | 0.623 | – | – |
| EMD-19425, deposited map | 0.8576 | 0.820 | 0.840 | 0.674 | – | – |
| 2D route (RELION, ours) | 0.8586 | 0.799 | 0.801 | 0.659 | 7.17 | 6.83 |
| Benchmark threshold | 0.8668 | – | 0.826 | 0.653 | – | – |
| Start threshold | 0.8686 | – | 0.831 | 0.689 | – | – |

Comparing with R1 tests the effect of end-to-end refinement on identical particle picks, whereas comparing with EMD-19425 benchmarks the full workflow against the published map, which uses different particle selection and map post-processing. We scored the mean of the two aligned half maps against the atomic coordinates of PDB 7A4M (Nakane *et al*., 2020) from an independent study. The final refinement schedule and the epoch-selection rule, which selected the final epoch of stage 2, were fixed before training. All benchmark and start-improvement decision thresholds were computed before any End2EndPtychoSPA epoch was scored.

Compared with the matched R1 baseline, End2EndPtychoSPA increased the map-to-model FSC at 8 Å from 0.659 to 0.710 and improved the map-to-model FSC 0.5 resolution from 7.17 to 7.06 Å (Table 3). The reconstruction exceeded all six pre-declared decision thresholds, each defined as a reference score plus a fixed decision margin rather than a statistical confidence interval. Compared with the post-processed EMD-19425 deposited map, End2EndPtychoSPA achieved higher scores in *C*_w_, *C* and FSC_8_ while trailing slightly in FSC_10_ (0.835 versus 0.840). The largest improvement occurred at 8 Å resolution, where FSC_8_ increased from 0.685 at the end of stage 1 to 0.710 at the end of stage 2 (Figure 5(d)).

**Figure 5.**
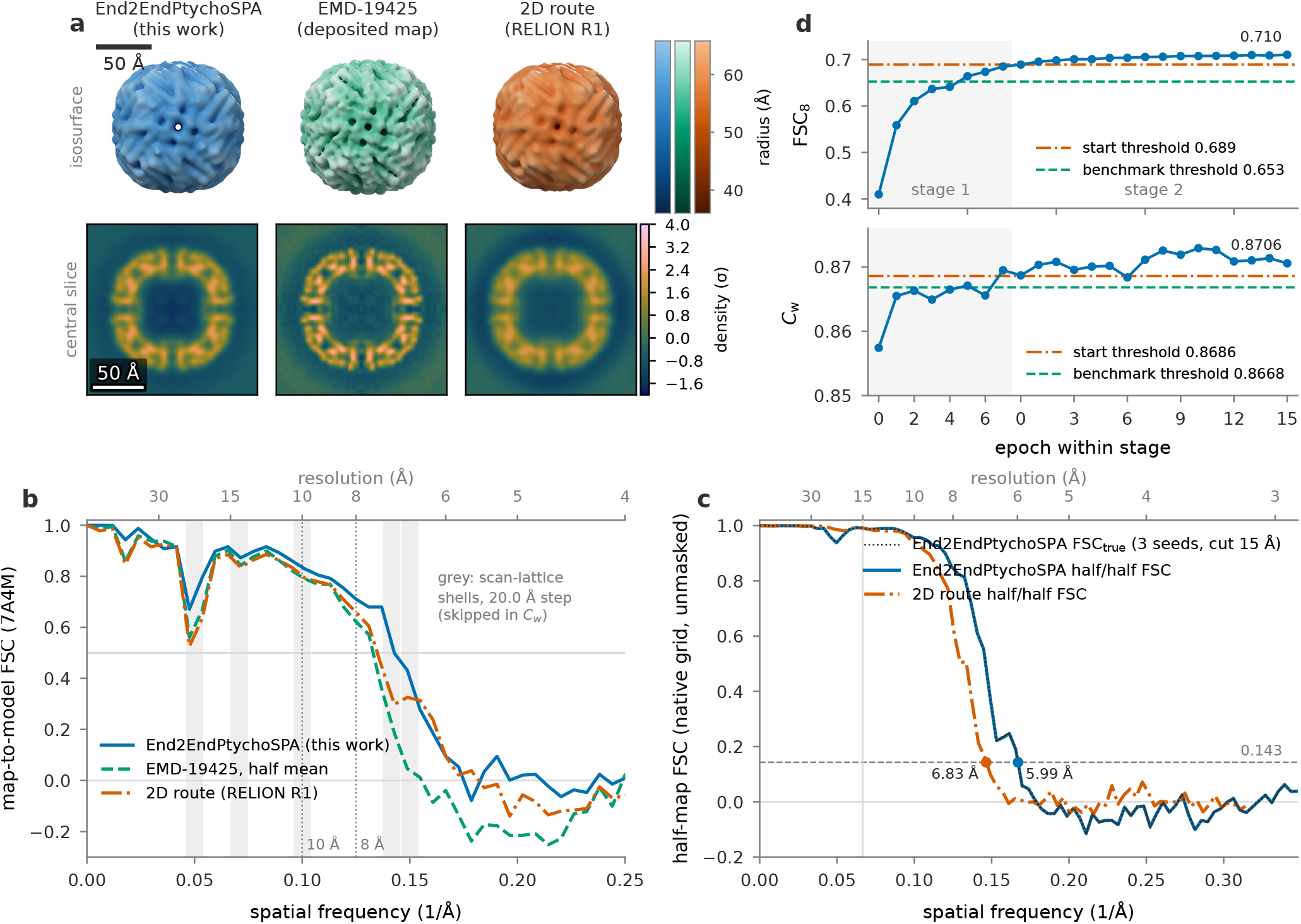
Experimental apoferritin reconstruction at 11,552 particles comparing End2EndPtychoSPA, our matched RELION 2D-route baseline on the same particles (R1) and the deposited EMD-19425 map. (a) Half-map means of End2EndPtychoSPA and our 2D route alongside the deposited EMD-19425 map, each aligned to PDB 7A4M and rendered under identical display filtering (6 Å Gaussian low-pass filter and *z*-score normalization inside the 70 Å evaluation sphere). Top: isosurface contoured at the volume of the 7A4M 24-mer assembly viewed along the four-fold symmetry axis; note that the four-fold channel is resolved as an open pore in the End2EndPtychoSPA map. Bottom: central equatorial slice. Scale bars: 5 nm. (b) Map-to-model FSC against PDB 7A4M within the 70 Å evaluation sphere. The EMD-19425 curve is the half-map mean, not the deposited map of panel (a). Vertical grey bands indicate the reciprocal scanlattice shells of the nominal 20.0 Å step that *C*_w_ excludes (Table 3 gives the 20.6 Å values); vertical dotted lines mark 10 Å and 8 Å, where the benchmark decision thresholds apply. (c) Unmasked, unaligned gold-standard half-map FSC evaluated on the full native voxel grid of each reconstruction. Dotted curves show the phase-randomized corrected curve FSC_true_ for End2EndPtychoSPA beyond 15 Å across three random seeds. (d) Progression of FSC_8_ and *C*_w_ across End2EndPtychoSPA training epochs (stage 1 shaded), plotted against the pre-declared benchmark and start-improvement thresholds. All numerical crossings are listed in Table 3.

We also ran the high-frequency defect diagnostic (Section 2.5) to check for filter coupling between the half sets. The End2EndPtychoSPA apoferritin map passed all three checks: its pooled half-map FSC between 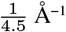and Nyquist was −0.008, and the phase-randomization check on the full native grid gave a first FSC_true_ 0.143 crossing at 5.98–5.99 Å across all three random seeds and all three frequency cutoffs. No high-frequency rebound was detected under these checks. The check randomizes the phases of the completed half maps and therefore does not by itself establish their complete statistical independence (Section 2.5). On the full native voxel grid without a mask, the End2EndPtychoSPA half-map FSC crossed 0.143 at 5.99 Å, compared with 6.83 Å for our matched R1 2D-route baseline evaluated with the same function.

To test whether the optimization schedule and symmetry constraints can produce high-frequency correlations in the absence of macromolecular diffraction signal, we ran a null control using simulated vacuum diffraction patterns paired with the experimental scan windows, particle poses, incident probes and matched detector shot noise (V2b-null, Section S3).

Several controls test the stability and limits of the apoferritin result. The margins by which End2EndPtychoSPA clears the pre-declared *C*_w_ and FSC_10_ thresholds are small. Four replicate runs varied only in minibatch ordering: one used the original order, and three used independent batch seeds with the integrated library code. Across *C*_w_, *C*, FSC_10_ and FSC_8_, the replicate runs had standard deviations of 0.0002–0.0004 and passed all high-frequency diagnostic checks. The minimum gains over our 2D route were +0.012 in *C*_w_ and +0.051 in FSC_8_, and the minimum benchmark margin was +0.0037 in *C*_w_. These replicates show that the result is insensitive to minibatch ordering, but because they share the same scans, particles, half sets and initialization, they do not measure sampling variability across micrographs.

To estimate micrograph-level sampling variability, we ran a five-fold leave-one-fifth-out jackknife across scans, repeating both the RELION 2D-route refinement (including particle selection) and the two-stage End2EndPtychoSPA refinement on each 80% scan subset. The two routes use the same particles in each fold, except that the End2EndPtychoSPA fold tables omit 1–4 particles that lie on scans without probe calibration (of 8,945– 9,460 per fold). In all five jackknife folds, End2EndPtychoSPA improved upon the matched 2D route by +0.011 to +0.014 in *C*_w_, +0.028 to +0.037 in FSC_10_ and +0.043 to +0.062 in FSC_8_. The delete-one-group jackknife standard errors of the gain are 0.003, 0.006 and 0.013, respectively (the library initialization route used in the folds reproduces the full-dataset scores to within 2 × 10^−4^).

Because End2EndPtychoSPA initializes from the 20 Å low-pass-filtered half maps and poses of the 2D route, its gains represent a refinement of the 2D-route solution rather than an **ab initio** reconstruction. We also do not report a spatially masked half-map FSC resolution for End2EndPtychoSPA: under tight solvent masks matched to the deposited workflow, post-hoc masking introduced high-frequency correlations that failed our phase-randomization check, so we report only the unmasked native-grid half-map FSC crossing of 5.99 Å, which is not directly comparable to the masked 5.8 Å value reported for EMD-19425 in cryoSPARC (Küçükoğlu *et al*., 2024*d*).

We also checked whether the apoferritin comparison depends on the docking frame of the atomic reference or on the scan-step calibration. When PDB 7A4M was evaluated directly in its deposited octahedral symmetry frame without preliminary docking into EMD-19425, all four maps scored slightly higher. The margins over the benchmark thresholds increased to +0.007, +0.012 and +0.063. Excluding scan-lattice shells at the calibrated 20.6 Å step rather than 20.0 Å left *C*, FSC_10_ and FSC_8_ unchanged, and combining both changes yielded margins of +0.011, +0.012 and +0.063. These results are reported as secondary checks because a voxel-size metadata discrepancy was detected in the 2D-route map during validation.

Similarly, a 1% discrepancy between the voxel-size metadata used to write and read the initial table maps shifted the best-fit magnification of the End2EndPtychoSPA map against PDB 7A4M to 0.995. This remained within the pre-declared tolerance of ±0.5%. Evaluating each map at its own optimal isotropic scale preserved positive benchmark margins across all three metrics: +0.003, +0.005 and +0.051.

#### 3.2.2. Scaling with particle number on apoferritin

To see how reconstruction quality scales with dataset size, we ran both the 2D route and End2EndPtychoSPA on nested subsets of 1,444, 2,888, 5,776 and 11,552 apoferritin particles (Table S5, Figure 6). Every End2EndPtychoSPA subset reconstruction passed the high-frequency defect diagnostic.

**Figure 6.**
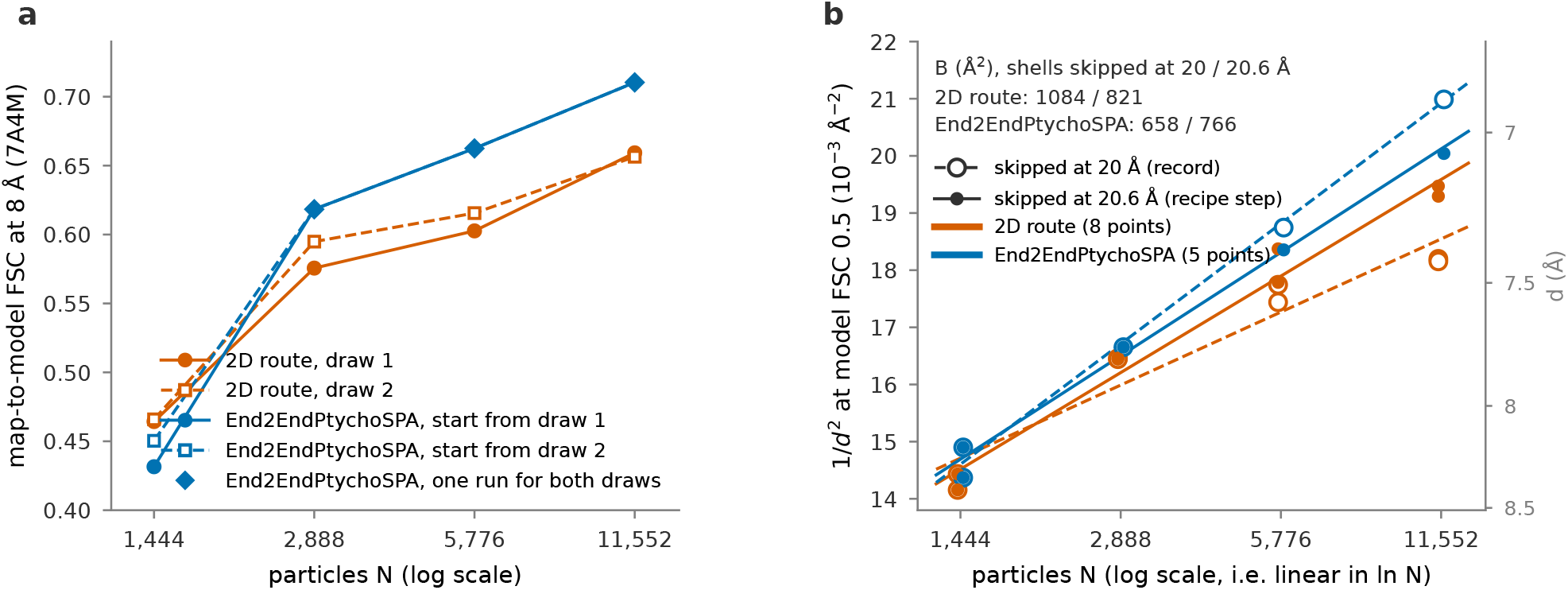
Scaling of apoferritin reconstruction quality across nested subsets of 1,444, 2,888, 5,776 and 11,552 particles evaluated against PDB 7A4M on the half-map mean. The 2D route corresponds to RELION refinements on two independent nested subset draws (circles and squares), while each End2EndPtychoSPA refinement (diamonds) initializes from the corresponding 2D-route solution at the same *N*. End2EndPtychoSPA was run on both draws only at 1,444 particles; at the larger *N*, one End2EndPtychoSPA run (from the first draw) is used for both series, so the two series are not two complete end-to-end repeats. (a) Map-to-model FSC at 8 Å (FSC_8_), which is unaffected by scan-lattice shell exclusion. (b) Rosenthal–Henderson ResLog plot (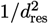 versus ln *N*, where *d*_res_ is the map-to-model FSC 0.5 crossing) evaluated with scan-lattice shells excluded at the nominal 20.0 Å step (open symbols) or at the calibrated 20.6 Å step (filled symbols), together with ordinary least-squares fits *B* = 2/slope (dashed and solid lines). Whereas the *B*-factor difference is 426 Å2 under the 20.0 Å lattice mask, it narrows to 55 Å2 under the 20.6 Å mask (below the pre-declared robustness threshold of 136 Å2), demonstrating that the fitted slope depends on lattice-shell placement whereas the FSC_8_ advantage in (a) does not.

The map-to-model correlation at 8 Å (FSC_8_) does not depend on how the reciprocal scan-lattice shells are masked. At 1,444 particles, End2EndPtychoSPA scored slightly below the 2D route (by −0.02 to −0.03 in FSC_8_). At 2,888, 5,776 and 11,552 particles, End2EndPtychoSPA reached higher FSC_8_ than the 2D route, with gains of +0.04, +0.06 and +0.05 relative to the first 2D subset draw and +0.02, +0.05 and +0.05 relative to the second draw (Figure 6(a)).

We also fitted a pre-registered Rosenthal–Henderson ResLog line (Rosenthal & Henderson, 2003) to the map-to-model FSC 0.5 resolution *d*. When scan-lattice shells were excluded at the nominal 20.0 Å step, the fitted ResLog *B*-factor was 658 Å^2^ for End2EndPtychoSPA compared with 1,084 Å^2^ for the 2D route. However, a pre-registered sensitivity check showed that this *B*-factor difference depends on where the excluded scan-lattice shells are placed, because the FSC 0.5 crossings at large *N* lie next to the 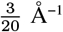 lattice shell. When the lattice shells are excluded at the calibrated 20.6 Å scan step, the fitted *B*-factors shift to 766 Å^2^ and 821 Å^2^ (a difference of 55 Å^2^); under the deposited-frame placement of PDB 7A4M, the difference is 626 Å^2^ at the 20.0 Å lattice setting and 65 Å^2^ at 20.6 Å. Because our pre-registered robustness criterion required the *B*-factor difference to exceed 136 Å^2^ (the combined draw-to-draw spread) in all three variants, the *B*-factor difference is not robust to lattice-shell placement. The fitted End2EndPtychoSPA *B*-factor was lower than that of the 2D route in every variant (and smooth logistic fits to the FSC curves gave differences of 122–160 Å^2^), but because of this sensitivity we do not claim a *B*-factor reduction and report only that End2EndPtychoSPA reached higher FSC_8_ than the matched 2D route at 2,888, 5,776 and 11,552 particles. A secondary ResLog fit to the unmasked native-grid half-map FSC 0.143 crossings gave *B* = 507 ± 112 Å^2^ (95% confidence interval 297–1,723 Å^2^) for End2EndPtychoSPA and 667 ± 77 Å^2^ (521–928 Å^2^) for the 2D route across all points of both draws, with overlapping confidence intervals. In addition, each End2EndPtychoSPA subset is initialized from the corresponding 2D-route half maps and poses at that particle count, so the two series are coupled, and the map-to-model FSC flattens out at large *N*.

#### 3.2.3. Regimes where End2EndPtychoSPA does not yet improve upon the 2D route: T20S, TMV and Phi92

To test how the algorithm performs on lower-symmetry and helical specimens, we ran pre-registered comparisons on three additional experimental datasets: the T20S proteasome, tobacco mosaic virus (TMV) and the Phi92 bacteriophage tail sheath (Table 4, Figure 7). None of these three datasets showed a validated improvement over the external benchmark.

**Table 4.** Summary of pre-registered evaluation outcomes across all four experimental specimens. Outcomes for reconstructions that fail the high-frequency defect diagnostic are marked as not valid. ^a^On Phi92, the benchmark threshold is already exceeded by our cryoSPARC 2D-route baseline and the End2EndPtychoSPA initialization, with no additional signal gained during End2EndPtychoSPA training.

| Specimen | defect gate | adds signal over start | benchmark vs published | note |
|---|---|---|---|---|
| apoferritin (O) | pass | pass | pass | main result |
| T20S (D7) | pass | pass | fail (FSC<sub>8</sub>) | published inputs incomplete |
| TMV (helical) | fail | pass (not valid) | fail (not valid) | behind our 2D route |
| Phi92 (helical, C6) | fail | fail (not valid) | pass<sup>a</sup> (not valid) | 2D route already passes |

**Figure 7.**
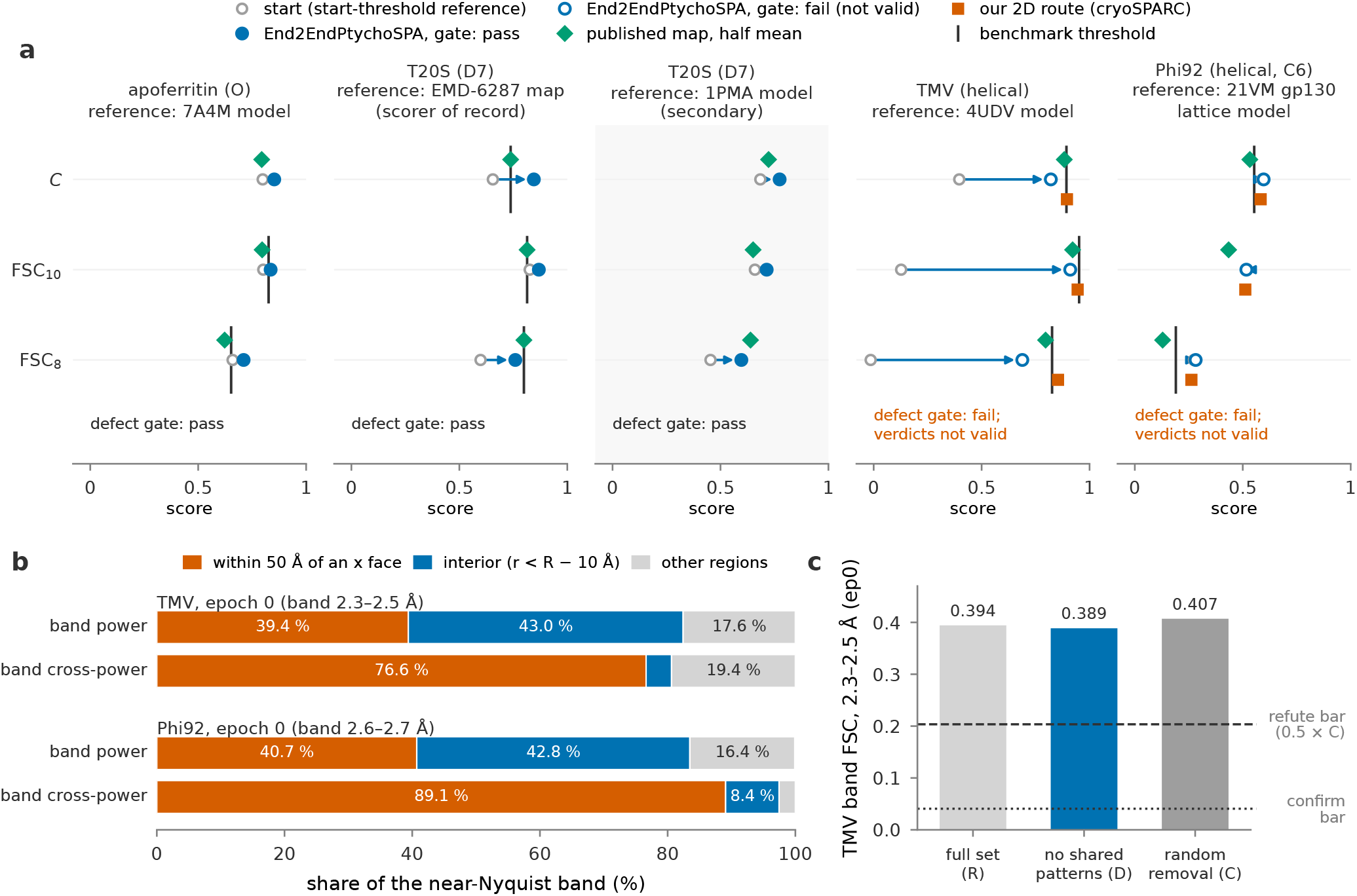
Cross-specimen quantitative summary and helical boundary analysis. (a) Map-to-model metrics (*C*, FSC_10_, FSC_8_) for the low-pass-filtered initialization (grey open circles), the End2EndPtychoSPA reconstruction at its pre-declared epoch (blue circles), the deposited EMDB half-map mean (green diamonds) and, for the helical specimens, our matched cryoSPARC 2D-route refinement on the same segments (vermillion squares; for Phi92, 1,648 of the 1,651 End2EndPtychoSPA segments). Each specimen is evaluated against its own reference (Table 1), so absolute values are comparable only within each column; the shaded panel (T20S scored against PDB 1PMA) is a secondary evaluation without pre-declared thresholds. Note that the TMV initialization scores are low because the starting map is hard low-pass filtered at 15 Å. Horizontal black ticks denote the pre-declared benchmark thresholds (Section 2.5; for T20S, zero margins were set so the tick coincides with the green diamond; for Phi92, the additional reverse-fit requirement is not plotted). Filled blue markers indicate reconstructions that pass the high-frequency defect diagnostic; open blue markers fail the diagnostic (TMV and Phi92). (b) Spatial localization of the shared near-Nyquist frequency band that causes the helical reconstructions to fail the phase-randomization diagnostic (TMV: 0.400–0.435 Å^−1^ Phi92: 0.370–0.385 Å^−1^), evaluated on the band-pass-filtered update at epoch 0. The boundary zones within 50 Å of the two *x* box faces where the continuous helix is truncated contain 39% and 41% of the total band power, but account for 76.6% and 89.1% of the inter-half cross-power. (c) Scan-overlap control on TMV at epoch 0, comparing the high-frequency band FSC of the full segment set (R), a disjoint subset removing every segment whose scan window shares diffraction frames with the opposite half set (D), and a random removal of equal size (C). D remains well above the 0.5 × *C* refutation threshold, showing that cross-half scan-window overlap is not required for the epoch-0 near-Nyquist correlation.

On the 20S proteasome (T20S), the End2EndPtychoSPA reconstruction passed the high-frequency defect diagnostic and improved over its 2D-route initialization across every reference metric (with *C* against the reference map EMD-6287 increasing from 0.655 at initialization to 0.845 after End2EndPtychoSPA refinement). Its initialization is the 2D-route map of all 24,973 particles, whereas End2EndPtychoSPA reconstructs 17,970 items because of a cap of 9,000 items per half set (Section 2.3.3); the start comparison is therefore not particle-matched. However, it fell below the external benchmark threshold against the deposited EMD-67291 half-map mean at 8 Å (FSC_8_ of 0.759 versus 0.799). In our secondary evaluation against the atomic coordinates of PDB 1PMA, End2EndPtychoSPA achieved *C* /FSC_10_ /FSC_8_ values of 0.774 /0.714 /0.597 (model 0.5 crossing at 7.75 Å), compared with 0.723 /0.651 /0.639 (7.24 Å) for the deposited EMD-67291 half maps. Because the particle coordinates used to reconstruct EMD-67291 were not included in the public deposition, a particle-matched comparison is not possible, and we did not optimize T20S further.

On tobacco mosaic virus (TMV), End2EndPtychoSPA improved upon its 15 Å low-pass-filtered initialization, but remained below both the deposited EMD-19885 half-map mean (*C* /FSC_10_ /FSC_8_ of 0.821 /0.911 /0.689 versus 0.883 /0.922 /0.796) and our own cryoSPARC 2D-route refinement on the same 4,436 helical segments (0.895 /0.945 /0.854), and failed the high-frequency phase-randomization diagnostic.

On the Phi92 bacteriophage tail sheath, our cryoSPARC 2D-route refinement on the 1,648 segments already exceeded the deposited EMD-19430 map (Mohammed et al., 2024*c*). Although the End2EndPtychoSPA map exceeded the external benchmark thresholds, its initialization already did so before training, and End2EndPtychoSPA neither improved upon its initialization nor passed the high-frequency phase-randomization diagnostic.

Both helical reconstructions failed the phase-randomization diagnostic even with the blur-last gradient hook because of a narrow shared spatial-frequency band just below the Nyquist frequency (2.3–2.5 Å in TMV and 2.6–2.7 Å in Phi92). Filtering the epoch-0 update to this near-Nyquist band showed that 77% (TMV) and 89% (Phi92) of the shared inter-half cross-power lies within 50 Å of the two axial *x* faces where the reconstruction box truncates the continuous helical rod (Figure 7(b)). Every refinement epoch using the Gaussian gradient filter regenerated this face-localized band, whereas replacing the Gaussian filter with a hard spectral cutoff that zeroes the band removed it. We ran three pre-registered diagnostic tests to find the source of this boundary band (Supporting Information S4) and found that zero-padding of the FFT blur, widening of the blur margin and trilinear interpolation during screw averaging do not produce the artefact. In a linear forward model, truncating a continuous helix at the box boundaries did produce a face-localized shared band, but at an amplitude 10^3^–10^4^ times weaker than in the full refinement. In the TMV control at epoch 0, removing every helical segment whose scan window shares diffraction patterns with the opposite half set left the near-Nyquist band correlation almost unchanged (0.389 versus 0.407 for a random removal of the same number of segments; Figure 7(c)), showing that cross-half scan-window overlap is not required for this epoch-0 correlation. The exact coupling mechanism at the helical box boundaries remains unresolved (Section 4).

Finally, the stored TMV and T20S End2EndPtychoSPA volumes and their initializations have inverted axial handedness (*z*-mirror); because our evaluation pipeline searches both hands before computing map-to-model metrics, none of the scores in this work are affected.

## 4. Discussion

Our multislice simulations separate initial particle alignment from 3D potential reconstruction, which cannot be separated in experimental data alone. On simulated myoglobin (17.9 kDa), the 2D ptychographic phase micrographs preserve structural information up to the Nyquist frequency when reconstructed with the ground-truth poses, but global orientation search from a low-pass filtered known-structure reference fails in the 2D route, and our local gradient-based refinement in End2EndPtychoSPA does not recover the missed orientations (Section S9). For small macromolecules under these imaging conditions, pose determination is therefore the bottleneck; while higher electron dose improves alignment in simulation and the two learned 2D image priors we tested degraded orientation accuracy, global or orientation-marginalized 4D-STEM pose search remains to be developed.

On simulated haemoglobin (64.5 kDa), the 2D route recovers initial particle orientations. Here, End2EndPtychoSPA refines the angular poses and improves the 3D resolution against the ground-truth potential relative to RELION under the same evaluation mask, provided that the forward model includes a back-ground layer to model the surrounding vitreous ice (Section 3.1). The resolution gain of 0.19 Å at FSC 0.5 exceeds the spread across independent minibatch orderings (0.012 Å), although we have not tested variability across independent initial models or half-set splits, and local pose refinement retains the pseudo-dyad orientation flips from the 2D route. Unmodelled phase shifts from the vitreous-ice solvent limit end-to-end 3D refinement when particles are modelled in vacuum, an effect not yet included in our pre-registered experimental protocol.

On the experimental apoferritin dataset (11,552 particles), End2EndPtychoSPA improves agreement with an external atomic model (PDB 7A4M) relative to both our matched RELION 2D-route baseline on the same particles and the deposited EMD-19425 half-map mean (Section 3.2.1). The reconstruction passes our high-frequency phase-randomization diagnostic and shifts the unmasked native-grid half-map FSC 0.143 crossing by about 0.8 Å relative to our matched 2D route (from 6.83 Å to 5.99 Å). Across nested particle subsets, the two workflows perform similarly at 1,444 particles, while End2EndPtychoSPA reaches higher 8 Å map-to-model correlation (FSC_8_) from 2,888 particles onward (Section 3.2.2), indicating that larger experimental datasets benefit once initial alignments are established. We do not claim a reduction in Rosenthal–Henderson *B*-factor because our sensitivity analysis showed that the fitted *B*-factor difference varies from 55 Å^2^ to 626 Å^2^ depending on whether the excluded reciprocal scan-lattice shells are placed at 20.6 Å or 20.0 Å and how the reference model is docked.

Single-point FSC crossings near reciprocal scan-lattice reflections are sensitive to lattice masking in scanned diffractive imaging, so we recommend evaluating reconstructions with map-to-model metrics integrated over a spatial-frequency band and using non-periodic scan geometries experimentally. Evaluated on the unmasked half-map FSC 0.143 crossing, both of our workflows give ResLog *B*-factors of 500–670 Å^2^. For comparison, conventional plane-wave cryo-EM of the same apoferritin preparation achieved a ResLog *B*-factor of 24.5 Å^2^ from 411,705 particles at 1.09 Å resolution, and the published ptychographic map reported a Guinier *B*-factor of 642 Å^2^ ((Küçükoğlu *et al*., 2024*d*), Supplementary Table 1). Both the 2D route and End2EndPtychoSPA thus show a steep high-frequency signal fall-off on this dataset; how much of this fall-off comes from the experimental acquisition parameters (electron dose and 20 Å scan step) versus the reconstruction algorithms remains open.

Our experimental results also show the current limits of the method. Because End2EndPtychoSPA starts from the 20 Å low-pass-filtered half maps and poses of the 2D route, it serves as a 3D refinement step rather than a replacement for the 2D ptycho-graphic pipeline. While the gains on apoferritin are stable across minibatch orderings and remain positive across all five scan-level jackknife folds, the margins over the pre-declared *C*_w_ and FSC_10_ thresholds are small, and testing whether the algorithm improves on the 2D route across other globular proteins will require a second non-helical experimental dataset with publicly deposited particle coordinates, poses and half-set assignments.

On TMV and Phi92, bounded-box gradient updates on continuous helical specimens left a narrow shared correlation band just below the Nyquist frequency within 50 Å of the two axial box faces that truncate the helix (Section 3.2.3). Our three diagnostic tests (Supporting Information S4) ruled out the zero-padding of the FFT blur, the width of the blur margin, screw-symmetry interpolation and shared diffraction frames between overlapping scan windows as the cause, while a linear forward-model truncation test produced a face-localized shared band at the same axial position but with an amplitude three to four orders of magnitude weaker than in the full refinement. Possible causes that remain to be tested include multislice propagation or probe-boundary interactions at the axial cut faces, growing pose or defocus errors near the segment ends, and non-linear amplification during iterative gradient descent. Extending the 3D reconstruction box well beyond the illuminated scan field of every helical segment so that the axial cut faces do not intersect the probe footprint should avoid this boundary effect in future work; until this boundary coupling is resolved, we do not use our helical reconstructions as validated benchmarks.

The order of the gradient filters is a validation risk for iterative, gradient-based three-dimensional refinement (Section 2.4). When a real-space support mask multiplies a low-pass filtered update, the mask convolves the low-frequency update, which the two gold-standard halves share, into high-frequency shells beyond the low-pass cutoff. This ringing lies at the mask boundaries in the solvent region. Map-to-model scores inside the molecular support therefore do not show it, but the half-map FSC counts it as high-resolution signal. Any iterative method that combines spatial-frequency filtering with real-space masking of gradients, density updates or regularizer outputs can have this problem. Two inexpensive checks can detect it: monitoring the pooled half-map FSC in the frequency band beyond the gradient low-pass cutoff, and performing high-frequency phase randomization on the full, unmasked native voxel grid (since resampling and cropping onto a smaller evaluation grid can introduce its own boundary correlations). We have not measured how sensitive these checks are to weaker coupling, and passing them does not show that the half maps are independent.

Two ablations on apoferritin separate the contributions of pose refinement and multislice propagation. We repeated the End2EndPtychoSPA refinement twice with one recipe change each; data, start, half sets and batch order were unchanged. With the particle poses fixed at their 2D-route (R1) values, the map lost 44% of the *C*_w_ gain over R1 and 76% of the FSC_8_ gain, and the half-map FSC crossed 0.143 at 6.93 Å instead of 5.99 Å (R1: 6.83 Å). With one slice (the projection approximation) instead of six, the four map-to-model correlations changed by less than three standard deviations of four batch-order replicates. The model-map and half-map resolutions changed by 0.011 Å and 0.005 Å, just above that bar but 2% and 1% of the gain. At the present resolution of about 6 Å, pose refinement thus carries most of the gain, and multislice propagation carries no measurable part of it. We expect the multi-slice modeling to become more important as high resolution and larger samples are targeted. The per-item defocus offsets took up most of the axial shift of the single projection plane (82 Å of 101 Å).

For future work, we plan to test End2EndPtychoSPA on a second globular macromolecule with deposited particle coordinates, extend the axial reconstruction box for helical specimens, and include the learnable vitreous-ice background layer from our haemoglobin simulations in experimental reconstructions. Ablations of the remaining forward-model terms, the per-window defocus offsets and the overlapping neighbour particles, are also needed. Dynamical scattering simulations of TMV at 300 kV indicate that while single-slice models break down beyond 2.5 Å, propagation through slices thinner than 6 nm preserves structural information to 1.0 Å (Leidl et al., 2023). Our six slices are 4.0 nm thick, below that bound. Reaching higher resolution on future high-dose or counting datasets will require a finer voxel sampling than the 1.438 Å grid (Nyquist limit 2.88 Å) used here and tests of slice-thickness convergence; we have not tested which other factors, such as dose, pose accuracy or the forward-model terms, would then limit the resolution.

## 5. Conclusions

We have introduced End2EndPtychoSPA, an end-to-end multislice reconstruction algorithm that jointly refines a shared 3D electrostatic potential volume, per-particle poses and per-window defocus offsets directly from cryo-4D-STEM diffraction data. In multislice simulations, local gradient-based refinement does not recover particle orientations missed by the 2D ptychographic route on a 17.9 kDa protein, whereas combining End2EndPtychoSPA with a vitreous-ice background model refines the initial 2D-route poses on a 64.5 kDa haemoglobin dataset and improves the 3D resolution against the ground truth relative to RELION. On experimental apoferritin data (11,552 particles), End2EndPtychoSPA improves agreement with an external atomic model over both our matched 2D route on the same particles and the deposited EMD-19425 half-map mean, and scores higher than our 2D route from 2,888 particles onward. Replicate runs across independent minibatch orderings change the map-to-model metrics by less than 5 × 10^−4^ and leave-one-fifth-out scan jackknifing gives positive gains in all five folds, while a failed vacuum null control at discrete lattice-adjacent shells and the lack of validated benchmark gains on T20S, TMV and Phi92 show the current limits of the algorithm. Finally, we showed that applying a spatial support mask after a low-pass filter in a gradient hook couples gold-standard half maps at high spatial frequencies, and we used a native-grid phase-randomization check to detect this coupling in our runs; its sensitivity to weaker coupling remains to be measured.

## 6. Acknowledgements

During this work, the authors used the AI coding assistants Claude Code (Anthropic) and Codex (OpenAI) to write and test analysis and reconstruction code, to run and track computations, to prepare figures and to edit the manuscript text. The authors checked all code, results and text and take full responsibility for the content of this article. The authors gratefully acknowledge the scientific support and HPC resources provided by the Erlangen National High Performance Computing Center (NHR@FAU) of the Friedrich-Alexander-Universität Erlangen-Nürnberg (FAU). The hardware is partially funded by the German Research Foundation (DFG). The authors gratefully acknowledge the Gauss Centre for Supercomputing e.V. (www.gauss-centre.eu) for funding this project by providing computing time on the GCS Sovereign Supercomputer JUPITER Booster at Jülich Supercomputing Centre (JSC).

## 8. Data and code availability

The 4D-STEM data sets are publicly available in EMPIAR: apoferritin (EMPIAR-12236 (Küçükoğlu *et al*., 2024*a*)), TMV (EMPIAR-12234 (Küçükoğlu *et al*., 2024*b*)), Phi92 (EMPIAR-12233 (Küçükoğlu *et al*., 2024*c*)) and T20S (EMPIAR-13137, session 2 (Li et al., 2025*a*)). The published reference maps used for comparison and scoring are available from the EMDB: apoferritin EMD-19425 (Mohammed et al., 2024*a*), TMV EMD-19885 (Mohammed et al., 2024*b*), Phi92 EMD-19430 (Mohammed et al., 2024*c*), and T20S EMD-67291 (Li et al., 2025*b*) and EMD-6287 (Campbell et al., 2015). The atomic models used by the map-to-model scores are available from the PDB: 7A4M (Nakane et al., 2020) (apoferritin), 4UDV (Fromm et al., 2014) (TMV), 1PMA (Loewe et al., 1996) (T20S) and 21VM (Chen & Liu, 2026) (Phi92). The particle and segment tables, the refined poses, run configurations and score files of the runs of record, the End2EndPtychoSPA half maps of T20S, TMV and Phi92, the simulation results and the pre-registration record are available from Zenodo (doi:10.5281/zenodo.23814850). The atom coordinates of the simulated haemoglobin and myoglobin samples and their ground-truth object phases are available from Zenodo (doi:10.5281/zenodo.23814872). The code will be released upon publication in the open-source package scatterem.

