## Supplementary info for "End-to-end reconstruction for cryo electron ptychographic single-particle analysis"

Draft, 31 October 2026


### S1. Forward model and optimization equations

#### S1.1. Notation and array conventions

Table S1 lists the symbols and units. This section gives the implementation details that the Methods section does not state; it does not repeat the equations of the Methods section.

Throughout this section, the incident electron beam propagates along the positive $z$ axis. Three-dimensional volumes are stored in $zyx$ axis order and two-dimensional images in $yx$

**Table S1**
Symbols and units. Where End2EndPtychoTomo (You *et al.*, 2025) and End2EndPtychoSPA agree, we use the operator notation of You *et al.* (2025). Values in parentheses are those of the apoferritin run of record.

| Symbol | Meaning (units; apoferritin value) |
|---|---|
| $s, i, p, k, j$ | indices of scan, item (particle-centred scan window), particle, probe position and slice |
| $l$, $\boldsymbol{\Theta}_l$, $\boldsymbol{\tau}_l$ | tilt index, tilt angles and tilt translation of End2EndPtychoTomo |
| $d_k$, $\Delta k$, $N$ | native detector pixel (0.00261823 Å$^{-1}$); model detector pixel ($1.66d_k$); model grid ($N = 160$ px) |
| $a$ | object pixel, $\frac{1}{N\Delta k}$ (1.438 Å) |
| $\Delta r$, $\theta_s$, $R(\theta_s)$ | scan step (20.6 Å); scan–detector rotation of scan $s$; its rotation matrix |
| $\boldsymbol{t}_k$, $\boldsymbol{\rho}_k$, $\boldsymbol{n}_k$, $\boldsymbol{\varepsilon}_k$ | integer raster position of probe position $k$; probe position, its integer crop corner and its fractional part (object pixels, $yx$) |
| $\boldsymbol{r}$, $\boldsymbol{q}$, $q$ | real-space coordinate; reciprocal-space coordinate; $q = \lvert\boldsymbol{q}\rvert$ |
| $\lambda$, $k_0$, $\alpha$ | electron wavelength at 300 kV (Å); $k_0 = 2\frac{\pi}{\lambda}$; scattering semi-angle $\lambda q$ |
| $A_s$, $\chi_s$, $Z_n$ | measured aperture amplitude; aberration phase (rad); aberration polynomials of radial order $n$ |
| $\psi$, $\psi_i$ | probe; probe of item $i$, scaled to its measured counts |
| $V$, $\mathfrak{R}(V)$, $\mathfrak{I}(V)$, $\kappa$ | complex volume (a phase per voxel, $\sigma = 1$); phase shift; amplitude attenuation; coupling $\mathfrak{I}(V) = \kappa\mathfrak{R}(V)$ ($\kappa = 0.07$) |
| $V^0$ | initial volume of a half set (2D route) |
| $\boldsymbol{\Theta}_p$, $R_{\boldsymbol{\Theta}_p}$, $Q_p$, $\boldsymbol{\tau}_p$ | Euler angles of particle $p$; its rotation; sampling matrix $R^T_{\boldsymbol{\Theta}_p}$; lateral position (object pixels) |
| $Q^{\mathrm{a}}_p$, $\boldsymbol{\tau}^{\mathrm{a}}_p$ | anchor pose of particle $p$ (2D route) |
| $\boldsymbol{\tau}_{p,i}$, $\mathcal{N}_i$ | lateral offset of particle $p$ from the centre of item $i$; particle copies in item $i$ (its own particle first, at most 16) |
| $A_{\boldsymbol{\Theta},\boldsymbol{\tau}}$, $V_i$ | affine operator (trilinear inverse warp of $V$); sum of the copies in item $i$ |
| $B_b$, $b$, $T$ | binning along $z$; fine planes per slice (28); number of slices (6) |
| $\Delta z$, $V_{i,j}$, $\mathcal{T}$ | slice distance ($28a = 40.26$ Å); slice $j$ of $B_b V_i$; transmission operator |
| $\boldsymbol{C}_{\boldsymbol{\rho}_k}$, $S_{\boldsymbol{\varepsilon}_k}$ | cut of the probe-sized patch at $\boldsymbol{n}_k$; sub-pixel probe shift |
| $\mathcal{P}^d$, $h(\boldsymbol{q})$, $d_i$ | angular-spectrum propagator over $d$ (Å); its phase per Å; free path (defocus offset) of item $i$ (Å) |
| $\mathcal{M}$, $\mathcal{F}$ | multislice operator; unitary 2D Fourier transform on the model detector grid |
| $D_{\boldsymbol{\delta}_{i,k}}$, $\boldsymbol{\delta}_{i,k}$ | descan phase ramp; sub-pixel descan residual (model detector pixels) |
| $\mathcal{A}^{i,k}$, $I$, $\hat{I}$, $v$ | forward model of pattern $k$ in item $i$; gain-scaled counts on the model detector; model intensity; valid-pixel mask |
| $\mathcal{L}$, $P_{\mathrm{pose}}$, $\sigma_R$, $\sigma_\tau$ | data loss; pose prior; prior widths (2°; 3.3 object pixels) |
| $w$, $\gamma_1$, $\Gamma$ | weight of the defocus anchor ($6 \times 10^{-12}$); volume step size (10); item order of an epoch |
| $g$, $H$, $\mathcal{Z}$, $\sigma_{\mathcal{Z}}$ | volume gradient of an item; gradient hook; axial smoothing of the slice gradient; its width (11 planes) |
| $m$, $r_0$, $\mathcal{S}$ | spherical support mask; its radius (72 Å); symmetry gather |
| $G$, $\sigma_G$ | Gaussian blur of the hook; its width (4 Å) |
| $\mathcal{H}$, $n_h$, $\omega_{n_h}$ | screw-symmetry average (helical specimens); screw copy; its Fejér weight |
| $\mu$, $\varepsilon$ | weight of the $L_1$ penalty and small constant of End2EndPtychoTomo (Equation (3)) |

order; because three-dimensional rotation matrices act on Cartesian column vectors in $(x, y, z)^T$ order, the volume sampler permutes coordinates between the two conventions. All Fourier-space arrays place the zero-frequency (DC) component at index $(0, 0)$, and the two-dimensional discrete Fourier transform $\mathcal{F}$ is normalized to be unitary on the $160 \times 160$ model detector grid.

#### S1.2. Resampling of diffraction patterns onto the model detector grid

To compare the measured diffraction patterns with the forward model on a common reciprocal-space grid, we map each native detector frame onto the model detector grid at data loading. Each raw frame is cropped to a $174 \times 174$ native-pixel window with top-left origin $\mathrm{round}(\boldsymbol{c}) - 87$, where $\boldsymbol{c}$ is the fitted bright-field centre of mass on the descan plane; this integer crop tracks the coarse beam descan shift across the scan in whole native pixels. After dividing the raw detector counts by the empirical scan noise gain $g_s$ from the single-sideband fit, we integrate the intensities onto coarser model detector pixels of pitch $\Delta k = 1.66 d_k$. Each model pixel $j$ (indexed relative to the zero-frequency origin along each axis) sums native pixels $i$ weighted by their one-dimensional interval overlap:

$$I_j = \sum_i w_{ji} X_i, \quad w_{ji} = \left| \left[ j\Delta k - \Delta\frac{k}{2}, j\Delta k + \Delta\frac{k}{2} \right] \cap \left[ id_k - \frac{d_k}{2}, id_k + \frac{d_k}{2} \right] \right| \frac{1}{d_k}. \tag{S1}$$

We keep only the model pixels whose intervals lie completely inside the 174-pixel native crop, corresponding to integer offsets $-52, \ldots, 51$ ($104 \times 104$ model pixels), and zero-pad this block to the $N = 160$ pixel model grid. The resampled data are stored as amplitudes $\sqrt{I}$; the Poisson data loss squares them back to intensities.

The binary valid-pixel mask $v$ is formed as the product of two indicator masks: the first retains only those model pixels whose contributing native pixels all fall within the physical detector sensor after the integer descan shift, and the second restricts evaluation to a circular pupil disk of radius 49 model pixels around the zero frequency.

#### S1.3. Scan geometry and particle-centred reconstruction items

Each reconstruction item $i$ consists of the $12 \times 12$ subarray of scan positions whose integer starting index on the scan raster is $\boldsymbol{w}_i = \mathrm{round}(\boldsymbol{s}_p) - 6$, where $\boldsymbol{s}_p$ is the fractional scan-grid coordinate of the central picked particle $p$. The probe positions $\boldsymbol{\rho}_k$ follow Equation (5) (column-vector convention; the implementation evaluates the equivalent product with row vectors). Denoting the lateral dimensions of the item object field by $\boldsymbol{f}$ and defining the centering offset $\boldsymbol{h} = \frac{\boldsymbol{f} - (N, N)}{2}$, we decompose each scan position into an integer pixel corner and a fractional sub-pixel remainder:

$$\boldsymbol{n}_k = \mathrm{round}(\boldsymbol{\rho}_k + \boldsymbol{h}), \quad \boldsymbol{\varepsilon}_k = \boldsymbol{\rho}_k + \boldsymbol{h} - \boldsymbol{n}_k. \tag{S2}$$

Here, $\boldsymbol{n}_k$ specifies the top-left corner of the $N \times N$ object patch extracted by the cropping operator $\boldsymbol{C}_{\boldsymbol{\rho}_k}$, while $\boldsymbol{\varepsilon}_k$ is the residual sub-pixel shift applied to the incident probe wavefunction.

#### S1.4. Incident probe synthesis and normalization

We construct the incident probe wavefunction for each scan $s$ from its measured vacuum aperture amplitude $A_s$, which is cropped and area-resampled onto the model detector grid in the same manner as the diffraction frames. Parameterizing reciprocal-space polar coordinates by the scattering semi-angle $\alpha = \lambda\,|\boldsymbol{q}|$ and azimuthal angle $\varphi_q = \mathrm{atan2}(q_y, q_x)$, the aberration polynomials $Z_n$ of Equation (7) are

$$Z_2 = C_{10} + C_{12a}\cos 2\varphi_q + C_{12b}\sin 2\varphi_q, \tag{S3}$$

$$Z_3 = C_{21a}\cos\varphi_q + C_{21b}\sin\varphi_q + C_{23a}\cos 3\varphi_q + C_{23b}\sin 3\varphi_q, \tag{S4}$$

$$Z_4 = C_{30} + C_{32a}\cos 2\varphi_q + C_{32b}\sin 2\varphi_q + C_{34a}\cos 4\varphi_q + C_{34b}\sin 4\varphi_q, \tag{S5}$$

and the probe of Equation (7) is evaluated as

$$\psi_s^0 = \mathcal{F}^{-1}[A_s(\boldsymbol{q})\exp(-i\chi_s(\boldsymbol{q}))J(\boldsymbol{q})], \quad J_{u,v} = -(-1)^{u+v}. \tag{S6}$$

Here, the checkerboard phase factor $J_{u,v}$ shifts the real-space probe origin to the centre of the $N \times N$ array (with the overall minus sign corresponding to a global phase). The 12 aberration coefficients (defocus $C_{10}$, two-fold astigmatism $C_{12a,b}$, axial coma $C_{21a,b}$, three-fold astigmatism/trefoil $C_{23a,b}$, spherical aberration $C_{30}$, star aberration $C_{32a,b}$ and four-fold astigmatism $C_{34a,b}$, all in Å) are taken from the scan-level 2D ptychographic refinement (where the library convention defines $-C_{10}$ as positive underfocus). We evaluate the aperture wavefunction on the $104 \times 104$ covered block and embed it into the $160 \times 160$ model grid by zero-padding in Fourier space, which preserves total intensity under unitary Fourier normalization.

The item normalization of Section 2.3.4 reads, with the 49-pixel bright-field disk $D$,

$$\psi_i = \psi_s^0 \sqrt{\frac{N_i}{\sum_{\boldsymbol{u}} |\psi_s^0(\boldsymbol{u})|^2}}, \quad N_i = \sum_{\boldsymbol{q}\in D} \left\langle I_{k,\boldsymbol{q}} \right\rangle_k. \tag{S7}$$

#### S1.5. Shared volume, pose parameterization and neighbour copies

We enforce the coupling $\mathfrak{I}(V) = \kappa\mathfrak{R}(V)$ (Section 2.3.5) by projecting $\mathfrak{I}(V) \leftarrow \kappa\mathfrak{R}(V)$ at initialization and after every mini-batch update, and by mapping each complex volume gradient $g$ to $(\mathfrak{R}(g) + \kappa\mathfrak{I}(g))(1 + i\kappa)$ so that gradient descent steps remain on the coupled subspace. Under our sign convention (Equation (S11)), a positive imaginary part $\mathfrak{I}(V) > 0$ attenuates the transmitted wave amplitude, and the interaction constant $\sigma$ of You *et al.* is absorbed into $V$ ($\sigma = 1$).

Each particle orientation is parameterized by three Euler angles $\boldsymbol{\Theta}_p = (\varphi_p, \theta_p, \psi_p)$ in the convention of the volume sampler: $\varphi_p$ rotates about $z$, $\theta_p$ about $y$ and $\psi_p$ about $x$, and they define the inverse sampling rotation matrix

$$Q_p = R_x(\psi_p)R_y(\theta_p)R_z(\varphi_p) = R^T_{\boldsymbol{\Theta}_p}. \tag{S8}$$

These angles are not the ZYZ Euler angles of RELION. The particle table stores each RELION orientation as a rotation matrix, and the sampler angles are extracted from that matrix. When the `--mirror-hand` flag is active, each input rotation matrix from the particle table is conjugated by the reflection matrix $H = \mathrm{diag}(1, 1, -1)$ while the initial volume $V^0$ is inverted along the beam axis $z$, leaving all two-dimensional projections invariant. Denoting the geometric centre of the volume grid by $\boldsymbol{c} = \left(\frac{N_x - 1}{2}, \frac{N_y - 1}{2}, \frac{N_z - 1}{2}\right)$, the affine sampling operator for particle copy $p$ within item $i$ evaluates $V$ at transformed coordinates

$$\boldsymbol{\xi}_{p,i}(\boldsymbol{x}) = \boldsymbol{c} + Q_p\left(\boldsymbol{x} - \boldsymbol{c} + (\tau_x, \tau_y, 0)^T\right), \tag{S9}$$

via separable trilinear interpolation with zero boundary conditions outside the volume box:

$$\left(A_{\boldsymbol{\Theta}_p, \boldsymbol{\tau}_{p,i}} V\right)(\boldsymbol{x}) = \sum_{\boldsymbol{n}\in\mathbb{Z}^3} V[\boldsymbol{n}] \prod_{a\in\{x,y,z\}} \max(0, 1 - |\xi_{p,i,a} - n_a|). \tag{S10}$$

The real and imaginary components of $V$ are interpolated identically without a Jacobian determinant factor. Here, $\boldsymbol{\tau}_{p,i} = \boldsymbol{\tau}_p - \boldsymbol{o}_i$ denotes the in-plane offset between the particle centre $\boldsymbol{\tau}_p$ and the item centre $\boldsymbol{o}_i$ expressed in object pixels, and individual copies within an item share a common axial plane ($z$-offset of zero).

The neighbour copy list $\mathcal{N}_i$ of Equation (8) is built once before training from the picked particles of the same scan and half set, and it stays fixed during refinement.

#### S1.6. Axial slice binning and phase-grating transmission

The fine grid of item $i$ has $N_z = \left\lceil 241\frac{\text{Å}}{a}\right\rceil = 168$ axial planes. With the complex volume, the binning and transmission of Equation (10) read

$$V_{i,j} = [B_b V_i]_j = \sum_{z\in Z_j} V_i(z,y,x), \quad \mathcal{T}^{V_{i,j}}\psi = \psi\cdot\exp(-\Im(V_{i,j}) + i\Re(V_{i,j})). \quad \text{(S11)}$$

The axial smoothing $\mathcal{Z}$ of Equation (13) is a custom autograd operator immediately upstream of $B_b$.

#### S1.7. Angular-spectrum propagation and per-item defocus offset

For evanescent components ($k_0^2 \leq (2\pi)^2 q^2$), the propagator of Equation (9) uses $h(\boldsymbol{q}) = -k_0$. The sub-pixel probe shift operator is evaluated in Fourier space via the linear phase ramp

$$S_{\boldsymbol{\varepsilon}}\psi = \mathcal{F}^{-1}[\exp(-2\pi i\boldsymbol{\nu}\cdot\boldsymbol{\varepsilon})\mathcal{F}\psi], \quad \text{(S12)}$$

where $\boldsymbol{\nu}$ denotes spatial frequency in cycles per object pixel, and $\boldsymbol{C}_{\boldsymbol{\rho}_k}$ extracts the $N\times N$ patch of each slice at the integer corner $\boldsymbol{n}_k$. We write $\psi^{(6)}$ for the exit wave of Equation (10).

#### S1.8. Far-field detector intensity prediction

Because the native detector frames are cropped at integer pixel coordinates, a fractional sub-pixel descan residual $\boldsymbol{\delta}_{i,k}$ remains on each frame. We account for this sub-pixel shift by multiplying the real-space exit wave by a two-dimensional linear phase ramp prior to Fourier transformation:

$$D_{\boldsymbol{\delta}}\psi(u,v) = \psi(u,v)\exp\left[2\pi i\frac{\delta_y u + \delta_x v}{N}\right], \quad \text{(S13)}$$

which yields the predicted far-field detector intensity

$$\hat{I} = |\mathcal{F}D_{\boldsymbol{\delta}_{i,k}}\psi^{(6)}|^2 = |\mathcal{A}^{i,k}_{\psi,\boldsymbol{\rho},\Theta,\boldsymbol{\tau}}(V)|^2, \quad \mu = \hat{I} + 10^{-10}. \quad \text{(S14)}$$

A forward hook then crops the $160\times160$ predicted intensity array to the $104\times104$ covered detector block. The forward model uses a single coherent probe mode and does not include a detector point-spread function or a fitted per-frame intensity scale factor.

#### S1.9. Poisson loss implementation and rotational gauge

The data loss of Equation (11) is PyTorch's `poisson_nll_loss` with `log_input=False` and `full=False` (which omits the constant Stirling log-factorial term); its denominator is bounded below by one. To fix the global rotational gauge of the reconstruction, the orientation gradient of the first particle in the table is zeroed.

#### S1.10. Optimizers and signed-gradient defocus updates

The SGD updates of the volume and the poses (Section 2.4) use no momentum, weight decay or learning-rate schedule. The per-item defocus offsets $d_i$ are updated using `MaskedAdam` configured with zero first- and second-moment decay factors ($\beta_1 = \beta_2 = 0$) and numerical stabilizer $\varepsilon = 10^{-30}$. Denoting the raw defocus derivative by $g_i = \partial\frac{\mathcal{L}}{\partial}d_i$, a gradient hook forms the regularized update

$$\tilde{g}_i = \begin{cases} g_i + 2wd_i & \text{if } g_i \neq 0 \\ 0 & \text{if } g_i = 0 \end{cases}, \quad d_i \leftarrow d_i - \gamma_d\cdot\frac{\tilde{g}_i}{|\tilde{g}_i| + 10^{-30}}, \quad \text{(S15)}$$

a step of $\gamma_d$ along the sign of $\tilde{g}_i$ at each visit of item $i$; the lateral free-path tilt gradients are masked to zero.

#### S1.11. Point-group and helical volume gradient hooks

Let $g$ denote the complex volume gradient extracted on the central cubic subarray of side length $n = \min(N_z, N_y, N_x) = 168$ voxels immediately after the axial smoothing filter $\mathcal{Z}$ and the amplitude-coupling projection. The spherical support mask $m(r)$ is defined as a function of radial distance $r$ from the geometric centre of the cube:

$$m(r) = \begin{cases} 1 & r \leq r_0 \\ \frac{1+\cos\left(\pi\frac{r-r_0}{w_m}\right)}{2} & r_0 < r < r_0 + w_m, \\ 0 & r \geq r_0 + w_m \end{cases} \quad \text{(S16)}$$

with $r_0 = 72$ Å and $w_m = 24$ Å. For the octahedral group O, the symmetry gather

$$(\mathcal{S}f)(\boldsymbol{x}) = \frac{1}{|O|}\sum_{S\in O} f(S^T\boldsymbol{x}) \quad \text{(S17)}$$

uses the exact signed coordinate permutations of Section 2.4. The Gaussian filter $G$ multiplies the three-dimensional Fourier transform of the cube, zero-padded by $2\lceil 4\sigma_G\rceil$ voxels per axis, by $\exp(-2\pi^2\sigma_G^2 f^2)$, with $\sigma_G$ in voxels. The result of Equation (14) is embedded into the full volume with zeros outside the central $168^3$ cube. The box check of Section 2.4 tests $r_0 + w_m + \lceil 3\sigma_G\rceil a \leq (n-1)\frac{a}{2}$. When it fails under the `taper` policy, the mask is multiplied by a separable raised-cosine face taper of width $\max(w_m, 20\ \text{Å})$ along each axis that reaches zero $\lceil 3\sigma_G\rceil$ voxels inside the cube boundaries.

For helical specimens, a cylindrical block along the helical axis $x$ replaces the central cube. Each screw-symmetry copy $(n_h, j)$ (indexing axial screw step $n_h$ and $n_{\text{pt}}$-fold rotational step $j$) resamples the gradient after rotating about $x$ by $n_h$ times the helical twist plus $j\cdot 360\frac{\circ}{n_{\text{pt}}}$ and translating along $x$ by $n_h$ times the axial rise using trilinear interpolation with zero boundary conditions. The Fejér-weighted screw average is then evaluated as

$$(\mathcal{H}f)(\boldsymbol{x}) = \frac{\sum_{n_h,j}\omega_{n_h} f_{n_h,j}(\boldsymbol{x})}{\max\left(1, \sum_{n_h,j}\omega_{n_h}\chi_{n_h,j}(\boldsymbol{x})\right)}, \quad \omega_{n_h} = 1 - \frac{|n_h|}{n_{\max}+1}, \quad \text{(S18)}$$

over $|n_h| \leq n_{\max}$ and $j = 0, \ldots, n_{\text{pt}} - 1$, where the indicator $\chi_{n_h,j}(\boldsymbol{x}) = 1$ if and only if the transformed source coordinate falls within the volume bounds. The complete blur-last helical gradient hook then takes the form

$$\tilde{g} = G\star P_{\text{ax}}[m_x\cdot\mathcal{H}(m_r g)], \quad \text{(S19)}$$

where $m_r$ is the radial cylindrical support mask, the 40 Å face taper $m_x$ ends $\lceil 3\sigma_G\rceil$ voxels inside the boundaries, and $P_{\text{ax}}$ projects out the mean-subtracted axial ($x$-only) profile weighted by $m_x m_r$. The screw sense is determined automatically from the initial map.

### S2. Additional methods

#### S2.1. Specimen-specific adaptations for T20S, TMV and Phi92

For the T20S proteasome, we use the same forward model and optimizers under D7 point-group symmetry. Because seven-fold rotations around the symmetry axis do not map Cartesian grid coordinates onto integer voxels, the symmetry gather $\mathcal{S}$ uses trilinear interpolation; the refinement runs across three sequential stages of 24, 16 and 8 epochs using the blur-last filter ordering.

For the helical specimens (TMV and Phi92), we replace the point-group symmetry gather with a screw-symmetry average $\mathcal{H}$ about the helical axis $x$. This operator forms a weighted sum over $2n_{\max}+1$ screw-transformed copies of the gradient, where copy $n_h$ is rotated by $n_h$ times the helical twist and translated along $x$ by $n_h$ times the axial rise, weighted by the triangular Fejér kernel $\omega_{n_h} = 1 - \frac{|n_h|}{n_{\max}+1}$ for $|n_h| \leq n_{\max}$. At each voxel, the screw sum is normalized by the sum of weights of those screw copies whose source coordinates fall inside the volume bounds; for Phi92, each screw step is additionally averaged over the six-fold rotational symmetry C6. To suppress longitudinal density stripes, the helical hook projects out pure axial ($x$-only) gradient modes, and replaces the spherical support mask with a soft cylindrical mask (radius 100 Å for TMV and 115 Å for Phi92). The complete blur-last helical hook thus applies: radial cylindrical masking, screw averaging, cylindrical re-masking with a 40 Å raised-cosine taper at both $x$ boundaries, axial-mode projection and a final 2 Å Gaussian blur. In our helical runs, particle poses and defocus offsets are held fixed, neighbour copies are omitted, and the initial volumes are hard low-pass filtered on the voxel grid of $V$ at 15 Å (TMV) and 5 Å (Phi92). For TMV, the volume gradient in each Fourier shell is additionally weighted by the radial Wiener factor $2\frac{\text{FSC}}{1+\text{FSC}}$ computed from the half-map FSC after epoch 0, which introduces a weak radial amplitude coupling between the two TMV halves.

#### S2.2. High-frequency rebound and phase-randomization diagnostic

Because the phase-randomization check randomizes the Fourier phases of the completed half maps rather than re-running the iterative reconstruction from phase-randomized input data as in Chen *et al.* (2013), it detects high-frequency FSC rebound artefacts rather than proving statistical independence of the half maps. For apoferritin, this diagnostic uses three criteria:

1. The pooled half-map FSC in the high-frequency band $\frac{1}{4.5} \le |\boldsymbol{k}| < 1.0$ Å$^{-1}$ must not exceed 0.1. Because the 4 Å Gaussian gradient filter attenuates update amplitudes at $\frac{1}{4.5}$ Å$^{-1}$ to about $2 \times 10^{-7}$ of their low-frequency value, a large correlation in this band indicates shared filter leakage rather than signal from the data.
2. The pooled half-map FSC in the transition band $\frac{1}{5} \le |\boldsymbol{k}| < \frac{1}{4.5}$ Å$^{-1}$ must not exceed 0.2.
3. The phase-randomization rebound check adapted from Chen *et al.* (2013) must pass across all tested cutoffs and random seeds.

In the phase-randomization check, the Fourier phases of each completed half map are randomized independently beyond a spatial-frequency cutoff $f_c$. Denoting the unmasked half-map FSC by $\text{FSC}_t$ and that of the phase-randomized half maps by $\text{FSC}_n$, we compute the phase-randomized corrected curve

$$\text{FSC}_{\text{true}} = \frac{\text{FSC}_t - \text{FSC}_n}{1 - \text{FSC}_n} \quad (|\boldsymbol{k}| > f_c). \qquad \text{(S20)}$$

We evaluate Equation (S20) across three random seeds and three spatial-frequency cutoffs (15 Å, 10 Å and twice the sustained 0.143 crossing resolution of $\text{FSC}_t$). The check passes if, for every combination of seed and cutoff, (i) $\text{FSC}_{\text{true}}$ first crosses below 0.143, (ii) no three consecutive Fourier shells beyond that crossing reach or exceed 0.143 up to the Nyquist frequency, and (iii) $|\text{FSC}_n| \le 0.05$ from two shells past $f_c$ to Nyquist. Criterion (ii) flags any high-frequency FSC rebound.

For T20S and the helical specimens, we apply criteria (1) and (3) without criterion (2). On TMV and T20S, a reconstruction that exceeds the 0.1 threshold in criterion (1) still passes if each half map correlates with the external reference model across the same high-frequency band by at least $0.8\sqrt{\text{HF FSC}}$; Phi92 needs no such rule because its evaluation grid reaches Nyquist before $\frac{1}{4.5}$ Å$^{-1}$. Any reconstruction that fails this diagnostic is still reported in our tables, with its pre-registered comparison marked as not valid.

#### S2.3. Map-to-model alignment and lattice shells

To evaluate the reconstructed volumes, we compared each map, calculated as the mean of its two aligned half maps, with an independent atomic model or high-resolution cryo-EM map. For atomic references, non-hydrogen atoms were weighted by atomic number and occupancy and convolved with a 1 Å Gaussian kernel. Before generating the reference density, the atomic model was rigidly docked into its deposited cryo-EM map, considering both hands. Each reconstruction was then aligned to the reference by searching over handedness, isotropic voxel scale and sub-voxel translation. For apoferritin, alignment was performed on a $112^3$ grid of 1.5 Å voxels with a soft spherical mask of 70 Å radius and 6 Å raised-cosine edge. The voxel scale was searched from 0.95 to 1.05 in increments of 0.005. TMV and Phi92 were evaluated on their respective grids using cylindrical masks.

For calculating $C_\text{w}$, the shell-wise radial Fourier amplitudes of each scored map were scaled to those of the reference on the unmasked grid, while Fourier phases were preserved. This normalization makes the score insensitive to isotropic amplitude filters, such as $B$-factor sharpening. The voxel scale was fitted once for each map family using the unfiltered half-map mean and was then held fixed across its variants. Handedness and sub-voxel translation were fitted for each map.

Because periodic probe scanning at step $\Delta r \approx 20$ Å imprints reciprocal-space lattice artefacts on Fourier shells at $\frac{n_l}{20}$ Å$^{-1}$ ($n_l = 1, 2, 3$) and $n_l \frac{\sqrt{2}}{20}$ Å$^{-1}$ ($n_l = 1, 2$), we exclude these lattice shells, using a half-width of 0.004 Å$^{-1}$, when computing $C_\text{w}$ and the model 0.5 crossing. The TMV and Phi92 evaluations use the same metrics with their respective atomic references, voxel grids and cylindrical masks, while T20S omits $C_\text{w}$ and is scored primarily against the cryo-EM map EMD-6287 (Campbell *et al.*, 2015). EMD-6287 was selected before the run because no matched atomic coordinate file had been configured at pre-registration. PDB 1PMA was reported alongside it as a secondary reference. The excluded scan-lattice shells were placed using the nominal scan step of 20.0 Å, whereas the reconstruction used the calibrated step of 20.6 Å. The values of $C$, $\text{FSC}_{10}$ and $\text{FSC}_8$ are unaffected by this choice. $C_\text{w}$ decreases by 0.009–0.013 for every map, and the model 0.5 crossing shifts by up to 0.25 Å (Section 3.2.2). We report $C_\text{w}$ at the pre-declared 20.0 Å step and give the 20.6 Å values beside it, and we report model 0.5 values under both lattice-shell definitions.

### S3. Log of pre-registered experimental and diagnostic tests

Our timestamped laboratory log records, in chronological order of pre-registration, all 29 pre-registered experimental and diagnostic tests referenced in this study, including exploratory apoferritin tests conducted prior to the final refinement configuration as well as tests whose outcomes were negative. For each entry, the log contains a pre-declaration specifying the scientific question, the computational run, the epoch selection rule, the quantitative decision thresholds and the validation gates. Every pre-declaration was recorded before the corresponding run generated any numerical output, and all protocol amendments were logged prior to executing the modified runs. The log keeps each result entry verbatim, and each pre-registered outcome stands as declared even where subsequent diagnostics identified a flaw in an underlying test premise. Some tests below use, or compare against, an earlier ('legacy') order of the gradient filters, $m \cdot \mathcal{S}[G \star (mg)]$, in which the support mask comes after the Gaussian filter. This order couples the half maps at high frequencies. The final maps of this paper use the blur-last order (Section 2.4).

Two descriptive lookups, the PDB 7A4M sequence check (`apoferritin-sequence`) and the GPU compute-time tally (`apoferritin-cost`), carried no pass/fail threshold and are not counted among the 29 tests.

Two timestamp details in the log are: the `apoferritin-fair v2` pre-declaration refers to the initial scoring check as "the

21:05 verdict” whereas the corresponding log entry is stamped 20:56, and three entries (`apoferritin-strong`, `apoferritin-fair` and `apoferritin-fair v2`) carry a calendar date without a clock time, although their line positions in the append-only log precede all runs that depend on them.

### S4. Summary of exploratory 14-arm apoferritin refinements

Before pre-registering our final apoferritin protocol, we ran 14 exploratory End2EndPtychoSPA refinement arms on the full 19,064-particle pool. All 14 exploratory arms used the legacy gradient-filter ordering and preceded all pre-registered apoferritin tests (Section S3). Because these runs used 19,064 particles rather than the 11,552-particle subset of EMD-19425, they are not a particle-matched comparison with the deposited map, and none of these runs was used as the final apoferritin reconstruction (whose blur-last recipe, R1 initialization and final-epoch selection rule were fixed independently in `fixed-runs`). We list all 14 exploratory arms here because their map-to-model metrics against PDB 7A4M were inspected when comparing these earlier parameter settings.

### S5. Diagnostic tests on the helical near-Nyquist boundary band

Even with the blur-last gradient hook, the TMV and Phi92 End2EndPtychoSPA reconstructions failed our high-frequency phase-randomization diagnostic because of a narrow correlation band shared between the two half maps just below the Nyquist frequency (2.33–2.54 Å in TMV and 2.6–2.7 Å in Phi92). When the epoch-0 TMV update was band-pass filtered to this frequency interval, the boundary zones within 50 Å of the two axial $x$ faces where the reconstruction box truncates the helix contained 76.6% of the inter-half cross-power, with a local correlation of 0.77, whereas the interior region contained 43% of the band power but only 4.0% of the cross-power (local correlation 0.04). In Phi92, the axial $x$-face zones contained 89.1% of the shared band cross-power (local correlation 0.37), with the cross-power peak located 3–6 voxels inside each $x$ boundary in both specimens.

The per-epoch volume updates showed that this correlation is regenerated during each gradient step whenever the spatial-frequency filter has nonzero weight in the near-Nyquist band. In TMV epoch 0 (which used the Gaussian filter), the volume update had a band FSC of 0.394 with 77% of the cross-power at the $x$ faces, whereas in epochs 1 and 2 (which used the radial Wiener filter whose weight is zero in that band) the update band FSC dropped to 0.0004 and −0.0001. In Phi92, all four epochs used the Gaussian filter and gave update band FSC values of 0.167, 0.159, 0.162 and 0.154 with 88–89% of the cross-power at the $x$ faces, about $10^4$ above the single-precision (`float32`) rounding floor.

We ran three pre-registered diagnostic tests to investigate three possible causes of this boundary band (Table S2). None of the three tests reproduced the band at the observed amplitude.

#### S5.1. Helix-Nyquist: spectral transfer of the helical gradient hook

Inspection of the code shows that the FFT Gaussian blur pads each axis by $2\lceil 4\sigma \rceil$ voxels, which prevents circular wrap-around between the opposite $x$ boundaries of the box (arm A2 with extra $x$-padding is identical to baseline arm A0). The blur margin of $\lceil 3\sigma \rceil$ voxels sets only the inset of the axial face taper, and widening it to $\lceil 4\sigma \rceil$ left the output unchanged (arm A1 equals arm A0). The legacy filter ordering produced strong high-frequency correlation as a positive control (arm A4), but passing a shared low-frequency update through the blur-last helical hook gave a near-Nyquist band power fraction of only $7.3 \times 10^{-19}$ in TMV and $2.1 \times 10^{-17}$ in Phi92, compared with $3.4 \times 10^{-12}$ and $1.3 \times 10^{-11}$ in the actual refinement updates. Trilinear interpolation during screw averaging increased this floor by a factor of 600 in Phi92 and 2 in TMV, but the resulting power remained five to six orders of magnitude below the experimental updates (while reproducing the equatorial $k_x = 0$ plane correlation of 0.110–0.403 versus 0.121 in TMV epoch 0). The helical gradient hook

**Table S2**

Pre-registered diagnostic tests investigating the helical near-Nyquist correlation band (0.400–0.435 $Å^{-1}$ in TMV and 0.370–0.385 $Å^{-1}$ in Phi92). Where three band FSC values are listed, they correspond to random seeds 0 / 1 / 2.

| Test | Candidate and design | Key numbers | Verdict as pre-registered |
|---|---|---|---|
| helix-Nyquist<br>P 10-11 12:22<br>A1 12:34<br>R 14:42 | The blur-last hook writes the band from a shared low-frequency update (blur padding, blur-margin width, screw interpolation). Planted pairs $g_i = s + n_i$ through the real hook in float64; arms A0 (current), A1 (blur margin $\lceil 4\sigma \rceil$), A2 (extra x padding), A3, A4 (legacy order, positive control), A5 (shared start), A6 (no screws), A7 (Wiener weight); seeds 0–2. | Bar $3/\sqrt{N_{\text{ind}}}$: TMV 0.001813, Phi92 0.002007. A0 band FSC (K = 1): TMV −0.00688 / −0.01509 / −0.00501; Phi92 +0.00213 / −0.00489 / −0.00766. A4: TMV 0.867, Phi92 0.673. Band power of the hook output from a shared field: TMV $7.3 \times 10^{-19}$, Phi92 $2.1 \times 10^{-17}$, against $3.4 \times 10^{-12}$ and $1.3 \times 10^{-11}$ in the real updates. | The planted model does not reproduce the band; no arm can remove it. Positive control holds. The hook writes 5 to 6 orders of magnitude too little; the hypothesis is refuted by code and test. |
| helix-face<br>P 10-11 15:52<br>R 21:47 | The model volume ends at the x faces, but the helix in the data continues. Linear projection forward through the real sampler arithmetic, all table items (TMV 4,436; Phi92 1,651), real poses and scan windows, noise at the measured level. Arms F0 (truncated), F1 (no model error), F2 (longer box), F3 (screw-periodic sampling), F4a / F4b (x taper), F5 (hard cutoff). | Bar (band): TMV 0.034078, Phi92 0.029750. F0 band FSC: TMV −0.001614 / +0.022065 / −0.008389; Phi92 +0.008580 / +0.009235 / +0.009998; F1 equal to 5 digits. | Planted model inadequate. Cause confirmed: no (TMV), no (Phi92). No fix arm credited. |
| scan-overlap<br>P 10-11 22:20<br>A1 22:30<br>R 10-12 00:01 | Items of different halves whose scan windows intersect fit the same measured patterns and the same noise. One TMV Gaussian epoch for arm D (drop every overlapping item) and arm C (drop the same number at random), 2,079 items per half in each arm (A1: equal counts). | Overlap: 1,031 cross-half pairs; 122 (5.50 %) and 139 (6.27 %) items; 1.67 % and 1.88 % of the patterns. Band FSC: reference 0.39425, D 0.38859, C 0.40695. Confirm bar 0.040695, refute bar 0.20348. | Refuted ($D_b/C_b$ = 0.955). |

itself therefore does not generate the near-Nyquist boundary band from a shared low-frequency update.

Our initial test script evaluated the hook in `float32`, where adding a shared component to independent noise vectors imparts correlated single-precision rounding errors at the $10^{-18}$ power floor; Amendment 1 promoted the test evaluation to `float64` before reading any test outcomes, retaining a `float32` reference row (`A0f32`).

#### S5.2. Helix-face: axial box truncation in a linearized forward model

In our second diagnostic (`helix-face`), we tested whether truncating a continuous helical rod at the axial $x$ faces of the volume box injects a shared near-Nyquist band into the backprojected gradients under a linear projection forward model at matched experimental noise. At the experimental shot-noise level, the truncated-helix arm (F0) and the truncation-free null arm (F1) gave identical band FSC values to five decimal places. This indicates that linear truncation error would only exceed the detection threshold if the detector noise were reduced by a factor of 0.00236 in TMV and 0.00048 in Phi92. In the noise-free shared gradient component, the inter-half band FSC after the blur-last hook was 0.920 in TMV and 0.971 in Phi92, with 73.8% and 55.6% of its cross-power within 50 Å of the $x$ faces (93.5% and 88.4% before the hook), matching the spatial distribution observed in the experimental refinements (77% and 89%). Thus, axial box truncation of a continuing helix does generate a face-localized near-Nyquist band at the observed position, but in a linear single-step model its amplitude is three to four orders of magnitude too weak to explain the experimental band. Among the mitigations tested on the noise-free component, applying the axial $x$-face taper both before and after screw averaging (arm F4b) reduced the shared band power to 0.41% (TMV) and 0.014% (Phi92), while screw-periodic boundary sampling (arm F3) reduced it to 3.9% and 0.05%.

#### S5.3. Scan-overlap: shared diffraction frames between helical half sets

The TMV gold-standard half sets are split by helical filament (no filament is shared between halves), but 31 of the 60 scans contain filaments assigned to both halves, and in 12 scans the $12 \times 12$ scan windows of segments in opposite halves intersect. In our third diagnostic (`scan-overlap`), arm D removed every segment whose scan window shared any diffraction pattern with the opposite half set (yielding disjoint diffraction frames), while control arm C removed an equal number of segments at random (retaining 90–92% of the shared frames). After one Gaussian refinement epoch on 2,079 segments per half, the near-Nyquist band correlation remained essentially unchanged ($D_b = 0.389$ versus $C_b = 0.407$, with 78.0% of the cross-power at the $x$ faces in both arms compared with 76.6% in the full dataset). This rules out shared diffraction frames as the cause of the boundary band.

Possible mechanisms that were not tested include multislice propagation or probe-boundary interactions at the axial box faces, growing pose or defocus errors near the segment ends, and non-linear amplification across iterative updates.

### S6. Sensitivity analysis of the apoferritin benchmark comparison

Because the apoferritin End2EndPtychoSPA reconstruction clears the pre-declared $C_w$ and $FSC_{10}$ benchmark thresholds by small margins (+0.00375 and +0.00930), we performed three pre-registered sensitivity checks (Table S3) to test whether three choices in the evaluation pipeline (the docking frame of PDB 7A4M, the scan-lattice shell step and the relative map magnification) alter the sign of the benchmark margins.

#### S6.1. Reference model placement (`apoferritin-placement`)

Our primary evaluation docked PDB 7A4M into EMD-19425 via a coarse rotational search, whereas the `apoferritin-placement` test evaluated PDB 7A4M directly in its deposited crystallographic symmetry frame. The 24 biological assembly matrices of PDB 7A4M are exact signed coordinate permutations sharing a single fixed point at 68.249605 Å along each Cartesian axis, placing the four-fold symmetry axes directly along $x, y$ and $z$ without reference to any experimental map (verified to $5.7 \times 10^{-14}$ Å with a minimum octahedral self-correlation of 0.99999999).

In this script, the pre-declared reproduction gate failed on a single metadata entry: although the End2EndPtychoSPA map and both EMD-19425 maps reproduced all baseline numbers to within $1.2 \times 10^{-16}$, our R1 RELION map differed in $FSC_8$ by $1.0923 \times 10^{-5}$ (exceeding the $10^{-5}$ gate tolerance) because the test script read the full floating-point MRC header voxel size of 1.59140587 Å whereas the baseline evaluation rounded the R1 voxel size to 1.5914 Å (at 1.5914 Å, all numbers reproduce identically). Because the gate failed, we report the new-placement numbers as secondary checks.

Evaluated in the deposited octahedral frame of PDB 7A4M, all four reconstructions achieved higher scores ($C_w$ increased by 0.036–0.039 and $FSC_8$ by 0.060–0.069) because the 1°-grid rotational fit into EMD-19425 had left the reference slightly misaligned relative to the grid axes shared by the octahedral maps (reducing octahedral self-correlation to 0.851 and lowering all maps equally). Under the exact symmetry frame, all three benchmark margins increased (Table S3).

**Table S3**
Benchmark margins (score(End2EndPtychoSPA) − [score(EMD-19425 half mean) + margin], with margins $+\frac{0.010}{+}\frac{0.030}{+}0.030$) across five evaluation variants. Every margin remains positive across all variants; as noted in the rightmost column, only the primary row and the `start-scale` row carry formal pre-registered verdicts.

| Scoring variant | $C_w$ | $FSC_{10}$ | $FSC_8$ | Status |
|---|---|---|---|---|
| Primary evaluation (7A4M docked into EMD-19425, lattice 20 Å) | +0.00375 | +0.00930 | +0.05786 | primary evaluation: benchmark passed |
| Deposited symmetry frame of 7A4M, lattice 20 Å | +0.00680 | +0.01235 | +0.06266 | secondary result (apoferritin-placement: gate failed) |
| Primary placement, lattice 20.6 Å | +0.00796 | +0.00929 | +0.05785 | secondary result (apoferritin-lattice: gate failed) |
| Deposited frame, lattice 20.6 Å | +0.01129 | +0.01235 | +0.06266 | secondary result |
| Each map evaluated at its own optimal scale | +0.00253 | +0.00486 | +0.05124 | start-scale C2: scale-robust benchmark result |

#### S6.2. Reciprocal scan-lattice step (`apoferritin-lattice`)

Our primary evaluation excluded the reciprocal scan-lattice shells at the nominal 20.0 Å scan step, whereas the calibrated scan step used during refinement was 20.6 Å (20 Å × 1.03). Sharing the same reproduction gate as `apoferritin-placement`, `apoferritin-lattice` is likewise reported as a secondary check: shifting the excluded lattice shells to 20.6 Å left $C$, $\mathrm{FSC}_{10}$ and $\mathrm{FSC}_8$ unchanged (maximum difference 0.0) and reduced $C_\mathrm{w}$ by 0.0088–0.0131 across maps while increasing the End2EndPtychoSPA $C_\mathrm{w}$ margin to +0.00796. In contrast, the model 0.5 crossing shifted by −0.25 to +0.16 Å (moving End2EndPtychoSPA from 6.90 Å to 7.06 Å and R1 from 7.41 Å to 7.17 Å) because the 0.5 crossings fall near the $2\frac{\sqrt{2}}{\mathrm{step}}$ lattice reflection; for this reason, we do not interpret model 0.5 differences at the 0.1 Å level.

Within the same check, linear interpolation of RELION's internal half-map FSC table for run R1 (reported as 6.881755 Å by RELION's discrete shell lookup) yielded an interpolated 0.143 crossing of 6.8274 Å (and 0.5 crossing at 7.4755 Å), in exact agreement with our native-grid unmasked half-map FSC evaluator (6.827 Å).

#### S6.3. Relative voxel scaling (`start-scale`)

Because the initial R1 table maps were written with a header voxel size of 5.1 Å but read by the initialization routine at 5 Å × 1.03 = 5.15 Å (a factor of 1.0098), we performed a fine magnification scan (step 0.0025) against PDB 7A4M (`start-scale`). This check passed its reproduction gate exactly and found optimal isotropic scale factors of 0.9950 for End2EndPtychoSPA (parabolic sub-grid peak at 0.99567, within the pre-declared ±0.005 tolerance of criterion C1), 1.0225 for the EMD-19425 half-map mean and 1.0050 for RELION R1. Evaluating each map at its own optimal isotropic scale (criterion C2) preserved positive benchmark margins of +0.00253, +0.00486 and +0.05124 across $C_\mathrm{w}$, $\mathrm{FSC}_{10}$ and $\mathrm{FSC}_8$, even though rescaling EMD-19425 by 1.0225 increased its individual scores by 0.0012, 0.0044 and 0.0066.

The 20 Å hard low-pass-filtered initial volume $V^0$ cannot by itself calibrate a 1% magnification offset over the 8–30 Å band because its nonzero Fourier content is confined to 20–30 Å, where its band correlation (0.260–0.380) sits near the random-orientation floor (0.32), yielding noisy scale optima (1.040 on $V^0$ versus 0.973 on the unfiltered table maps at 5.15 Å).

#### S6.4. Sensitivity of the ResLog slope (`ResLog-robust`)

Applying the same reference-placement and scan-lattice variants to the four-point ResLog series (`ResLog-robust`, which passed its reproduction gate across all 13 resolutions and six $B$-factors) showed that the fitted $B$-factor difference $B(\mathrm{2D}) - B(\mathrm{End2EndPtychoSPA})$ is sensitive to the lattice step: the difference was 426.0 Å² under the primary evaluation and 626.1 Å² in the deposited PDB 7A4M frame at the 20.0 Å lattice setting, but narrowed to 54.7 Å² and 65.1 Å² when excluding lattice shells at the calibrated 20.6 Å step (below the pre-declared 136 Å² robustness bar). Replacing the discrete shell crossing with a smooth logistic fit to the map-to-model FSC curve yielded $B$-factor differences of 160.3 Å² (primary placement) and 122.2 Å² (deposited frame).

### S7. Enantiomorphic hand verification

To check the enantiomorphic hand of each final reconstruction against an independent atomic reference (`hand-check`), we evaluated each delivered volume and its axial $z$-mirror image with the automatic hand-inversion search disabled. We measured hand preference via the differences $d_\mathrm{band}$ (in 8–30 Å band correlation $C$) and $d_\mathrm{F8}$ (in $\mathrm{FSC}_8$) between the delivered and mirrored volumes, using the apoferritin reconstruction as a positive control to set the decision thresholds $M = \max(0.02, 0.5 d_\mathrm{apoferritin})$ (requiring the apoferritin control differences to exceed 0.04 and three times the inter-half noise).

Each specimen was evaluated against its corresponding atomic coordinate model (PDB 7A4M for apoferritin, PDB 4UDV for TMV, single-body PDB 21VM gp130 for Phi92 and PDB 1PMA for T20S). Both validation gates passed: disabling the hand loop reproduced all baseline scores on the preferred branch (gate G1), and scoring the synthetic PDB 1PMA density map against itself versus its mirror image yielded 1.000 versus 0.367 in $C$ and 1.000 versus 0.213 in $\mathrm{FSC}_8$ (gate G2), confirming that the 8–30 Å and 8 Å metrics resolve the chiral hand of the T20S barrel.

As shown in Table S5, the stored TMV and T20S End2EndPtychoSPA map files have inverted handedness, directly inheriting the inverted hand of their 2D-route initializations (`cryoSPARC J364` and `RELION table_gs_a1`, whereas our apoferritin pipeline explicitly inverted $z$ via `--mirror-hand` at initialization), while the deposited EMD-19885 and EMD-67291 maps have the correct hand. Because our evaluation scripts automatically search both hands and score the matching branch, all quantitative metrics reported in the manuscript are unaffected, and the TMV and T20S volumes simply require a $z$-reflection before rendering or deposition.

On Phi92, the delivered hand was strongly preferred over the mirror image in both metrics ($C$ of 0.5963 versus 0.0537, and 0.5967 versus 0.0537 in `Phi92-fix`), but the pre-registered test was classified as formally inconclusive because the $\mathrm{FSC}_8$ difference (+0.240) fell slightly below the apoferritin-derived margin (0.269), which exceeds almost the entire absolute $\mathrm{FSC}_8$ correlation of the lower-resolution Phi92 map (0.2815).

**Table S4**

Scaling of apoferritin reconstruction quality across nested particle subsets evaluated against PDB 7A4M. At $N = 1,444$, the two entries in $\mathrm{FSC}_8$ correspond to two independent random subset draws. Map-to-model FSC 0.5 crossings are reported with the reciprocal scan-lattice shells excluded at the nominal 20.0 Å step and at the calibrated 20.6 Å step. ResLog $B$-factors are obtained from ordinary least-squares fits of $1/d_\mathrm{res}^2$ against $\ln N$ across all four subset sizes ($B = 2/$ slope).

| N | $\mathrm{FSC}_8$ | | model FSC 0.5 (Å), lattice 20 / 20.6 Å | |
|---|---|---|---|---|
| | 2D route | End2EndPtychoSPA | 2D route | End2EndPtychoSPA |
| 1,444 | 0.464 / 0.466 | 0.432 / 0.450 | 8.32 / 8.32 | 8.34 / 8.34 |
| 2,888 | 0.576 | 0.618 | 7.79 / 7.79 | 7.75 / 7.75 |
| 5,776 | 0.603 | 0.662 | 7.57 / 7.50 | 7.30 / 7.38 |
| 11,552 | 0.659 | 0.710 | 7.41 / 7.17 | 6.90 / 7.06 |
| ResLog B (Å²) | – | – | 1084 / 821 | 658 / 766 |

**Table S5**
Enantiomorphic hand evaluation across experimental reconstructions. Control margins derived from apoferritin are $M_{\text{band}} = 0.277$ and $M_{\text{F8}} = 0.269$ (with apoferritin inter-half noise of 0.003 in $C$ and 0.057 in $\text{FSC}_8$). Deposited EMDB half-map means were evaluated only as external checks and never used as alignment references.

| Map (reference model) | role | $C$, delivered / mirror | $d_{\text{band}}$ | $\text{FSC}_8$, delivered / mirror | $d_{\text{F8}}$ | Verdict |
|---|---|---|---|---|---|---|
| apoferritin stage 2, epoch 15 (7A4M) | control | 0.8523 / 0.2982 | +0.554 | 0.7104 / 0.1719 | +0.538 | valid; hand correct |
| TMV End2EndPtychoSPA fjw_ep0 ep2 (4UDV) | map | 0.1081 / 0.8397 | −0.732 | 0.0047 / 0.6638 | −0.659 | wrong |
| T20S End2EndPtychoSPA blur2 ep3 (1PMA) | map | 0.3341 / 0.7825 | −0.448 | 0.1676 / 0.6013 | −0.434 | wrong |
| Phi92 End2EndPtychoSPA r4B_lr2.5 ep0 (21VM) | map | 0.5963 / 0.0537 | +0.543 | 0.2815 / 0.0419 | +0.240 | inconclusive as pre-registered |
| apoferritin RELION R1 | secondary result | 0.332 / 0.799 | | 0.182 / 0.659 | | wrong |
| TMV cryoSPARC J364 | secondary result | 0.050 / 0.895 | | −0.021 / 0.854 | | wrong |
| EMD-19885 half mean | secondary result | 0.883 / 0.048 | | 0.796 / −0.021 | | correct |
| T20S RELION aff half mean | secondary result | 0.337 / 0.783 | | 0.179 / 0.637 | | wrong |
| EMD-67291 half mean | secondary result | 0.722 / 0.351 | | 0.640 / 0.116 | | correct |

Because the helical symmetry lattice for PDB 21VM was docked into EMD-19430, this check verifies hand consistency with EMD-19430.

On apoferritin, where $\alpha$-helical chirality in PDB 7A4M defines the hand, the delivered End2EndPtychoSPA volume scored 0.8523 in $C$ versus 0.2982 when mirrored along $z$.

### S8. Haemoglobin reconstruction controls

#### S8.1. Evaluation grids and initialization controls

The primary comparisons in Section 3.1 use native evaluation grids of 1.2 Å for RELION and 0.72 Å for End2EndPtychoSPA. Under the molecular mask, the **de novo** RELION and end-to-end maps crossed FSC 0.143 at 2.59 and 2.36 Å, respectively. The latter lies beyond the 2.4 Å Nyquist limit of the original ground-truth grid and does not establish resolution at that scale; the primary comparison therefore uses FSC 0.5.

When both maps were evaluated on the common 1.2 Å grid, End2EndPtychoSPA improved the FSC 0.5 resolution by 0.17 Å under the molecular mask and 0.36 Å under the soft spherical mask of 42 Å radius. On the common 0.72 Å grid, the corresponding gains were 0.18 and 0.36 Å. On the native grids, the spherical-mask FSC 0.5 resolutions were 3.70 Å for RELION and 3.35 Å for End2EndPtychoSPA.

For the control from the 20 Å ground-truth start (Section 3.1), the spherical-mask FSC 0.5 resolution improved from 3.36 to 3.15 Å. The common-grid gains were 0.09–0.15 Å under the molecular mask and 0.21–0.23 Å under the spherical mask. At FSC 0.143, the RELION curve remained above threshold through its last shell (2.45 Å); the end-to-end crossing at 2.29 Å lay beyond the ground-truth grid's 2.4 Å band limit. The native-grid molecular-mask FSC 0.5 gains, calculated before rounding, were 0.19 Å from the **de novo** start and 0.14 Å from the 20 Å start.

#### S8.2. Reconstruction repeats

The two recipe-seed runs shown in Figure 4 used the same minibatch ordering: in this software version, changing the recipe seed did not re-seed the sampler. Their molecular-mask FSC 0.5 resolutions were 3.2187 and 3.2184 Å. A repeat with the original batch order reached 3.2180 Å, while two additional runs with independent orderings reached 3.2075 and 3.2068 Å. The spread across the tested orderings was 0.0119 Å; all improved on the 3.4081 Å RELION start. The two independent-order runs reached approximately 3.34 Å under the spherical mask.

Relative to the first run, the maximum absolute voxelwise difference divided by the peak map amplitude was approximately 2% for repeated runs with the same batch order and 9% for the independent-order runs. These repeats probe minibatch ordering and GPU nondeterminism while retaining the same initial maps and half-set assignments; they do not assess variability across independent initializations or half-set splits.

### S9. Ground-truth simulation of myoglobin: orientation search limits reconstruction of a 17.9 kDa protein

To test the lower molecular-weight limit of cryo-electron ptychographic single-particle analysis under known ground-truth conditions, we simulated 50 micrographs of sperm-whale myoglobin (PDB 1A6M, 17.9 kDa) embedded in a 60 Å thick layer of atomistic vitreous ice at an accelerating voltage of 100 keV and an electron fluence of 30 e/Å² (Figure S1). The incident probe was configured with a convergence semi-angle of $\alpha = 10.28$ mrad and 1.0 µm underfocus, scanning each micrograph across a $21 \times 21$ raster at a step size of 30.7 Å. Our multislice forward simulation included an absorptive potential parameterized from elemental inelastic scattering cross-sections and included the resulting inelastically scattered electrons on the detector to model unfiltered acquisition. Across the 50 simulated fields, 3,093 myoglobin molecules were placed at known 3D positions and orientations. Using only the simulated diffraction patterns without ground-truth input, our windowed single-sideband (SSB) defocus sweep recovered the true defocus $C_{10}$ to within 2% on every micrograph, and automated template picking on the empirical-Bayes SSB phase micrographs identified the particles with a pooled $F_1$ score of 0.96 (precision 0.98, recall 0.94, median in-plane centering error 3.0 Å).

To test whether the reconstructed 2D ptychographic phase micrographs preserve high-resolution structural signal at 17.9 kDa, we performed an oracle 3D reconstruction of the 3,004 picked and truth-matched particles using their exact ground-truth poses. With known orientations, the 2D ptychographic projections reconstructed to the Nyquist limit (2.5 Å resolution at FSC 0.5 against the ground truth) and gave a Rosenthal–Henderson $B$-factor of 68 Å² across nested particle subsets (Figure S1(e)).

When particle orientations were unknown, however, global pose search in the 2D route failed to align the 17.9 kDa projections. Whether initialized from a 40 Å or a 20 Å low-pass-filtered ground-truth reference, RELION 3D auto-refinement stalled at 11–14 Å resolution after one to two iterations: 97–99% of the assigned particle orientations remained more than 10° away from the true pose, with median angular errors of 125° (20 Å reference) and 140° (40 Å reference; Figure S1(f)) that are close to that of a uniform random distribution over SO(3). Repeating the refinement on particle projections extracted from the empirical-Bayes SSB phase micrographs produced the same failure. Despite these near-random orientations, the RELION gold-standard half maps agreed to 8.4 Å and the resulting map correlated with the ground-truth potential to 7.8 Å at FSC 0.143. Because the particle orientations are random, this correlation reflects template bias from the ground-truth-derived initial model rather than genuine resolution. Projections of the ground-truth potential at the misassigned poses had a median cross-correlation against the extracted particle images nearly identical to that obtained at the true poses (0.53 versus 0.55 at 6 Å low-pass), confirming that 2D projections of a 17.9 kDa protein in 60 Å of vitreous ice at 30 e/Å² lack sufficient contrast to distinguish between competing orientations.

To test whether local refinement can succeed when initialized within the correct orientation basin, we perturbed the true particle poses by a 10° rotation and 2 Å random in-plane translation. Starting from these locally perturbed poses, RELION local refinement converged to a median orientation error of 6.8° and a median translation error of 0.7 Å, yielding a three-dimensional map that reached 3.4 Å resolution against the ground truth (Figure S1(g,h)). From the same perturbed initialization, End2EndPtychoSPA ran its full refinement schedule of 16 epochs (8 epochs of stage 1 with pose refinement from the second epoch, then 8 epochs of stage 2). It did not converge toward the true poses. The median angular error grew from 10.0° to 13.5° (12.2° after a global gauge fit), and the fraction of particles within 10° of the truth fell from 49% to 26%. The median translation error ended at 2.8 Å (start 2.3 Å). The FSC against the ground truth was best after the second epoch (19 Å at FSC 0.5) and fell to 27 Å at the end. Over the same epochs, the half-map FSC 0.143 crossing improved from 28 Å to 22 Å in stage 2, so the two half maps agree more with each other but less with the truth (Figure S1(d,g,h)). From a local start, RELION local refinement therefore outperforms our gradient-based pose refinement in this low-contrast regime.

When End2EndPtychoSPA was initialized from the failed 2D-route poses of the global orientation search from the 20 Å reference (median angular error 125°), our local gradient-based refinement left the orientations unchanged after the 8 epochs of stage 1 (median error shifting from 125.3° to 124.9°, with only 2.6% of particles within 10° of the truth) and the reconstructed volume remained at 34 Å resolution against the ground truth. This is expected: our local Gaussian pose prior ($\sigma_R = 2°$ or 10°) restricts the refinement to a local basin, so it cannot search across 100° orientation errors. At this particle size, the refinement does not hold the poses even inside that basin (see above).

To test whether higher electron fluence or learned 2D denoising priors improve single-particle alignability, we measured the fraction of particles for which the true orientation achieves a higher projection correlation against the ground-truth map than 2,000 competing random orientations across 310 particles (Figure S1(i)). Increasing the incident dose from 30 e/Å² to 45 e/Å² (via an independent Poisson noise draw from the noise-free diffraction patterns) increased the fraction of correctly identified orientations from 0.38 to 0.61 at 8 Å low-pass and from 0.70 to 0.87 at 5 Å low-pass. Because our multislice simulation does not model cumulative radiation damage, this high-dose improvement is an upper bound: an electron dose of 45 e/Å² at 100 keV corresponds in damage to about 71 e/Å² at 300 keV (Naydenova *et al.*, 2019). In contrast, adding a plug-and-play DRUNet denoising prior to the 2D ptychographic reconstruction (with its regularization weight calibrated from the in-band noise variance) reduced the orientation-recovery fraction to 0.14 at 30 e/Å² and 0.33 at 45 e/Å² (at 8 Å) while adding spurious texture across both the vitreous ice and the protein. Applying an untrained deep image prior (trained on each scan and stopped at peak cross-validation against a binomial count split), either inside the ptychographic loop or as a post-reconstruction filter, reduced the orientation-recovery fraction to 0.00–0.05 on myoglobin and from 0.89 to 0.01–0.10 on haemoglobin by smoothing the particles into featureless envelopes. When the 45 e/Å² micrographs were refined through RELION across all 50 scans (using the same particle picks, half sets and 20 Å start), the increased dose reduced the median angular pose error from 127° to 35° and nearly tripled the fraction of particles aligned within 20° (from 13% to 36%), although 89% of the particles still remained more than 10° from their true orientation. Two repeat refinements gave medians of 34° and 54°, so the size of the gain varies between runs.

### S10. Ground-truth simulation methods

#### S10.1. Atomistic specimen and vitreous-ice model

The primary simulated specimen is human deoxy-haemoglobin (PDB 2DN2, 64.5 kDa (Park *et al.*, 2006)), and the Supporting Information benchmark uses sperm-whale myoglobin (PDB 1A6M, 17.9 kDa (Vojtěchovský *et al.*, 1999)). To model structural noise from the solvent, we built an atomistic vitreous-ice layer by tessellating three-dimensional Voronoi cells filled with randomly rotated, translated and mirrored copies of the periodic 100 Å TIP4P-Ew water box (Horn *et al.*, 2004) from MeasureIce (Brown & Hanssen, 2022) (32,985 water molecules at a density of 0.987 g/cm³). The Voronoi tessellation avoids periodic tile seams that would otherwise produce artificial lattice reflections in the projected potential. Each simulated micrograph has its own independent 80 Å thick vitreous-ice slab spanning a 900 × 900 Å field into which 70 protein molecules are embedded with uniform random three-dimensional orientations and a minimum centre-to-centre separation of 80 Å. Each composite solvated system was relaxed in OpenMM 8 (Eastman *et al.*, 2024) with all water molecules unconstrained and the protein heavy atoms harmonically restrained (energy minimization, followed by NVT equilibration cooled from 250 K to 100 K, and a final energy minimization). A rigid Kabsch alignment of each relaxed protein copy against the reference coordinates then defined its ground-truth pose.

#### S10.2. Multislice 4D-STEM simulation

We simulated 4D-STEM diffraction data at an accelerating voltage of 100 keV with a convergence semi-angle of $\alpha = 10.28$ mrad on a 192 × 192 pixel detector spanning a maximum scattering angle of 1.5$\alpha$. The probe defocus was set to $C_{10} = -1.0 \times 10^4$ Å (1.0 µm underfocus, matching the sign convention of the

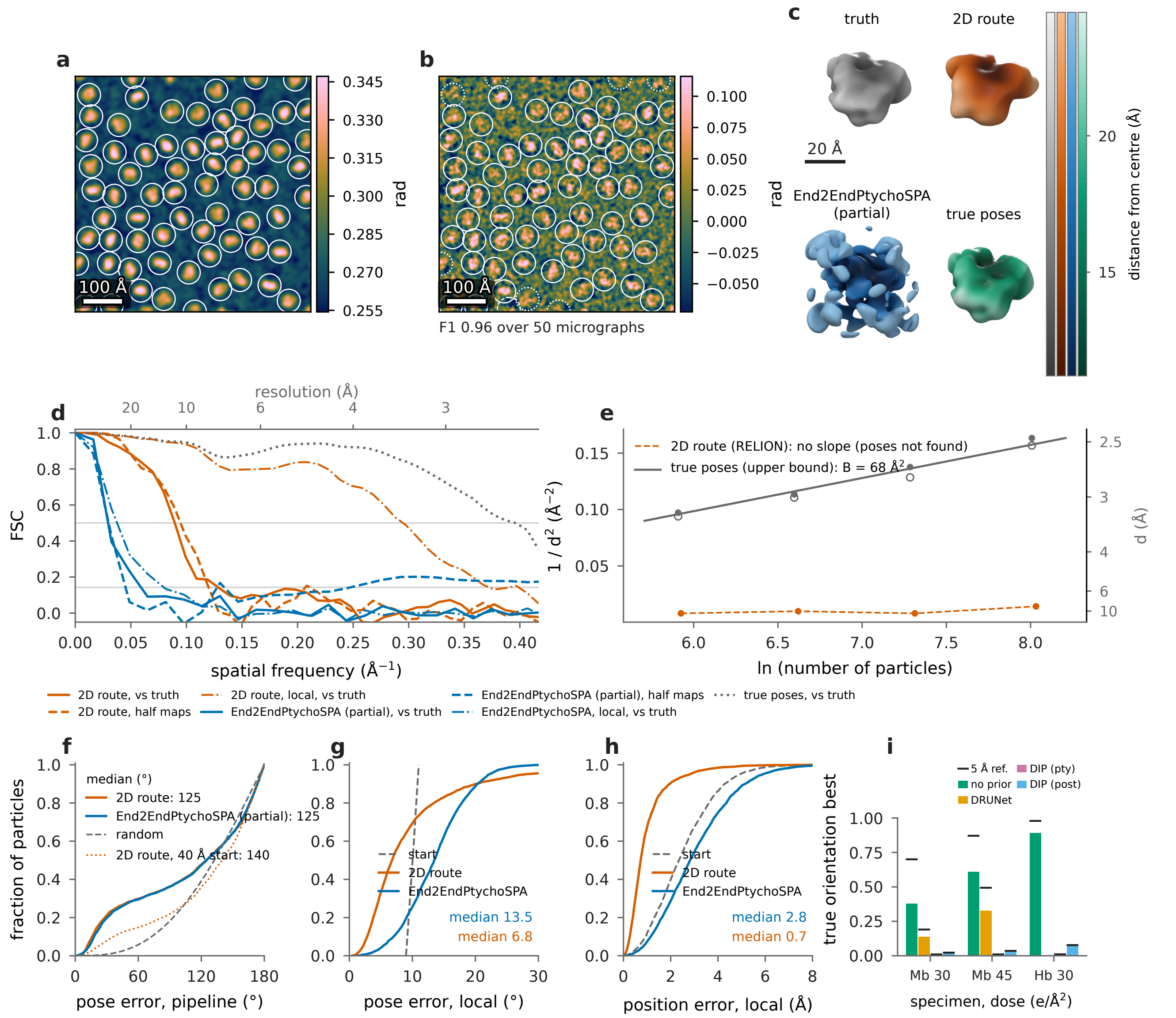


**Figure S1**
Ground-truth multislice simulation of sperm-whale myoglobin (PDB 1A6M, 17.9 kDa) across 50 micrographs containing 3,079 picked particles in 60 Å of atomistic vitreous ice at 100 keV and 30 e/Å². (a) Ground-truth projected phase of one micrograph, with circles marking the true particle centres. (b) Reconstructed 2D ptychographic phase micrograph overlaid with template picks (solid circles: matched to a true particle; dashed: unmatched false positives; dotted: missed true particles). (c) Isosurface renderings contoured at the molecular volume for the ground-truth potential, the 2D-route RELION reconstruction, End2EndPtychoSPA after stage 1 (initialized from the failed 2D route) and the oracle reconstruction using the true particle poses. (d) Gold-standard half-map FSC (dashed curves) and map-to-truth FSC (solid curves), together with local refinements initialized from 10°-perturbed poses (dash-dot) and the oracle reconstruction from true poses (dotted, representing the information ceiling of the 2D micrographs). The End2EndPtychoSPA run from the perturbed poses shows the end of the full schedule (16 epochs); the End2EndPtychoSPA run from the failed 2D route shows stage 1 (8 epochs). (e) ResLog scaling ($1/d^2$ versus $\ln N$, where $d$ is the half-map FSC 0.143 crossing and open symbols denote the map-to-truth FSC 0.5 crossing); no linear fit is shown for the 2D route with global orientation search because global pose alignment failed. (f) Angular orientation error distributions for the complete pipelines compared against uniform random orientations in SO(3) (dashed curve). (g, h) Angular orientation and in-plane translation error distributions following local refinement from ground-truth poses perturbed by 10° rotation and 2 Å translation (dashed curves: initial perturbed poses; End2EndPtychoSPA after the full 16 epochs). (i) Single-particle alignability, defined as the fraction of particles whose true orientation achieves the highest cross-correlation among 2,000 random orientations against the ground-truth map, evaluated for myoglobin (Mb, 30 and 45 e/Å²) and haemoglobin (Hb, 30 e/Å²). Bars: map low-pass filtered to 8 Å; black ticks: low-pass filtered to 5 Å. The 2D ptychographic micrographs are reconstructed without a prior, with a plug-and-play DRUNet prior (not run on Hb), and with a deep image prior embedded inside ptychography (pty) or applied as a post-filter (post).

experimental apoferritin data), which is the largest defocus magnitude at which at most 1% of the incident probe intensity falls outside the 230.4 Å probe window; all higher-order aberrations were set to zero. Each micrograph is a 21 × 21 raster scan with a step size of 30.7 Å (85% linear probe overlap) across a 614.4 × 614.4 Å scan field at an incident fluence of 30 e/Å² (28,312 electrons per diffraction pattern). Multislice wave propagation (Cowley & Moodie, 1957) was performed on a real-space grid of 0.3 Å pixels with 2 Å slice thickness within a 691.2 × 691.2 Å window around each scan position, band-limited to $\frac{2}{3}$ of the Nyquist frequency. Amplitude contrast from inelastic scattering was included via an absorptive potential parameterized from the inelastic cross-sections of Langmore & Smith (1992) following Vulović *et al.* (2013) without empirical rescaling. The far-field exit wave intensity was integrated over each detector pixel without interpolation. Because the experimental datasets were recorded without an energy filter, the absorbed electron fraction was added back to the detector as an incoherent inelastic scattering channel (modelled at a single mean energy loss of 25 eV) before drawing Poisson shot noise for each pattern across all 50 simulated micrographs.

#### S10.3. Baseline 2D reconstruction route on simulated data

We estimated the defocus $C_{10}$ of each micrograph via a windowed single-sideband defocus sweep (Rodenburg *et al.*, 1993; Pennycook *et al.*, 2015) and reconstructed a Wiener-filtered SSB phase image (You *et al.*, 2026) for automated particle picking. Template matching used 300 projections of the heavy-atom coordinates of the model used to construct the simulated protein (2DN2 for haemoglobin and 1A6M for myoglobin), with the template-resolution parameter set to 20 Å. Otsu thresholds were determined from the template-matching scores. Thus, picking used a structural template, while RELION's primary initial volume was generated **de novo**.

For haemoglobin, the pooled picking score used a 10 Å matching tolerance and excluded a 35 Å border from both candidate picks and true particle centres; this evaluation region contained 1,287 true centres across 50 micrographs. All 1,409 candidates above the Otsu threshold, including those outside this scoring region, were retained for extraction. One-to-one matching against all true centres yielded 1,374 matches within 10 Å. The separate count of 1,322 fully contained particles requires every projected heavy atom of a particle to lie inside the scanned field (Table S6).

We reconstructed a single-slice iterative 2D ptychographic phase micrograph for each scan (400 epochs, pure-phase object constraint), extracted particles into 100 × 100 pixel boxes at a pixel size of 1.2 Å, and refined them in RELION 5.0.1 (Scheres, 2012) in C1 without CTF correction. Gold-standard half sets were split by micrograph parity, and the primary refinement started from a **de novo** VDAM initial model (Kimanius *et al.*, 2021). A parallel control refinement started from the ground-truth atomic model low-pass filtered to 20 Å.

#### S10.4. End2EndPtychoSPA configuration and vitreous-ice background model on haemoglobin

On the simulated haemoglobin dataset, End2EndPtychoSPA was initialized from the **de novo** RELION half maps and poses and followed the apoferritin reconstruction protocol, with parameters set from the optical geometry and the initial maps without using the ground truth. Both the Poisson data loss and the probe intensity normalization were restricted to the bright-field pupil disk of radius 67 pixels ($1.05\alpha$), excluding the dark-field amorphous-ice scattering ring. The initial volume $V^0$ was scaled by a factor of 4.36, determined by minimizing the initial forward-model loss across candidate scale factors between 0.5 and 16. Because our target resolution lies near 3 Å rather than the 6–7 Å range of apoferritin, we reduced the Gaussian gradient blur from 4 Å to 1 Å and set the spherical support radius to 35 Å with an 8 Å raised-cosine edge, matching the radius (34.9 Å) at which the initial map density falls to two standard deviations above its mean. Each reconstruction item is a 9 × 9 window of scan positions centred on one particle, discretized at a voxel size of 0.72 Å on a 320 × 320 pixel model grid. Refinement ran across three stages of 8, 8 and 16 epochs, with stage 3 using an object step size of 20 (selected over 10 via the unmasked half-map FSC of 3.48 Å versus 3.58 Å).

Because the shared particle volume $V$ represents isolated macromolecules in vacuum, it does not account for the phase shift of the surrounding vitreous ice within the 230.4 Å probe footprint. To model the solvent background, we first supplied each micrograph across all stages with a static background phase map built by inpainting the picked particle regions in its 2D SSB phase image and scaling the result by 0.8. This scale factor was calibrated by minimizing the initial forward-model loss over candidate scales from 0 to 4 on a subset of about 10% of the particles in half 1 (31 items). Although these 31 items were excluded from the training table during the scale scan, they are not independent of the training set: every one of their 9 × 9 scan windows shares diffraction patterns with neighbouring training items (91% of their frames are shared), they come from the same micrographs, and all production refinements train on the full particle set including these items. The factor of 0.8 is therefore an internal calibration on retained data rather than validation on an independent holdout. In addition to this static inpainted background, we added a learnable two-dimensional phase layer per micrograph on the model grid that carries over across stages and is masked to zero inside the 35 Å support radius of every picked particle so that it cannot absorb macromolecular density. The Adam learning rate for this learnable ice layer ($3 \times 10^{-3}$ versus $10^{-3}$, giving half-map FSC 0.143 crossings of 3.22 Å versus 3.27 Å) and its use across all three stages rather than stage 3 alone (3.22 Å versus 3.30 Å) were chosen using the unmasked half-map FSC of the production runs.

Reconstruction repeats and evaluation-grid controls are reported in Section S8.

#### S10.5. Ground-truth evaluation metrics

We generated the ground-truth reference potential on the three-dimensional Fourier grid from the atomic coordinates using the relativistic electron scattering factors of the multislice simulation. Each reconstruction, evaluated as the unfiltered mean of its two half maps, was aligned to the ground-truth potential across both enantiomorphic hands with a final rigid-body refinement at 5 Å resolution. Both reconstruction workflows were evaluated against the ground truth under two masks: a tight molecular mask (built by low-pass filtering the ground-truth potential to 15 Å, thresholding at the molecular volume for 1.21 Å³/Da, dilating by three voxels and adding a 6 Å raised-cosine edge) and a soft spherical mask of 42 Å radius with a 6 Å edge. In our primary table, each reconstruction is evaluated on its native voxel grid: RELION on its 1.2 Å grid (Nyquist limit 2.4 Å) and End2EndPtychoSPA on its 0.72 Å grid, onto which the ground-truth map and masks are resampled from the 1.2 Å grid. Because the two grids have different voxel spacings and Fourier shell widths, and because any FSC crossing finer than 2.4 Å on the 0.72 Å grid lies beyond the Nyquist frequency of the 1.2 Å ground-truth map, we also evaluated both reconstructions on a common grid: first on the 1.2 Å ground-truth grid (after band-limiting End2EndPtychoSPA to $\frac{1}{2.4}$ Å$^{-1}$ and downsampling) and second on the 0.72 Å grid (after upsampling the RELION map), using identical interpolation and shell binning. Any FSC curve that stays above the threshold up to the highest Fourier shell is reported as censored at that shell's Nyquist limit. We measure orientation error as the geodesic rotation angle between the refined and ground-truth particle poses modulo the C2 symmetry axis of haemoglobin; any particle whose orientation error exceeds 90° under C2 symmetry but is within 30° under the pseudo-$D_2$ symmetry relating the $\alpha$ and $\beta$ subunits is counted as a pseudo-dyad flip.

### S11. Computational cost

The two-stage End2EndPtychoSPA refinement of apoferritin requires 10.9 GPU-hours across two NVIDIA H100 GPUs, including 0.31 GPU-hours to construct the initial half volumes

**Table S6**

Particle counts and evaluation subsets for the haemoglobin comparisons in Figure 4. Counts are total (half 1 / half 2); half sets follow micrograph parity. The 1,322 fully contained particles have all projected heavy atoms inside the scanned field. Extraction matching uses the full pick list and all true centres, including particles extending beyond the field boundary, with a one-to-one matching tolerance of 10 Å. This differs from the boundary-restricted population used to score picking (Section S10). Only 688 particles can centre a complete 9 × 9 scan window; the other 721 still contribute as neighbouring copies. All 1,409 particle poses are refined, and pose comparisons use the same 1,374 matched particles. The scale-calibration subset is an internal calibration set with overlapping diffraction frames, and all production runs include these particles.

| Step | Rule | Particles or items (half 1 / half 2) | Excluded from this step | Used for |
|---|---|---|---|---|
| Simulated particles | 70 per micrograph, 50 micrographs | 3,500 (1,322 fully inside the scanned field) | – | ground truth |
| Picked and extracted | template matching with Otsu threshold; 100 × 100 pixel box at 1.2 Å | 1,409 (702 / 707) | 0 boxes | RELION refinement and its half maps |
| Matched to a true particle | one-to-one assignment of a pick to a true centre within 10 Å | 1,374 (685 / 689) | 35 from pose evaluation | pose errors of both routes |
| End2EndPtychoSPA particle table | same particles, halves and start poses as RELION | 1,409 (702 / 707) | 0 | particle poses refined as centres or copies |
| Reconstruction items | 9 × 9 scan window inside the 21 × 21 scan | 688 (345 / 343) | 721 (357 / 364) as item centres | End2EndPtychoSPA half maps |
| Particle copies per item | same half; centre inside the item field plus 35 Å | median 18 (8–30) | – | every one of the 1,409 particles is in at least one item |
| Scale-calibration subset | about 10 % of each half, removed from the table for the scale scan only | 141 (70 / 71); 67 items (31 / 36) | – | ice-background scale (31 items of half 1) |

(Table S7). The upstream 2D-route processing on which this initialization depends requires an additional 6.2 GPU-hours covering the single-sideband fits, the 89 iterative 2D ptychographic reconstructions and the three sequential RELION refinements, bringing the complete end-to-end dependency path to 17.1 GPU-hours. Computational times for our initial cryoSPARC reproduction and descan plane fits were not logged, and computational costs were not reported in the original publications for the deposited benchmarks.


**Table S7**

Computational cost of the experimental reconstructions. Top: breakdown of the apoferritin End2EndPtychoSPA refinement and its upstream 2D-route initialization. Bottom: computational cost of the remaining experimental reconstruction runs. GPU-h counts each GPU separately; the Phi92 run occupied one 4-GPU node, so 1.52 node-h equal 6.08 GPU-h. CPU post-processing and evaluation steps are not included.

| Stage | Machine | GPU-h |
|---|---|---|
| apoferritin End2EndPtychoSPA start halves | 2 × H100 | 0.31 |
| apoferritin End2EndPtychoSPA stage 1 + stage 2 | 2 × H100 | 10.61 |
| single-sideband fits (2D route) | H100; one scan on a local A6000 | 3.30 |
| 89 ptychographic reconstructions (2D route) | H100 | 2.08 |
| RELION: seeded, global, R1 (2D route) | local A6000 | 0.78 |
| **apoferritin dependency path, total** | | **17.08** |
| apoferritin ResLog End2EndPtychoSPA, $\frac{N}{8}$, $\frac{N}{4}$, $\frac{N}{2}$ | 2 × H100 | 5.61 |
| apoferritin ResLog End2EndPtychoSPA, second $\frac{N}{8}$ draw | 2 × H100 | 0.78 |
| T20S (3 stages) | 2 × H100 | 9.12 |
| TMV | 2 × H100 | 7.23 |
| Phi92 | 4 × GH200 (one node) | 6.08 (1.52 node-h) |

Rodenburg, J., McCallum, B. & Nellist, P. (1993). *Ultramicroscopy* **48**, 304–314.
Scheres, S. H. (2012). *Journal of Structural Biology* **180**, 519–530.
Vojtěchovský, J., Chu, K., Berendzen, J., Sweet, R. M. & Schlichting, I. (1999). *Biophysical Journal* **77**, 2153–2174.
Vulović, M., Ravelli, R. B., Vliet, L. J. van, Koster, A. J., Lazić, I., Lücken, U., Rullgård, H., Öktem, O. & Rieger, B. (2013). *Journal of Structural Biology* **183**, 19–32.
You, S., Romanov, A. & Pelz, P. M. (2025). *Physica Scripta* **100**, 015404.
You, S., Varnavides, G., Khavnekar, S., Palatkin, N., Shao, S., Wu, M., Stroppa, D., Chernikova, D., Zhu, B., Egoavil, R., Vespucci, S., Krishnan, D., Ye, X., Schur, F. K. M., Spiecker, E. & Pelz, P. (2026). *Advanced Science* **13**, e76620.
